# The Brain Time Toolbox, a software library to retune electrophysiology data to brain dynamics

**DOI:** 10.1101/2021.06.09.447763

**Authors:** Sander van Bree, María Melcón, Luca D. Kolibius, Casper Kerrén, Maria Wimber, Simon Hanslmayr

## Abstract

Human thought is highly flexible, achieved by evolving patterns of brain activity across groups of cells. Neuroscience aims to understand cognition in the brain by analysing these intricate patterns. We argue this goal is impeded by the time format of our data – clock time. The brain is a system with its own dynamics and regime of time, with no intrinsic concern for the human-invented second. Here, we present the Brain Time Toolbox, a software library that retunes electrophysiology data in line with oscillations that orchestrate neural patterns of cognition. These oscillations continually slow down, speed up, and undergo abrupt changes, introducing a disharmony between the brain’s internal regime and clock time. The toolbox overcomes this disharmony by warping the data to the dynamics of coordinating oscillations, setting oscillatory cycles as the data’s new time axis. This enables the study of neural patterns as they unfold in the brain, aiding neuroscientific inquiry into dynamic cognition. In support of this, we demonstrate that the toolbox can reveal results that are absent in a default clock time format.

## Studying dynamic cognition

Everyday tasks involve a plethora of cognitive functions that operate dynamically in tandem. Something as mundane as taking notes during a meeting or battling your friend in a video game requires attention, motor activity, perception, memory, and decision-making, each evolving over time. How does the brain achieve dynamic cognition? To answer this question, neuroscientists closely study how brain activity unfolds from one moment to the next using temporally precise neuroimaging methods. These include electroencephalography (EEG), magnetoencephalography (MEG), and single and multiunit recordings – grouped together under the term electrophysiology.

## Seconds are foreign to the brain

In a typical electrophysiology study, neuroscientists first probe cognition by introducing an experimental manipulation. For example, an attention researcher might introduce a set of moving dots. Then, to understand cognition in the brain, they perform a series of analyses on the recorded data. They might study changes in scalp topography over a second of data, apply machine learning to characterize how the representation of the dots evolves, or perform any other time-dependent analysis.

Critically, from the raw output of neuroimaging devices to the analysis of recorded brain signals, time is operationalized as clock time – sequences of milliseconds. We claim that clock time, with all its benefits for human affairs, is generally inappropriate for neuroscience. This is because clock time is defined by us and for us, based on how long it takes for Earth to rotate its axis. The brain itself, however, employs its own regime of time, dictated by its own dynamics. As such, the brain is indifferent to how many milliseconds, seconds, minutes, or hours have passed unless it is expressly relevant for specific behaviour, such as maintaining circadian rhythms [1] or tracking a time-dependent reward [2]. Instead, the brain is concerned with coordinating communication between cells in a delicate time-sensitive manner, such as sending information at one moment and receiving feedback signals at the next. Hence, the brain’s intrinsic time format – brain time – is dictated by the internal processes that clock brain activity (for an explanation of key terms, see the Glossary in Table 1).

**Table 1:**
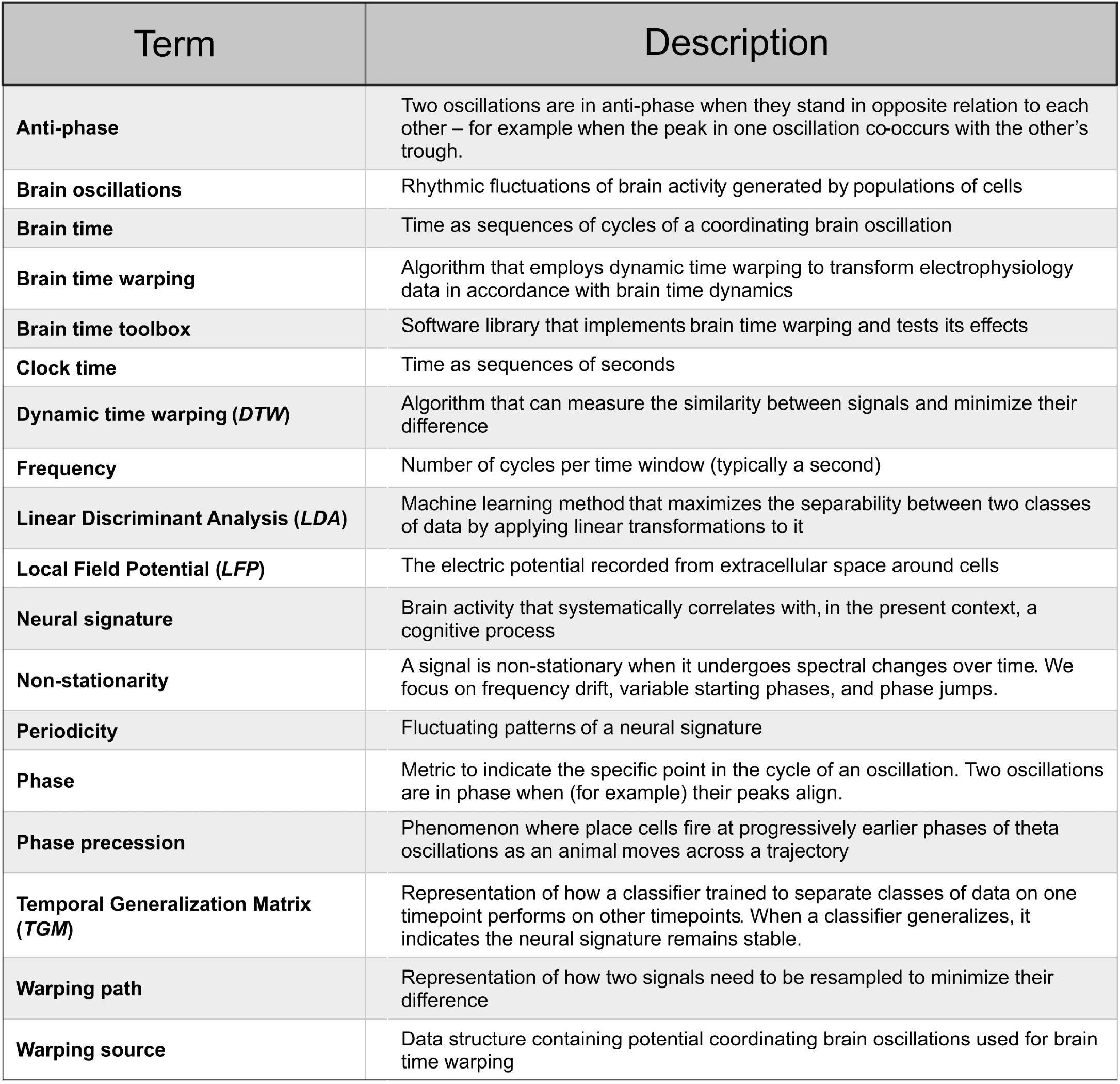
Glossary.

| Term | Description |
| --- | --- |
| **Anti-phase** | Two oscillations are in anti-phase when they stand in opposite relation to each other – for example when the peak in one oscillation co-occurs with the other's trough. |
| **Brain oscillations** | Rhythmic fluctuations of brain activity generated by populations of cells |
| **Brain time** | Time as sequences of cycles of a coordinating brain oscillation |
| **Brain time warping** | Algorithm that employs dynamic time warping to transform electrophysiology data in accordance with brain time dynamics |
| **Brain time toolbox** | Software library that implements brain time warping and tests its effects |
| **Clock time** | Time as sequences of seconds |
| **Dynamic time warping (DTW)** | Algorithm that can measure the similarity between signals and minimize their difference |
| **Frequency** | Number of cycles per time window (typically a second) |
| **Linear Discriminant Analysis (LDA)** | Machine learning method that maximizes the separability between two classes of data by applying linear transformations to it |
| **Local Field Potential (LFP)** | The electric potential recorded from extracellular space around cells |
| **Neural signature** | Brain activity that systematically correlates with, in the present context, a cognitive process |
| **Non-stationarity** | A signal is non-stationary when it undergoes spectral changes over time. We focus on frequency drift, variable starting phases, and phase jumps. |
| **Periodicity** | Fluctuating patterns of a neural signature |
| **Phase** | Metric to indicate the specific point in the cycle of an oscillation. Two oscillations are in phase when (for example) their peaks align. |
| **Phase precession** | Phenomenon where place cells fire at progressively earlier phases of theta oscillations as an animal moves across a trajectory |
| **Temporal Generalization Matrix (TGM)** | Representation of how a classifier trained to separate classes of data on one timepoint performs on other timepoints. When a classifier generalizes, it indicates the neural signature remains stable. |
| **Warping path** | Representation of how two signals need to be resampled to minimize their difference |
| **Warping source** | Data structure containing potential coordinating brain oscillations used for brain time warping |

## Cycles as the brain’s native unit

How is brain activity organized? A defining feature of the brain is that its activity waxes and wanes [3], pointing to a central role of brain oscillations. Brain oscillations are well-geared to structure brain activity. For one, each cycle of an oscillation contains a window of excitability where cells are more likely to fire [4,5]. Moreover, oscillations vary in their frequency, meaning the excitability windows vary in duration. The functional role of oscillations has been shown across a wide array of cognitive functions, including attention [6,7,8], perception (auditory [9,10], visual [11,12], tactile [13,14]), action [15,16], memory [17,18,19], and decision-making [20,21]. Together, this situates brain oscillations as the brain’s clocking mechanism, clustering brain activity in flexible ways to organize dynamic cognition. The brain’s base unit of time then, are the cycles of oscillations that coordinate neural firing, not the milliseconds with which we format our data.

## Clock and brain time are usually out of tune

Why does it matter that we use a foreign time format? When neuroscientists study dynamic cognition, they repeat measurements across trials, resetting their stopwatch at the start of each. However, some oscillations do not reset [22,23]. Even when most do, oscillations evolve continuously in frequency and show jumps in phase (Figure 1). Thus, besides a potential mismatch between clock and brain time from the get-go caused by variable starting phases, the disharmony between the dimensions accumulates due to frequency drift and phase jumps. These are prime examples of eccentricities in brain dynamics, formally called non-stationarities (and there are more [24]). Their presence makes clock time an ill-suited format to study temporal patterns of dynamic cognitive function – it distorts how the brain itself carries information forward in time.

**Figure 1.**
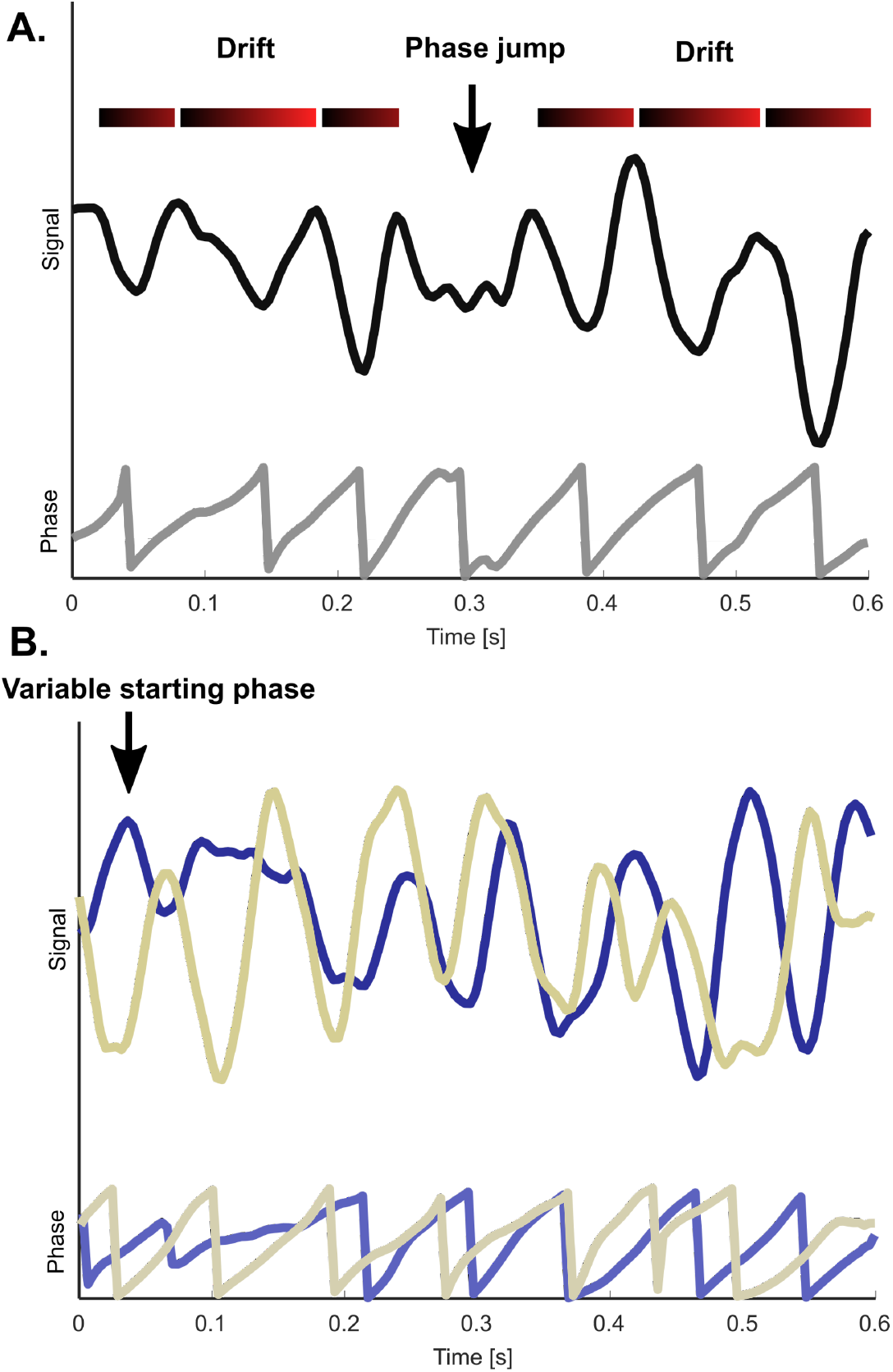
Sources of disharmony between clock and brain time. Brain oscillations are non-stationary, which causes a noncorrespondence or “disharmony” between the brain’s internal dynamics and clock time. **(A)** An oscillation shows frequency drift and a spontaneous jump in phase, resetting itself. **(B)** Two oscillations with different starting phases. The blue oscillation starts with a rising phase, while the sand-coloured oscillation starts with a falling phase. The top rows of each panel show the amplitude fluctuations of oscillations, while the bottom rows show the phase.

To demonstrate this point, take again the case of spatial attention. Studies show that alpha oscillations (8 to 12 Hertz; Hz) in parietal regions orchestrate the dynamics of spatial attention [25,8]. If these oscillations vary in their starting phase across trials, then the neural patterns of spatial attention will vary along with it. Likewise, if the oscillations slow down in frequency, the patterns slow down too. If a researcher is interested in, say, decoding the locus of covert attention in the visual field over time, it would muddy the waters to do so in clock time, ticking away with its equal periods. The slowing down of alpha oscillations means brain time falls behind relative to clock time, so analysing the data in its default format yields a sped-up readout of attention’s true pattern. Instead, we argue the dynamics of clocking oscillations should heavily inform data analysis. As a general mantra, the optimal approach to analyse the brain, like any other system, is with recourse to its own dynamics – from inside out [26,27] (Figure 2).

**Figure 2:**
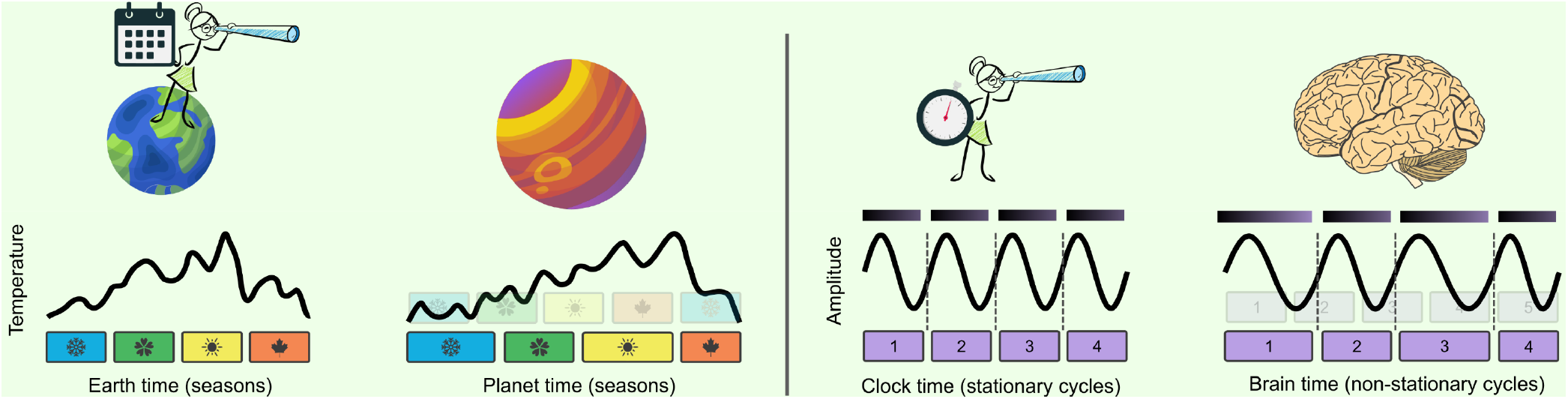
What is the best way to study a foreign system? *(Left)* Imagine a planet with seasonal dynamics radically different from Earth’s, where the duration of each season differs substantially (as a real-world example, planet Kepler-413b has erratic seasons due to its eccentric orbit [28]). To understand the system, we measure a variable of interest across time, such as surface temperature. Critically, how do we define time here? If we plot temperature as a function of Earth’s seasons (Earth time), the data will be heavily distorted, hampering interpretation. Instead, if we were to study the system with recourse to its own dynamics (system time), the same temporal patterns in the data become interpretable. *(Right)* As neuroscientists, we are in an analogous position – we are studying a foreign system with its own dynamics. So, in the same vein, we should interpret data patterns with reference to the brain’s dynamics, enabling an accurate readout of evolving patterns of information.

The problem of disharmony does not end here. Neuroscientists repeat measurements across participants to establish whether effects found in the data are representative and statistically robust. But different brains have different dynamics, resulting in disharmony *across* brains too. In the attention experiment, it is highly relevant that the clocking alpha oscillations differ in frequency from person to person [29] as it means the patterns differ too. Looking for evolving patterns of spatial attention by averaging across participants is like asking when spring turns summer in a solar system that contains diverse planets – it only makes sense after correcting for individual dynamics (Figure 2).

## Approaches to factor in brain time

Disharmony between clock and brain time impedes scientific analysis within and across brains. To overcome this problem, approaches have been developed that factor in brain dynamics. At minimum, the phase of oscillations can be tracked to explain some of the variance in brain data [19] or behaviour [30,31], and analyses can be locked selectively to oscillatory peaks or troughs [32]. Then there are more expansive techniques, where the electrophysiological data is restructured before any analysis is carried out. These include organizing the data with phase as the time axis [33], as well as linear time warping approaches that transform a template signal based on trial-by-trial variations in brain dynamics [34].

Such approaches can reveal brain patterns of interest that are otherwise obstructed by clock time’s distorting effects. For example, the phenomenon of phase precession has been extended from rodents to humans when using phase as the time axis, with no such effect visible in clock time [33]. As another example, linear time warping uncovers oscillatory brain patterns on the trial average by correcting for differences in brain time across trials [34].

While these approaches come a long way in factoring in brain dynamics, each is limited in their scope. Using phase to explain variance in the data or locking analyses to peaks or troughs provides insights about selective data samples but does little to enable the readout of dynamic patterns. Setting the time axis to phase may not always be possible and departs significantly from the original data structure – such that unique phase-based analyses are needed. Linear time warping can equalize brain time across trials but is less equipped to deal with non-stationarities throughout the trial due to its linear nature.

## A dynamic and holistic way to factor in brain time

Here, we introduce **brain time warping** as a method to account for the disharmony between clock and brain time. This approach overcomes previous limitations in the following way. First, it identifies segments throughout each trial where clock and brain time fall out of tune. Then, it adapts the data to reduce their difference – winding back clock to brain time sample by sample (Figure 3). Brain time warping incorporates an algorithm called dynamic time warping (DTW), which characterizes the similarity of two signals [35,36]. DTW computes a warping path, which shows how the samples of each signal need to be transformed to optimize their alignment. For brain time warping, those signals are clock and brain time (Figure 3A).

**Figure 3:**
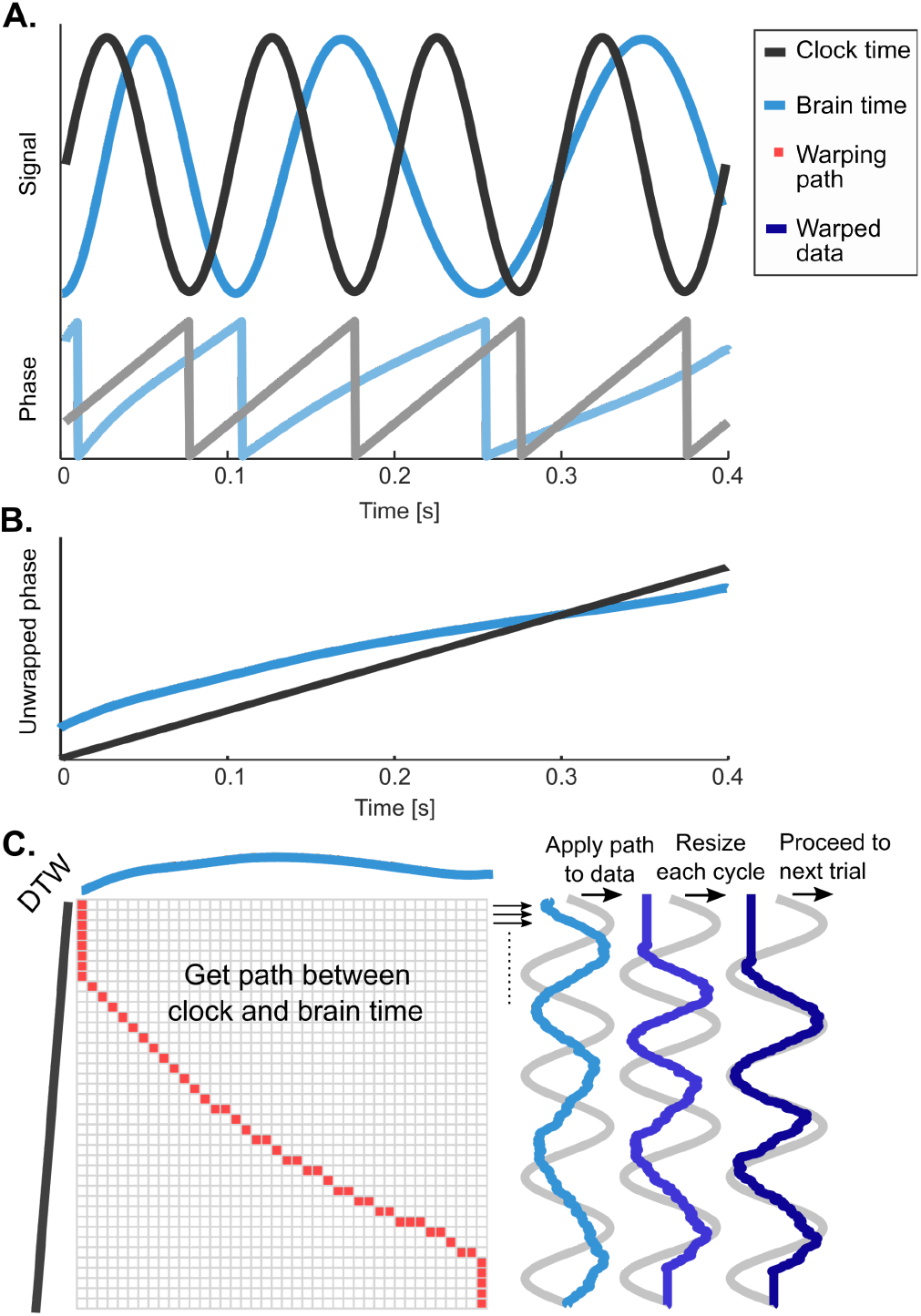
Brain time warping between clock and brain time. **(A)** Brain time starts in its rising phase and slows down its frequency over the course of the trial, both causing a mismatch to clock time (defined as a stationary signal fluctuating in sync with a researcher’s stopwatch). **(B)** To facilitate warping, the phase of clock and brain time are unwrapped, meaning phase is computed without cycle resets. **(C)** DTW calculates a warping path that minimizes the difference between the dimensions. Cycle by cycle, the path is applied to the input electrophysiology data, transforming its dynamics in accordance with the brain’s dynamics. To enable alignment of brain time across trials, the data of each cycle is resized to a constant number of samples. The previous steps are repeated for all remaining trials. Upon completion, the data’s time axis is changed from seconds to cycles of brain time. The data is no longer in clock time, but in brain time.

How are clock and brain time operationalized? Brain time can be characterized as the phase of the oscillations hypothesized to orchestrate a process’ dynamics, with its variable starts, drift, and phase jumps. Clock time can be characterized as the phase of a stationary sine wave, fluctuating away faithfully to seconds. Such a signal is what brain time would look like without the three sources of disharmony; here milliseconds and cycles map directly onto each other. (For example, in Figure 3A, multiples of 100 milliseconds correspond to each zero-crossing of the clock time signal.) DTW highlights during which samples clock and brain time fall into disharmony, and this warping path can be used to transform the original data. Concretely, at samples where the warping path suggests brain time needs to repeat itself before ramping back up to clock time, brain time warping repeats samples in the original data (Figure 3C). Looping back to our attention example, at segments where DTW indicates alpha oscillations slow down, brain time warping stretches the data by repeating samples in an attempt to bring its structure closer to the true dynamics of spatial attention.

Brain time warping loops over trials, continuously correcting disharmony by applying the warping path cycle by cycle. In effect, brain time warping adapts data to the dynamics of an oscillation of interest in a data-driven way. The result is a dataset in brain time rather than clock time, and as such, the time axis has changed from seconds to cycles. As the data is referenced to individual dynamics, it also becomes easier to look for temporal patterns across brains. Here, we introduce the Brain Time Toolbox (https://github.com/sandervanbree/braintime, Supplementary Methods), a toolbox built for MATLAB (The MathWorks, Natick, Massachusetts, USA) that implements brain time warping and tests its effect. This software library was built for electrophysiology data analysis, including EEG, MEG, and single and multi-unit recordings. The toolbox lets users select a brain time signal from one or more warping sources (e.g., channels or independent components extracted from a data structure; Supplementary Methods, Toolbox, section 2.2.1). This signal then serves as the basis for brain time warping as laid out in Figure 3. In the upcoming section, we methodologically validate brain time warping – utilizing the toolbox throughout.

## Results

We tested the algorithm’s effects across three electrophysiology datasets. In the first dataset (N = 10 participants), we simulated a basic attentional spotlight model, giving us full control over the ground truth brain patterns (Figure 4A). Briefly, we placed one conducting dipole in the right parietal cortex, which exerted top-down control over the phase of two follower dipoles, one located in each visual cortex. We generated two conditions, one where attention was oriented to the left hemifield, and one to the right. These conditions systematically differed in their underlying brain activity. Specifically, the conducting dipole always forced its contralateral follower to the same phase while setting an anti-phase relation with its ipsilateral follower (drawing upon experimental findings [37,38,39]). In addition, the conducting dipole inhibited the contralateral follower dipole, reducing its amplitude [40]. We then injected the data with variable starting phases and frequency drift to mimic natural brain activity.

**Figure 4:**
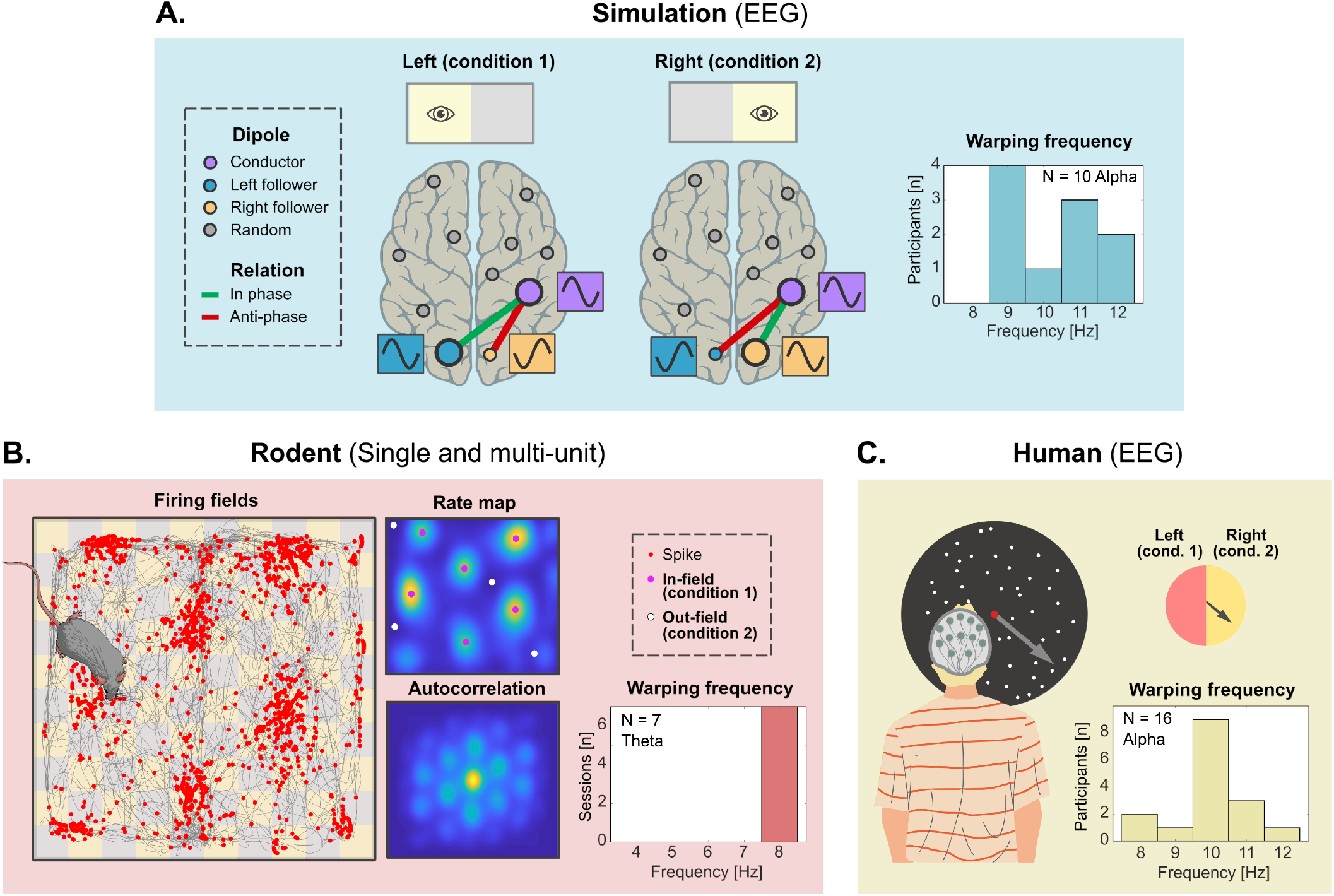
Electrophysiology datasets used to validate brain time warping. **(A)** Simulated electroencephalography (EEG) data (N = 10 virtual participants). We developed a basic attentional spotlight model with dipoles oscillating at alpha frequencies (8 to 12 Hertz [Hz]). One parietal conductor dipole controlled two follower dipoles, one in each visual cortex. Two conditions differed in brain activity due to the phase relation across dipoles and the suppressed amplitude of the contralateral follower dipole, together yielding attention’s neural signature. To introduce disharmony between clock and brain time, we added frequency drift and variable starting phases to the data. We brain time warped to alpha oscillations in visual regions (right). **(B)** Rodent single and multi-unit data (N = 7 sessions). Long-Evans rats navigated through a square field while single units and local field potentials (LFP) were recorded from the entorhinal cortex. We identified grid cells and cut the data depending on whether the animal was traveling through a location coded by a grid cell (condition 1) or not (condition 2). We display cell spike locations in the field, its smoothed representation (rate map), and this representation’s autocorrelation. We then brain time warped grid cell firing patterns to theta oscillations (4 to 8 Hz) measured in LFP recordings (bottom right). **(C)** Human EEG data (N = 16 participants). Participants viewed random dot kinematograms, with dots moving at one of two levels of coherence (25.6% or 51.2%) in a direction ranging from 1° to 360°. We pooled the levels of coherence and binarized direction toward left-(condition 1) or rightward (condition 2) motion. We then brain time warped the EEG data to parietal alpha oscillations (bottom right).

In a second dataset (N = 7 sessions), we warped intracranial data from rodents navigating through a square field (Figure 4B; data obtained from [41]). Here, we were interested in characterizing the dynamics of the local field potential (LFP) and grid cell firing patterns in the entorhinal cortex. Just like in the simulation, we again formatted the data into two conditions. Specifically, the data was split based on whether the animal was travelling through a field coded by the grid cell or through another location not coded by the grid cell.

In a third dataset (N = 16 participants), we warped EEG data recorded while human participants observed moving dots (Figure 4C; data obtained from [42]). The two conditions were set around the motion direction of dots – with one condition comprising trials with leftward motion, and the other comprising rightward motion. In Supplementary Methods (Datasets, section 1), we report additional details on each dataset.

### Basic analyses

In basic analyses, we asked whether warping recovers oscillatory neural activity for each dataset by comparing clock and brain time on a number of measures. For these analyses, we pooled across conditions – focusing on general effects of brain time warping on electrophysiology data and ignoring cognition for the moment. First, we tested whether warping increases the oscillatory structure of event-related potentials of channels near the predicted location of coordinating brain oscillations. This analysis provides a qualitative indication on the question of whether brain time warping overcomes non-stationarities.

Second, we performed a time frequency analysis across all channels and tested whether warped data reveal a higher peak at the predicted frequency of interest in the power spectrum. This quantifies the degree to which brain time warping is able to overcome frequency drift and phase jumps in the data.

Third, we compared the intertrial coherence (ITC) across all channels between clock and brain time. This measure tests for the consistency of oscillatory phase across trials and thereby additionally taps into the degree to which the algorithm overcomes variable starting phases (which ordinarily reduce ITC by jittering the phase relation across trials). We predicted that brain time warping increases ITC at the predicted frequency of interest by equalizing brain time across trials.

#### Brain time warping recovers oscillatory activity

In the simulated dataset, the event-related average of clock time data shows no robust oscillatory shape. After warping, the oscillatory structure hidden in the data becomes qualitatively uncovered (Figure 5A). Moreover, we found that the algorithm sharpens the power spectrum selectively around the warping frequency (Figure 5B), and, in line with our hypothesis, increases the peak’s magnitude. Third, the ITC in clock time shows comparatively low values around simulated alpha rates – with only weak clustering at low alpha (Figure 5C). ITC is enhanced by brain time warping, revealing a strong cluster at participants’ warping frequency. We find similar results for the rodent dataset (Supplementary Results, section 3.2.1). For the human data, we found a prominent increase in ITC at the warping frequency, but not in the peaks of the power spectrum (Supplementary Results, section 3.3.1). Together, these results demonstrate that warping can reveal oscillatory activity that is lost due to clock time’s distorting effects on the data.

**Figure 5:**
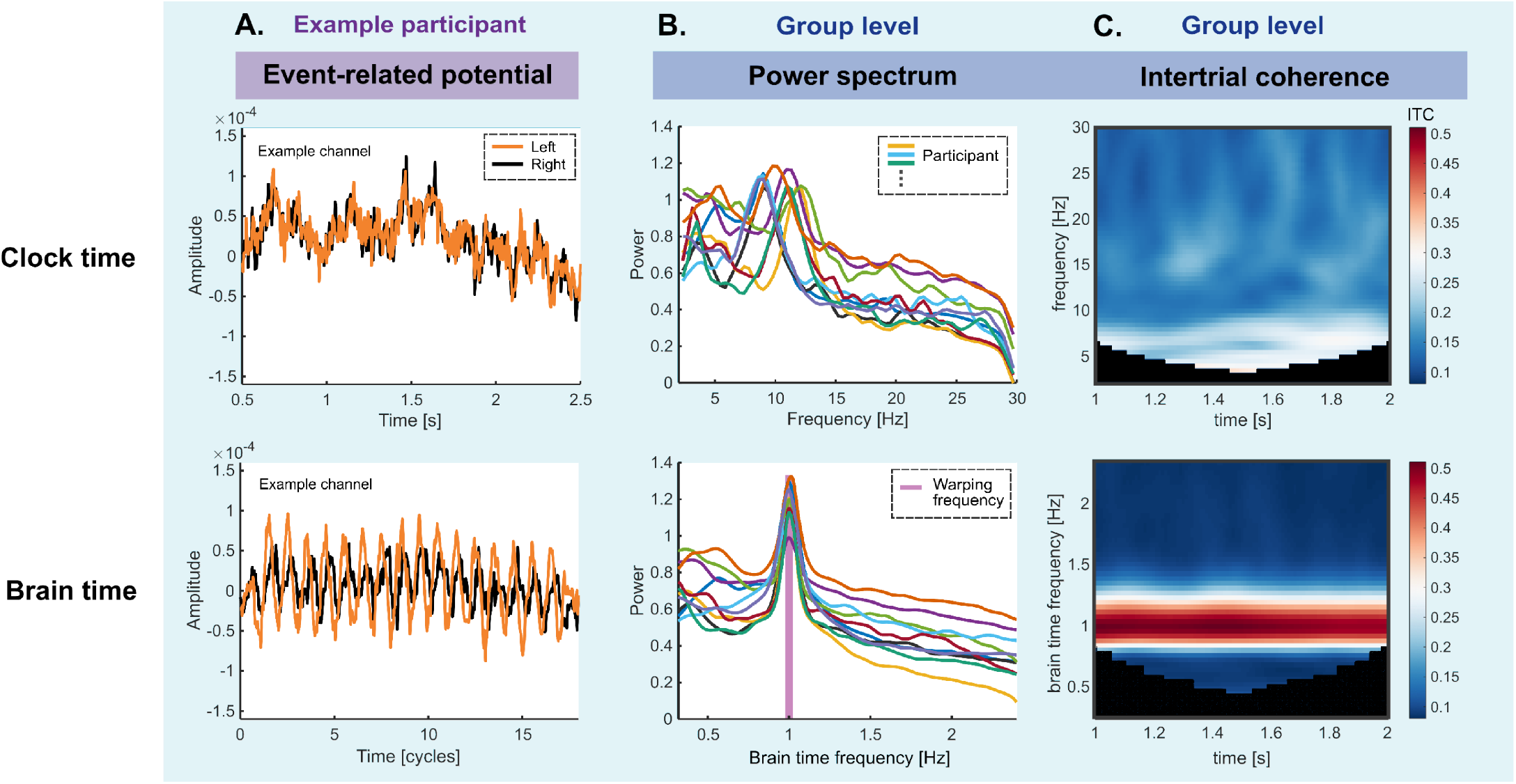
Results of basic analyses in the simulated dataset. **(A)** Event related potentials (ERPs) in the left visual cortex of one example virtual participant. In clock time, averaging across trials destroys the simulated oscillatory structure. By repairing disharmony, brain time warping recovers this structure. **(B)** Power spectra averaged across all channels and participants. Brain time warping increases the power of alpha oscillations at the simulated rate for each participant. Brain time results are always re-referenced to individual participants’ brain time rate 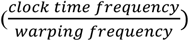. This sets 1 Hz as each participant’s warping frequency (and e.g., 0.5 Hz as half the warping frequency). We removed the aperiodic (1/f) component of the power spectrum using the FOOOF toolbox [47]. **(C)** Intertrial coherence (ITC) averaged across all channels and participants. Brain time warping increases ITC at the (ground truth) alpha frequency. The analysis window was restricted to 1 to 2 seconds to reduce artefacts. We equalized the y-axis in (B) and the colour axis in (C) between clock and brain time based on maximum values to enable a visual comparison between dimensions (this step is performed in all figures).

### Advanced analyses

In advanced analyses, we tested whether brain time warping unveils dynamic patterns of cognition. For each dataset, the conditions were set up such that their contrast was expected to yield a difference in brain activity that makes for a neural signature of cognition. For example, in the human dataset, the difference in neural activity between leftward and rightward motion trials estimates the neural signature of spatial attention because activity unrelated to attending to one or other spatial direction is factored out. If oscillations of interest dynamically clock the neural processes underlying cognition, this should cause the neural signature to vary along with the oscillations.

Pattern classifiers capitalize on neural signatures to make predictions about condition (i.e., class) membership of untrained data. As such, classifier performance makes for a useful index of the fidelity of neural signatures of cognition -with fluctuations of classifier performance indicating fluctuations in the neural signature (periodicity; [43]). Building off these assumptions, we predicted that brain time warping enhances periodicity of these neural signatures by adapting for non-stationarities in the data.

To test this, we used a linear discriminant analysis (LDA) to classify condition in each dataset. We trained the LDA on each timepoint and tested how well it generalized to all other timepoints [44]. This resulted in a two-dimensional temporal generalization matrix (TGM) that provides a robust map of classification performance over time (and so too of the fidelity of the neural signature). In summary, we assumed periodicity in TGMs is a high-level measure to detect periodic brain patterns of cognition, and we predicted that brain time warping would increase such patterns by factoring in the dynamics of clocking oscillations.

#### Quantifying periodicity

For the advanced analyses, we quantified TGM periodicity by applying a Fast Fourier Transform (FFT) over each row and column of TGMs. Then, we averaged the resulting spectra into a single spectrum – the *periodicity spectrum.* This spectrum quantifies how much the neural signature fluctuates at different frequencies. To test our hypothesis that brain time warping increases the oscillatory structure of neural signatures, we contrasted the spectra obtained from clock and brain time data against each other – comparing the periodicity at predicted frequencies of interest.

Moreover, to aid visual inspection of periodicity, we also report autocorrelation maps of TGMs for each dataset. These maps are generated by correlating TGMs with iteratively shifted versions of itself, resulting in a representation of its self-similarity that brings out latent periodic structure [45].

#### Statistically testing periodicity

To evaluate which frequencies in the periodicity spectrum are statistically reliable, we compared the empirically derived spectrum with periodicity spectra obtained under the null hypothesis that there is no fluctuating neural signature of cognition. To do so, we created a pool of TGMs obtained from an LDA trained using randomly permuted classification labels. This procedure destroys the true class structure inherent to the data [46], leaving the classifier with pseudo neural signatures that do not contain generalizable information about the cognitive process under investigation. To establish statistical reliability, we compared the magnitude of peaks in empirically derived TGMs with the distribution of magnitudes in TGMs obtained with permuted labels – both for clock time and brain time data. For more details on basic and advanced analyses, see Supplementary Methods (section 1.1.1 and 1.1.2).

#### Brain time warping recovers neural patterns of cognition

In the simulation, the neural signature provided by the classes reflected attention. The simulated patterns of this signature were not detectable in a default clock time format but did emerge after brain time warping (Figure 6).

**Figure 6:**
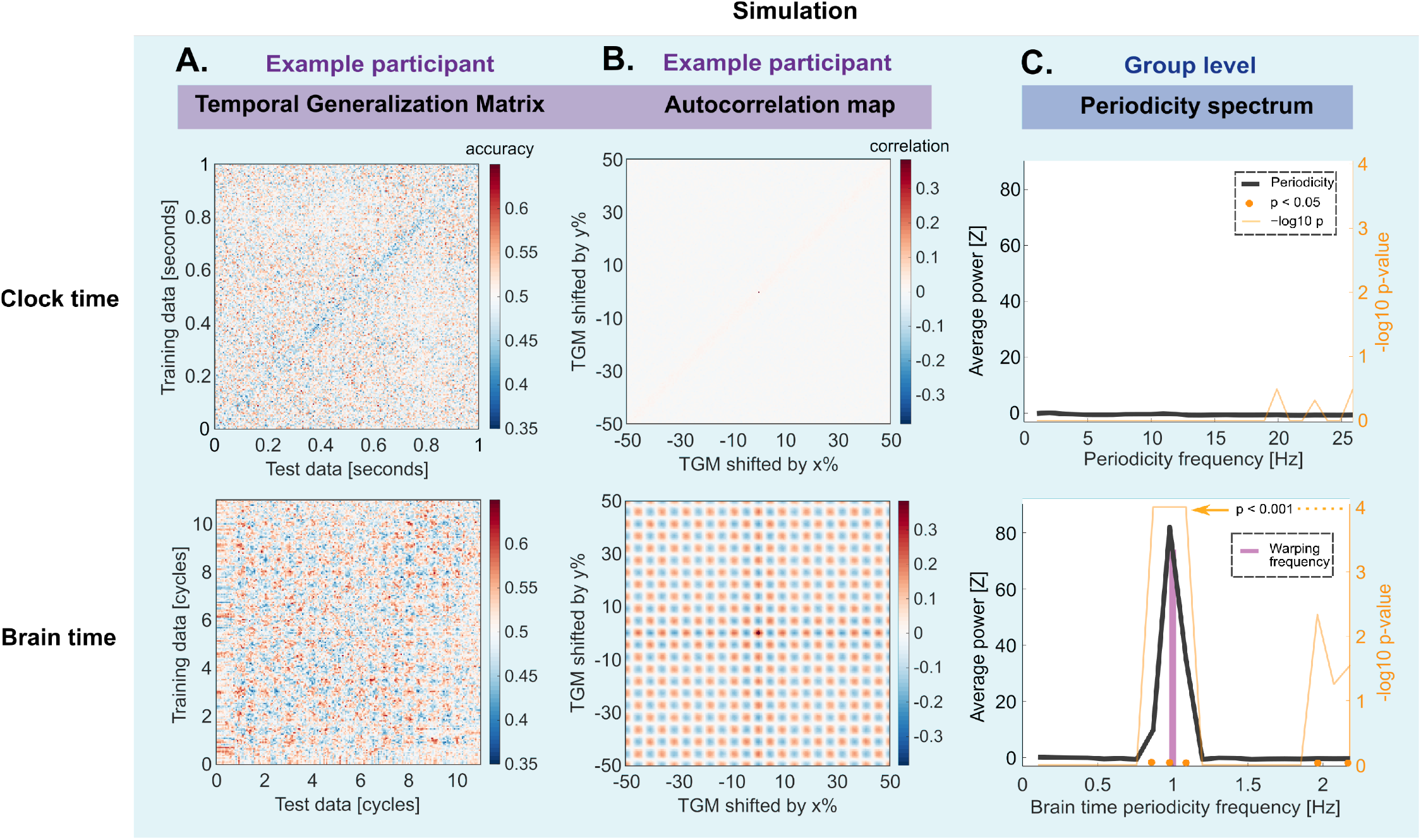
Results of advanced analyses in the simulated dataset. We tested for periodic patterns in the classifier’s temporal generalization matrix (TGM), which provides an index of neural signatures of cognition by demonstrating how the classifier’s performance generalizes across time. **(A)** An example participant’s TGM shows no periodic structure in clock time (top). After brain time warping, the simulated periodic structure in the neural signature is recovered – as evidenced by the checkerboard pattern (bottom). **(B)** The difference between clock and brain time becomes qualitatively striking in the TGM’s autocorrelation maps. **(C)** We quantified periodicity by applying a fast Fourier transform over all rows and columns of TGMs. Then, we perform second-level statistics by comparing empirical periodicity with permuted periodicity (obtained by shuffling class labels). Only brain time spectra show significant periodicity at the warping frequency (p < 0.001) and its first harmonic (p < 0.05), demonstrating brain time warping corrects for disharmony and unveils ground truth neural signature dynamics (bottom and top). Each participant showed periodicity peaks selectively at their warping frequency. We corrected for multiple comparisons (using false discovery rate; FDR [48]), except at specific frequencies in the brain time spectra at which we hypothesized classifier periodicity (0.5 Hz, 1 Hz, 2 Hz; 1 Hz remains significant when applying FDR). In Supplementary Methods, we report the full methods (section 1.2). In Supplementary Results, we provide additional plots, including all TGMs, autocorrelation maps, and periodicity spectra (section 3.1).

In the rodent data, the difference in activity between classes reflected whether the animal was in a place field coded by grid cells, yielding a signature of the neural basis for spatial navigation [49]. In clock time, no periodicity is evident over time (Figure 7A; top) despite grid cells strongly phase locking to theta oscillations ([50]; Supplementary Results, section 3.2.3). Importantly, brain time warping the firing rates of grid cells to theta oscillations obtained from the LFP does result in periodicity (Figure 7A; bottom).

**Figure 7:**
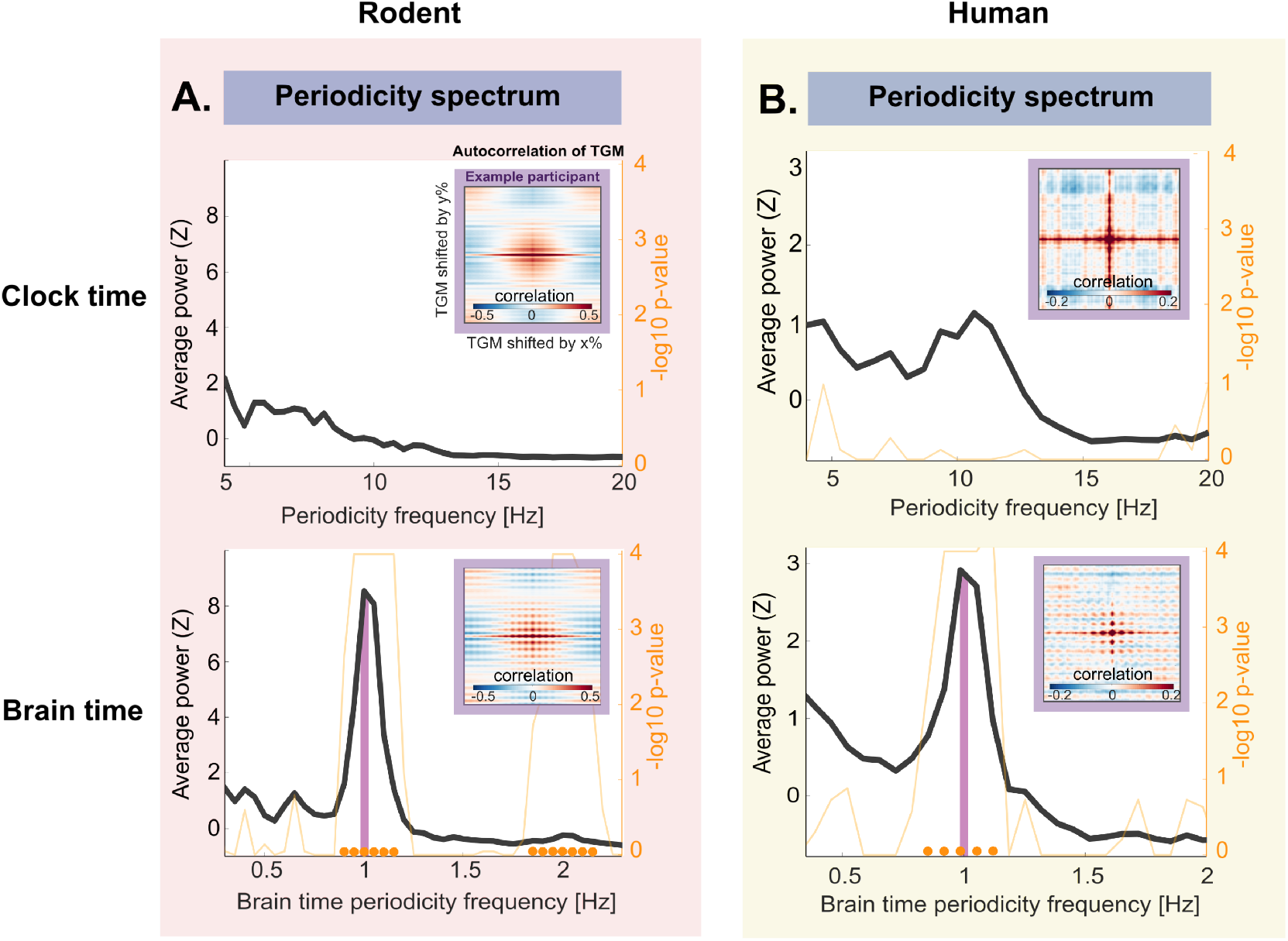
Results of advanced analyses in the rodent and human dataset. **(A)** Warping the rodent data to LFP theta unveils statistically robust periodicity around the warping frequency and its first harmonic (p < 0.001), while the clock time spectrum shows no significant peaks. **(B)** In the human dataset, brain time warping reveals periodic patterns around the warping frequency (p < 0.001), indicating the neural signature of attention fluctuates during motion perception. For the rodent data, each session showed periodicity peaks selectively at their warping frequency. For the human data, not all participants showed periodicity peaks, and there was some variance in the brain time frequency of peaks. The insets display autocorrelation maps of TGMs from example sessions (for the rodent dataset) or example participants (for the human dataset). In Supplementary Methods we report the full methods (section 1.3 and 1.4). In Supplementary Results, we provide additional plots (section 3.2 and 3.3). These plots include TGMs, autocorrelation maps, periodicity spectra, grid cell maps, and spikefield coupling between theta and single unit spikes

In the human data, the neural signature reflected spatial attention, provided by the difference in brain activity to left- and rightward motion. While the clock time data qualitatively shows some periodicity at alpha, the peaks do not stand out reliably from the peaks in periodicity spectra obtained with permuted labels (Figure 7B; top). In contrast, brain time warping reveals prominent and reliable periodicity at participants’ individual alpha frequency (Figure 7B; bottom). Together, the basic and advanced results indicate that brain time warping is a promising tool to repair disharmony between clock and brain time, facilitating the study of the dynamic cognitive brain.

## Discussion

### Is retuning using brain time warping circular?

A potential concern with brain time warping could be that it trivially imposes oscillatory structure onto the data. The worry here is that the algorithm makes data patterns fluctuate at the warped frequency no matter which frequency is selected. In this section we argue when the algorithm is on safe grounds, and when vigilance is needed.

First, as a note of clarification, the path used for brain time warping does not contain oscillatory structure. That is to say, if the path were applied to a random time series, it would not mould it to increase its oscillatory shape. This is because the path is instead designed to show when the oscillation used for warping undergoes non-stationary behaviour. Nevertheless, an important concern remains. Brain time warping applies the path to the data in order to align it with brain dynamics, so if the warping signal is within the data, its stationarity will increase – introducing some oscillatory structure after all. Whether circularity is a problem depends on two factors: the type of analysis carried out, and the dependence between warping signal and warped data. We discuss these points in turn, first covering each type of analysis.

#### Basic analyses

Brain time warping is expected to enhance oscillatory structure in the ERP, power spectrum, and ITC at least somewhat because there are usually at least some oscillations at any frequency to begin with. So, warping to a random frequency will cause those oscillations to have their non-stationarities reduced, affecting subsequent results. Hence, any increase around warping frequencies in basic analyses should not be taken as evidence that those frequencies are critical for cognition – there needs to be a further relation to independent measures (such as cognitive or orthogonal neural variables). In this sense, brain time warping can be used as a pre-processing step to aid subsequent high-level approaches that are sufficiently independent (to avoid circular inference [51]).

With that said, we find that warping to weak simulated oscillations results in only weak frequency-specific enhancements compared to warping to strong oscillations – speaking to the specificity of brain time warping even before cognition enters the scene (Supplementary Methods, section 1.2.2).

#### Advanced analyses

We have presented the advanced classification analysis as a circularity-free high-level approach – and it is the central analysis implemented in the Brain Time Toolbox. By tapping into neural signatures of cognition rather than oscillations themselves, warping-induced changes are nontrivial. Specifically, only warping to those oscillations which coordinate the underlying activity relevant for cognition is expected to yield fluctuations in cognition’s neural signature. In support of this, we performed a control analysis where we warp to control frequencies [52], resulting in no evidence of periodicity (Supplementary Methods, section 1.2.2). A further safeguard of the advanced analysis is that the obtained null distributions benefit equally from any trivially imposed oscillatory structure as the empirical distribution.

#### Data dependence

Beyond the type of analysis that is used, the dependence between warping source and transformed data is another factor to consider. After all, if the cause for circularity lies chiefly in changes to the warping signal, then removing it from the warped data reduces circularity concerns. We report suggestions on how to achieve data independence for each electrophysiology method, describing how they can be implemented before or after warping (Supplementary Methods, Toolbox, section 2.4.1). We also report cases in which independence between warping source and data is impossible. Here, it is important to determine on a case-by-case basis whether circularity is a concern based on the previous points (i.e., are subsequent analyses orthogonal or not?). To aid this process, the toolbox tracks dependence between warping source and warped data and raises a warning when circularity could become an issue.

#### Statistical false positives

With these reflections in mind, we finally emphasize that brain time warping is a hypothesis-driven method that capitalizes on the temporally coordinating nature of oscillations. Hence, we recommend warping only using oscillations predicted to fit that bill to avoid false positive results by chance, or to apply multiple testing correction when re-warping to a different signal in the data. To enable hypothesis-driven warping, the Brain Time Toolbox computes a variety of information about warping signals – including their time frequency characteristics, waveshape, and topographical profile – allowing users to make informed decisions about which signal they wish to designate as brain time.

### When is retuning clock and brain time necessary?

The need to use brain time warping or other methods to retune depends on (1) the degree to which clock and brain time are in disharmony, and (2) the degree to which such disharmony interferes with analyses. Below, we elaborate on both criteria.

The degree to which disharmony is present depends on the levels of processing involved in the employed experimental paradigm. Take once again the spatial attention study. Between the moving dots’ light hitting the participant’s retina and their pressing of the button to indicate motion direction, a vast amount of subcortical and cortical processing transpires. While internal dynamics start to coordinate stimulus processing as early as the thalamus [53,54], or even earlier [55], disharmony is most prevalent in higher level regions – where processes such as attention and decision-making operate in full swing. This is because at this late stage, information has passed through many cell ensembles where different oscillations have each exerted a temporal footprint. As these footprints add up, clock time falls increasingly out of tune with brain time. As a result, we lose track of brain dynamics and the information they provide about cognition, such as whether dots make it to awareness [56] or whether features of the dots (such as their colour) are available to working memory [57]. Thus, there is a gradation in clock time’s distorting effect which depends on the extent of cortical processing. Research that restricts analyses to low-level subcortical processing or very early evoked potentials benefits comparatively little from retuning, while retuning may be a vital step to understand brain patterns in late stages – with at least one exception: retuning is unnecessary for analyses that do not rely on temporal variations of the neural signature. For example, a researcher may want to study the trial-average activity in parietal regions as a function of the proportion of dots that moved, or map aggregate differences in network connectivity between participants. Here it is true as ever that brain dynamics play their part in the subprocesses involved, but the analyses are insensitive to time variations, leaving them unafflicted by oscillations’ footprint.

In short, retuning clock and brain time is necessary depending on the degree to which both the mechanisms of study and the employed analyses depend on the brain’s internal dynamics. On the whole, few electrophysiology studies are exempt from clock time’s distorting effects.

### Brain time is not unitary

The phrase “brain time” is used to emphasize the conceptual departure from a format extrinsic to the brain. We do not mean to suggest that there is a single brain time. Rather, different cognitive processes are clocked by different groups of oscillations, each with their own frequency and source. In this sense, brain time is analogous to a concept like “Earth time”, which contrasts itself with time on other planets whilst further decomposing into different time zones. Finally, brain time in the present context does not refer to *timekeeping* in the brain. Instead, it refers to the oscillatory dynamics by which the brain coordinates cognition generally, of which temporal cognition is a specific instance [58].

### Conclusion

Where does this leave us? We believe that rather than imposing a foreign unit of time onto the brain while studying its function, the brain is best understood as a system with its own dynamics and temporal organization. Upon inspection, the brain operates rhythmically, with brain oscillations as a key player. This has important consequences for scientific analysis. If it is true that oscillations clock brain activity to coordinate cognition, then their dynamics should heavily inform how we study evolving data patterns. In contrast, analysing such patterns in the default clock time format is likely to yield a distorted readout as the brain’s internal dynamics do not scale linearly to sequences of (milli)seconds. We introduce brain time warping as a method to account for disharmony in brain data, dynamically transforming electrophysiology data structures in a way that brings them in line with brain dynamics. The Brain Time Toolbox implements brain time warping, facilitating analysis on the oscillatory dynamics of the cognitive brain.

## Data availability

We re-structured and re-analysed the data of [41], and simulated new data, which are available in the Brain Time Toolbox at https://github.com/sandervanbree/braintime. We re-analysed the data of [42], which is available at https://osf.io/bpexa/.

## Code availability

Code for the brain time analysis of the rodent and simulated data is included in the Brain Time Toolbox at https://github.com/sandervanbree/braintime. Custom code for analysis of the human data is available from the corresponding author upon request.

## Acknowledgments

We thank Ehren Newman, Federica Meconi, Gi-Yeul Bae, and Steven J. Luck for sharing their data, and Benjamin Griffiths for his insightful comments on the Brain Time Toolbox. This research was funded by the European Research Council (Grant Number 715714 for M.W. & 647954 for S.H.), the Economic and Social Research Council (ES/R010072/2 for S.H.), and the Autónoma University of Madrid (FPI-UAM 2017 for M.M.). The funders had no role in study design, data collection and analysis, decision to publish or preparation of the manuscript.

## Author Contributions

The Brain Time Toolbox was conceived and developed by S.B., M.M. & S.H., with consultation from L.K., C.K., & M.W. The simulated dataset was generated and analysed by S.B., M.M. & S.H. The remaining datasets were analysed by S.B. & S.H. The manuscript was written by S.B. & S.H. with input from M.M, L.K., C.K., & M.W.

## Supplementary Information

### Supplementary Methods

#### 1. Data analysis

##### 1.1 General methods

We tested whether brain time warping increases dynamic patterns of cognition by applying the Brain Time Toolbox (abbreviated *braintime*) to three datasets. We divided each dataset into two classes, with their difference in brain activity predicted to reflect the neural signature of cognition (e.g., the difference in activity between left and right cued trials reflects spatial attention). We tested to what extent brain time warping increases (1) general oscillatory patterns present in the data and (2) periodic fluctuations in neural signatures of cognition. The logic across all analyses is the same: insofar the oscillation used for warping temporally coordinates a studied cognitive process, transforming data according to its dynamics should enhance rhythmicity of associated brain patterns.

###### Event-related potentials

Brain time warping is expected to overcome frequency drift, variable starting phases, and phase jumps, which in conjunction should result in a strong oscillatory structure in the event-related average, where trials are collapsed. We tested this hypothesis by applying a timelocked analysis in *FieldTrip* (Oostenveld et al., 2011) to each of the three datasets, separately for both conditions. We then selected a channel near the location of the warping signal to obtain a qualitative indication that brain time warping increases the simulated or predicted oscillatory structure (with similar results for neighbouring channels).

###### Power spectrum

We obtained Welch’s power spectral density estimates for each trial and channel per participant. We used the *fitting oscillations & one over f* (FOOOF) toolbox (the MATLAB implementation; Donoghue et al., 2020) to remove the aperiodic 1/f component, after which we averaged across trials and channels per participant. As such, each power spectrum we display shows the average model fit with the aperiodic component removed. For each dataset, we analysed the clock time data from 2 to 30 Hz. For brain time spectra, we re-referenced the frequency space to each participant’s individual warping frequency – which we then aligned in terms of spectral resolution and frequency space via the method explained in 2.3.4 *(Filtering to common frequencies).* We equalized the y-axis (power) between clock and brain time according to the minimum and maximum values between them to enable a direct visual comparison.

###### Intertrial Coherence

For each dataset, we used *FieldTrip’s* `ft_freqanalysis` to obtain the Fourier spectrum of each channel and trial, using a wavelet analysis that fits 5 cycles. We calculated the intertrial phase coherence (ITPC) across time and frequency for each participant in the following way. We divided the Fourier spectrum by its amplitude, summed the resulting angles, took the absolute value and normalized based on the number of trials (adapting the scripts provided by https://www.fieldtriptoolbox.org/faq/itc/). We averaged ITPC across all channels within participants, and then averaged across participants – separately for clock and brain time. We used the same brain time frequency re-referencing, spectral alignment approach, and axis equalization step as described in the previous paragraph.

###### 1.1.2 Advanced analyses

We tested the prediction that brain time warping increases the oscillatory structure of neural signatures of cognition by applying a linear discriminant analysis (LDA) to each dataset and by subsequently analysing its performance over time. Specifically, we generated temporal generalization matrices (TGMs) for each participant and applied a Fast Fourier Transform (FFT) over each of its rows and columns. We refer to this procedure as a “periodicity analysis”. For extensive details on periodicity analysis (as well as details on *braintime* terminology and all other analytical steps), we refer to *braintime’s* methods section (2.1 to 2.4). In this section, we first describe information common to the analysis of each dataset, before discussing them individually.

###### Classification

We compare second-level periodicity results between clock time data (input to the toolbox) and brain time data (output of the toolbox). To do so, we trained an LDA with 5 folds and 10 repetitions on the data and obtained 50 empirical TGMs and a null distribution of 50 permuted TGMs (obtained by class label shuffling, 2.3.3). Classifier performance was defined in terms of accuracy (i.e., the proportion of correctly classified trials). We used *braintime* to quantify periodicity in the TGMs, which yielded *n* empirical periodicity spectra and 50 × *n* permuted spectra, where *n* is the number of participants of each dataset. To obtain second-level results, we set *braintime* to obtain 10^6^ permuted power values for each periodicity frequency by randomly grabbing (with replacement) from the total pool of permuted TGMs, from which p-values are calculated (2.3.4).

###### Cluster correction

We established whether the classifier was able to differentiate both classes of data above chance using cluster correction in *MVPA Light* (2.3.4). We use 10^4^ permutations to test for significant clusters.

###### Warping source extraction

For the simulation and human analyses, we use independent component analysis (ICA) to decompose the input clock time data into additive subcomponents with statistically independent characteristics. Each component served as a warping source, meaning each contained a potential warping signal that could be designated as brain time.

After brain time warping, we removed the warping source (ICA component) that was used for warping from the data before continuing to periodicity analyses. This removes the warping signal from the output data, introducing data independence (2.4.1).

For the rodent analysis, we warp using local field potential channels, and introduce independence by removing the selected channel from the data, leaving only the warped single-unit data.

###### Brain Time Toolbox settings

We used *braintime’s* default *consistenttime* method to ensure the brain time warped data restricts itself to the specified time window for each analysis (2.4.2). To obtain time frequency information of each warping source, we applied a wavelet analysis (fitting 5 cycles to the data). To correct for multiple comparisons across the tested frequencies in the periodicity spectra, we adjusted p-values for the false discovery rate (FDR). We implemented the FDR correction as described in Benjamini, & Yekutieli (2001), which is optimized for analyses with statistical dependencies between tests – as is the case with frequencies across the periodicity spectra.

##### 1.2 Simulation

We simulated a basic attentional spotlight model (main Figure 4) with dipoles oscillating at an alpha frequency (8 to 12 Hz). To introduce disharmony between clock and brain time, we introduced variable starting phases and frequency drift in the primary dipoles. We predicted that brain time warping would selectively increase periodicity at the ground truth frequency by adjusting for non-stationarities in the warping signal. *Braintime* contains the MATLAB (The MathWorks, Natick, Massachusetts, USA) code used to generate the simulated dataset, which includes options to change simulation parameters (such as the degree of frequency drift, pink noise amplitude, the number and duration of trials, and so forth).

###### 1.2.1 Methods

We used *FieldTrips* (Oostenveld et al., 2011) `ft_dipolesimulation` function to generate an electrophysiology dataset where the ground truth consists in a periodic fluctuation of covert attention at alpha frequencies. We simulated *n =* 10 datasets with 120 trials and two conditions (covert attention to the left and right hemifield; LHF and RHF). Each trial lasted one second and had a sampling rate of 200 Hertz (Hz). Across both conditions and all participants, we held constant the orientation and location of the primary dipoles that instantiated the neural signature of attention. The remaining parameters were randomized, such as the frequency, location, and orientation of non-primary dipoles (“random” dipoles), as well as each dipole’s starting phase. We modelled 3 primary dipoles, and 8 random dipoles. The primary dipoles contained different parameters across the two classes of data. In contrast, the random dipoles contained parameters that were uncorrelated to each class, yielding no meaningful average difference in neural activity between conditions. In this simulation, our primary focus was to create a dataset suitable to test brain time warping (with a neural signature and non-stationarity) rather than to accurately model attention in close accordance with theoretical models.

###### Primary dipole configuration

We placed the conductor dipole in the right parietal cortex with an amplitude of 1 (in arbitrary units), oscillating at a random alpha frequency. This conductor dipole “conducted” the phase of two follower dipoles, one located in each visual cortex, oscillating at the same frequency and oriented with the same x, y, and z coordinates. We generated two classes: 60 trials of LHF and RHF data. The two conditions contained a difference in parameters – and thereby brain activity – that the classifier could exploit to differentiate the data and obtain the ground truth signature of attention. Specifically, for both conditions, we changed two parameters:

- The relative amplitude between follower dipoles. The follower dipole contralateral to the hemifield condition oscillated at half the amplitude (0.5) compared to the ipsilateral dipole (1) – mimicking contralateral inhibition of alpha.
- Phase relation between primary dipoles. The conductor dipole oscillated in antiphase with the follower dipole contralateral to the hemifield. In contrast, it oscillated in phase with the follower dipole ipsilateral to the hemifield (barring a 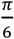 phase delay to mimic signal conductance delay).

In addition to these condition-specific differences, each primary dipole contained 1/F noise, variable starting phase, and frequency drift. We implemented frequency drift through a random walk approach, where the frequency of all three primary dipoles decreased or increased by 0.05 Hz per sample.

###### Random dipole configuration

To decrease the signal to noise ratio of the neural signature exerted by primary dipoles, we introduced 8 random dipoles that oscillated at random frequencies, limited to 2 Hz above and below the primary dipole’s frequency to avoid interference with the neural signature. These dipoles oscillated at an amplitude of 1, contained 1/F noise, and were placed randomly across the brain with a random orientation.

###### Brain time warping

We considered the resulting datasets as separate participants with their own alpha frequency. We used ICA to obtain warping sources and warped each participant to a source with (1) a topographical activity profile around occipital and parietal regions, (2) and a high amplitude alpha peak. We found that brain time warping unveils periodicity at the warping frequency which was absent in clock time. This demonstrates that the variable starting phases and frequency drift preclude the classifier from detecting reliable differences in neural activity between both classes of data. However, after brain time warping has been applied, the non-stationary dynamics of the primary dipoles are accounted for, resulting in a significant effect at the warping frequency (p = < 0.005; main Figure 4). We provide TGMs, autocorrelation maps and first-level periodicity spectra in 3.1.2.

###### Advanced analysis

To test for the specificity of brain time warping, we warped to the golden mean of each participant’s warping frequency as obtained in the main analysis (Supplementary Figure 1). The logic is that if brain time warping is non-specific and prone to false positive results by trivially imposing oscillatory structure, then it should do so even for frequencies predicted to not have a clocking role in cognition. To test this possibility, we repeated the main analysis by warping to the golden mean of each participant’s warping frequency (≈ *warping frequency ×* 1.618). We chose the golden mean as our control frequency because, at this frequency, the warping signal (which is predicted to be the clocking oscillation) is least expected to synchronize to different oscillations (Pletzer, Kerschbaum & Klimesch, 2010). The control analysis changed the warping frequencies from alpha (8 to 12 Hz) to approximately 12.9 Hz to 19.4 Hz. All other parameters were unchanged from the main analysis. The control analysis does not show significant periodicity at any frequency. We provide TGMs, autocorrelation maps and first-level periodicity spectra in 3.1.3.

**Supplementary Figure 1:**
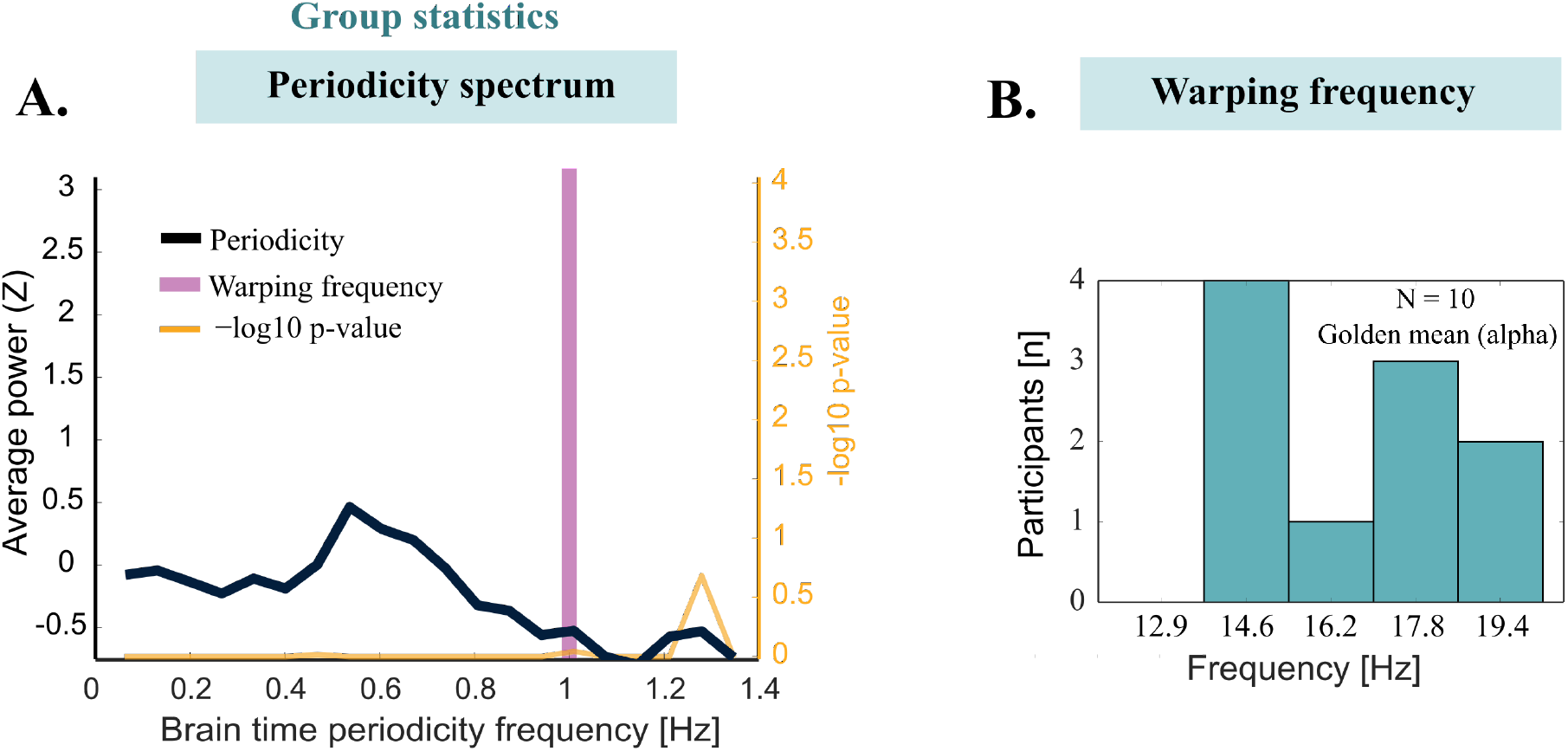
Advanced control analysis of the simulation dataset. **(A)** We found no significant periodicity when warping to the golden mean of each participant’s alpha oscillation. **(B)** Summary statistics of the warping frequency across participant.

These results demonstrate that brain time warping does not trivially impose oscillatory structure in high-level analyses by virtue of warping to a specific frequency if there are no such signatures hidden in the data. We next established whether the same conclusion extends to the family of basic analyses. Specifically, warping should not manifest in prominent peaks in the power spectrum, or high values in the ITC, at warping frequencies when those frequencies are only minimally present in the data.

###### Basic analysis

We repeated the control analysis to test for the effects of brain time warping on basic results like power spectra and ITC. As explained in the main text *(Is retuning using brain time warping circular?),* we expect brain time warping to at least show some increased power and ITC at any frequency because electrophysiology data is made up of different frequencies of oscillations which each have at least some non-stationarities that can be reduced. However, it should be the case that warping to oscillations not primarily simulated should result in lower power and ITC increases than warping to oscillations primarily simulated. To test this, we changed the main frequency of the random (1/f) dipoles to match the individual alpha frequency of the primary dipoles – strongly clustering the ground truth signals around alpha. We then again warped to the golden mean of alpha. We found that brain time warping to control frequencies only resulted in low spectral peaks and ITC at the control frequency (Supplementary Figure 2). In the power spectra, the primarily simulated frequency overshadows the brain time data even when warping to control frequencies.

**Supplementary Figure 2:**
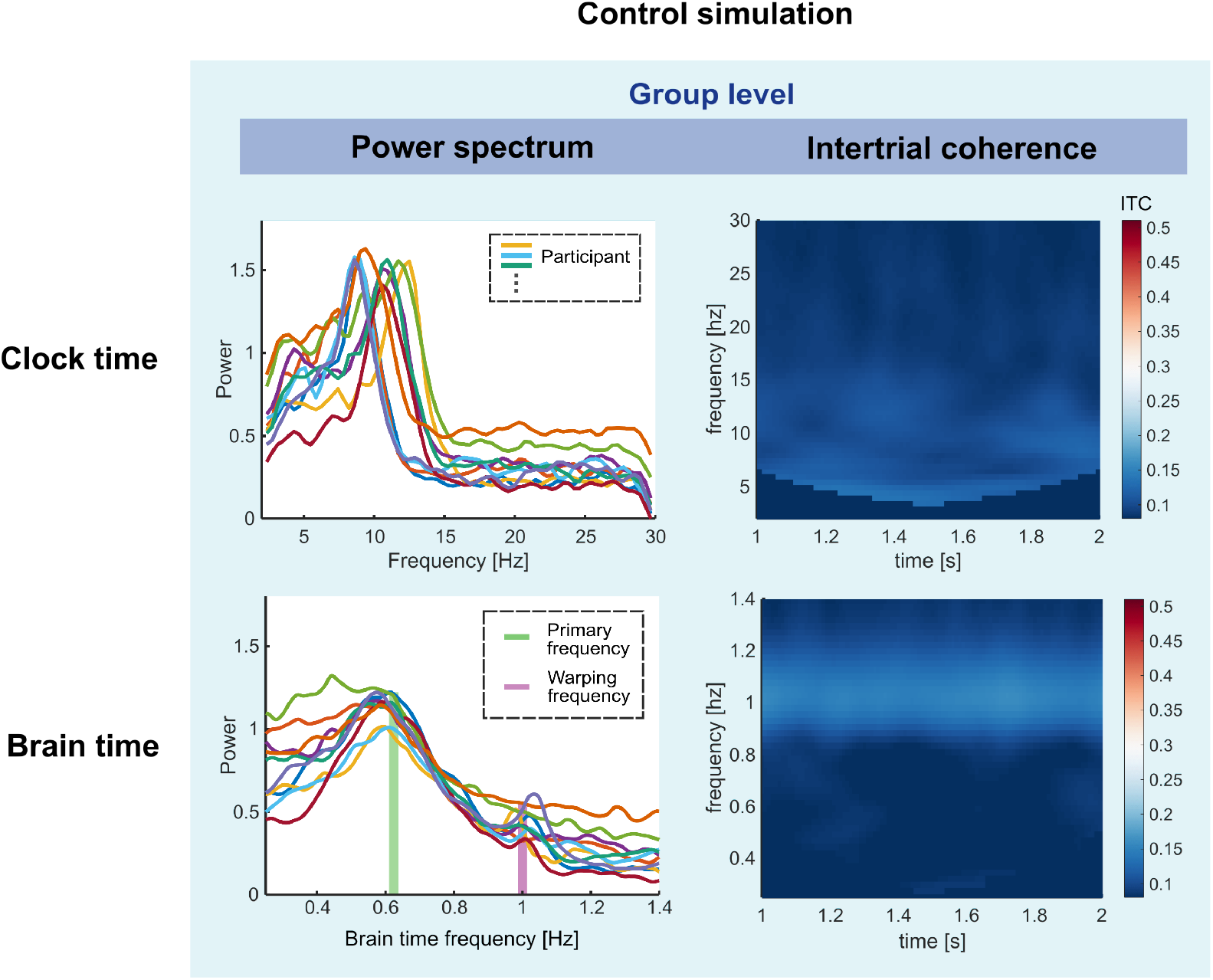
Basic control analysis for the simulation dataset. Brain time warping to control frequencies (i.e., at the golden mean of the primarily simulated frequency) resulted in weak increases of power and ITC at control frequencies. The brain time data shows stronger peaks at the primarily simulated frequency. The color axis of ITC is held constant from the main analysis to enable visual inspection of the difference in magnitude between main and control analyses.

###### 1.2.3 Nested oscillations

What is the effect of brain time warping if there are multiple nested oscillations that coordinate cognition? To address this question, we simulated a single dataset with two sets of nested oscillators that were cross-frequency coupled (Jensen & Colgin, 2007). Specifically, the nested oscillators were phase-amplitude coupled, and we added phase-phase coupling to further enhance the nested nature of the oscillators (Siebenhühner et al. 2016). One set of oscillators was in the left, and one set was in the right hemisphere (Supplementary Figure 3A). Within each set, one oscillator fluctuated at a high frequency and one at a lower frequency. Similar to the simulation in Supplementary Methods 1.2.1, we generated a different phase relation between two hemifield conditions to create a classifiable neural signature. We tested whether warping to the lowest frequency oscillator reveals periodicity at the frequency of the nested high frequency oscillator. This simulation script is added in *braintime* under `bt_dipsim_cfc` and the relevant analyses are provided and documented in `tutorial8_crossfrequency`.

**Supplementary Figure 3:**
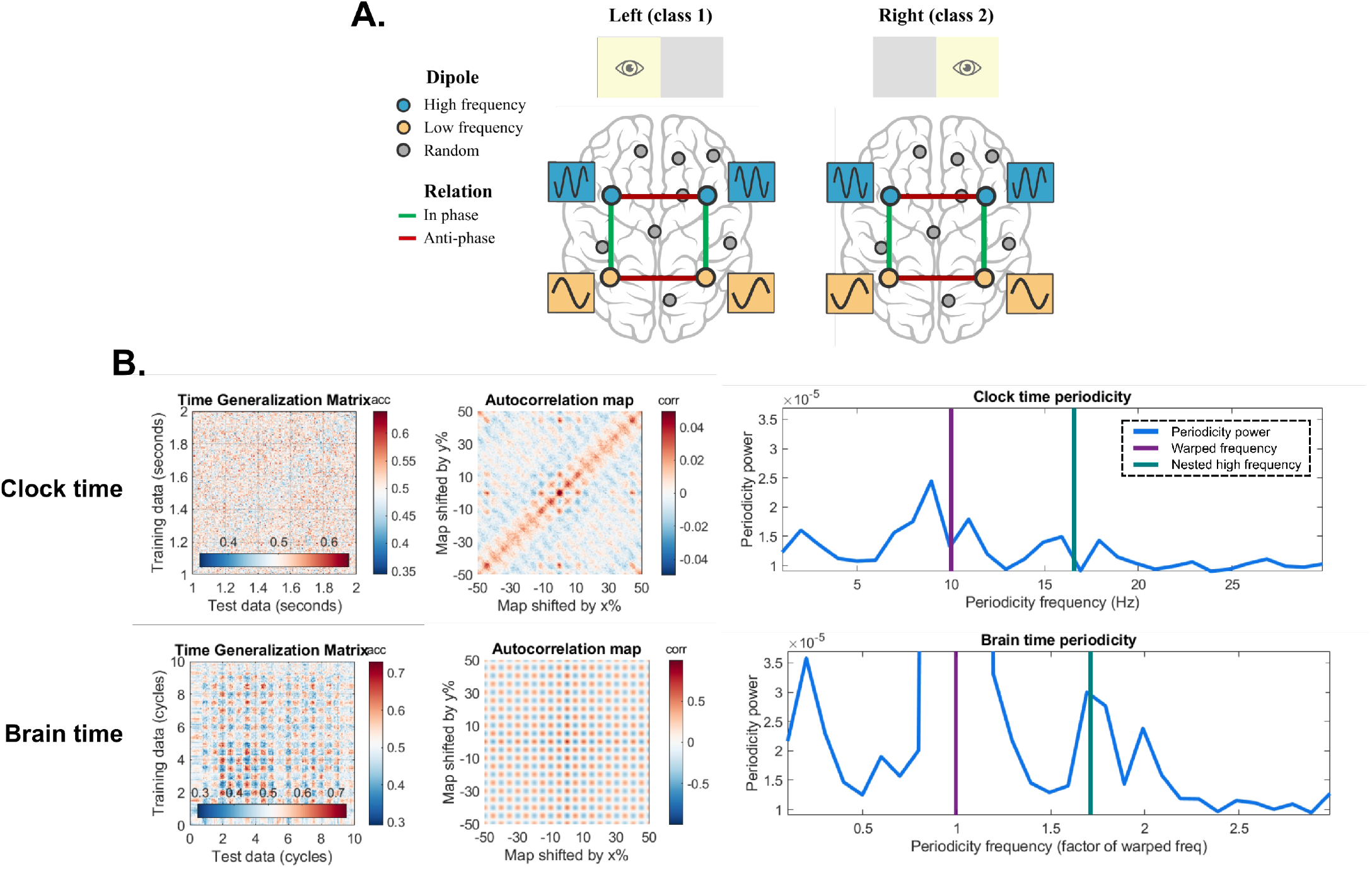
Dipole configuration for the nested oscillation simulated dataset. **A)** Dipole configuration for the nested oscillation simulation. A high and low frequency oscillator in each hemisphere were nested by means of phase-amplitude and phase-phase coupling. The two sets of nested oscillators had an anti-phase relationship with respect to each other, resulting in a classifiable neural signature that varies between left and right hemifield conditions. **B)** Brain time warping shows a stronger peak at the frequency of the nested oscillator that was not apparent in clock time. These results are inconsistent across repetitions.

We found that brain time warping is able to reveal the high frequency oscillator when warping to the low frequency oscillation’s frequency under some circumstances (Supplementary Figure 3B). However, the magnitude of the effect is small compared to standard analyses where we tested for periodicity at the warped frequency (i.e., in the simulation where no nested oscillations are present; main manuscript). Moreover, generating the same dataset several times – which introduces slight ground truth variability due to randomness in the simulation – showed that warping-induced periodicity increases around the coupled high frequency arose inconsistently across analyses. This suggests that while brain time warping can be used to study cross-frequency coupling, it is not optimally suited to do so and better methods are available (see Hülsemann, Naumann & Rasch, 2019 for an overview). From a different perspective, these results do demonstrate that if one is solely interested in the brain dynamics orchestrated by a single class of oscillators, brain time warped results are unlikely to be contaminated by extraneous brain dynamics – because even maximally coupled oscillators infrequently impinge on each other.

##### 1.3 Rodent

We brain time warped a rodent dataset obtained from Newman & Hasselmo (2014). In this study, local field potentials (LFP) and single units were measured from the entorhinal cortex of Long-Evans rats as they navigated through a rectangular open field. In the following section, we describe how we reformatted and analysed the data to test the effects of brain time warping. For methodological details on the recordings such as data acquisition, surgical procedures, and behavioural protocols, we refer to Newman & Hasselmo (2014). We hypothesized that brain time warping would increase the periodicity with which a classifier is able to differentiate when an animal is inside or outside the firing field of a grid cell. To this end, we brain time warped grid cell activity to theta oscillations in the LFP. The logic is that if theta clocks grid cell activity, then periodic patterns in their spiking activity should be enhanced after brain time warping (through its correcting of non-stationarities in brain time).

###### Primary grid cells

We inspected single cell recordings from 7 recording sessions (N = 2 rodents) and extracted one primary grid cell for each recording, which we used to define the in- and outfield condition. We selected a primary grid cell by comparing each cell’s firing rate map for triangular structure and the periodicity of these maps using a 2D autocorrelation analysis (Hafting, Fyhn, Molden, Moser & Moser, 2005). In 3.2.2, we display the smoothed firing rate maps of each session’s primary grid cell and how we devised classes based on their firing fields.

###### Spike-field coupling

Brain time warping is only expected to enhance periodicity if grid cells phase lock to theta. Thus, we verified spike-field coupling for selected grid cells. In 3.2.2, we show the spike-field coupling of one example grid cell.

###### Data preparation

For each session, we created a separate structure for the spike activity of grid cells (between 2 and 4 channels; convolved with a Gaussian) and LFP recordings (2 channels). We split each structure into two classes of data based on when rodents were inside a location coded by the primary grid cell’s firing field (defined by local maxima in the smoothed firing rate map) or outside of it (defined by local minima). As a consequence of this split, the data at t = 0 reflected the animal’s presence in a firing field (class 1), or not (class 2). This trivially introduces high classifier performance around these timepoints. However, we were not interested in the classifier’s performance to differentiate in- and out-field conditions. Rather, we tested whether brain time warped data increased periodicity in classifier performance at the warping frequency. We brain time warped 2.5 seconds of data (from 1.25 seconds before to after t = 0). The number of trials per session ranged from 60 to 168 (mean: 104.8 trials, standard deviation: 39).

###### Brain time warping

The average waveshape of LFP theta was asymmetric, so we generated the clock time template signal (used to detect segments of non-stationarity in LFP theta) based on a smoothed version of the average waveshape rather than a symmetric sine wave (which is the default method). This optimizes brain time warping by reducing differences between the clock and brain time signal, improving the accuracy of the warping path (see Toolbox methods). In 3.2.4, we show TGMs and their autocorrelation map for each session. The prepared data and scripts used for analysis are included in the Brain Time Toolbox.

##### 4.4 Human

We brain time warped a human dataset obtained from Bae & Luck (2019), which is available for download at https://osf.io/bpexa/. In this study, electroencephalography (EEG) was recorded while participants viewed random dot kinematograms, where a proportion of dots moved coherently toward a direction in 360° space. Each trial contained, in order, a fixation period of 1.5 s, a motion period of 1.5 s, and a direction report of variable duration. Again, we restrict our description to how we reformatted and analysed the data. For the full methodological details, see Bae & Luck (2019). We hypothesized that brain time warping to occipital-parietal alpha oscillations would increase the periodicity with which a classifier is able to differentiate whether motion is leftward or rightward, resting on evidence for alpha’s role in controlling spatial attention (Capotosto, Babiloni, Romani & Corbettam, 2009).

###### Data preparation

First, we pooled both recording sessions and coherence levels (with 25.6% and 51.2% of dots moving coherently in one direction). We used only the 1.5 s motion period for decoding, as we predicted the neural signature of spatial attention is most detectable during motion. We filtered each participant’s data (N = 16) to the 80% most accurate trials (defined based on the error between reported and presented motion angle), reducing the number of trials from 1280 to 1024. This removes the subset of trials where a rare reversal effect occurred in which participants report motion in roughly the opposite direction to the presented motion. Next, we defined two classes of data based on whether motion was leftward (class 1; angles between 90° and 270°), or rightward (class 2; the remaining angles).

###### Brain time warping

We extracted warping sources by applying ICA to the data. For each participant, we chose a component with sustained alpha power and a topographical profile that suggested an occipital-parietal origin. In 3.3.2, we show TGMs and autocorrelation maps for each participant.

#### 2. Toolbox

##### 2.1 Introduction

The Brain Time Toolbox, abbreviated *braintime*, is a MATLAB software library designed for use with electroencephalography (EEG), magnetoencephalography (MEG), and single and multiunit intracranial recordings (Supplementary Figure 4). The toolbox has two primary uses: brain time warping data structures (transforming them from clock to brain time; operation 1) and testing for periodic patterns in the data that reflect dynamic cognition (operation 2). The standard pipeline is to brain time warp data and test for a change in periodicity at frequencies of interest. However, each operation can be carried out without the other – brain time warped data may be analysed outside the toolbox and periodic patterns may be tested on data that has not undergone warping (clock time data).

**Supplementary Figure 4:**
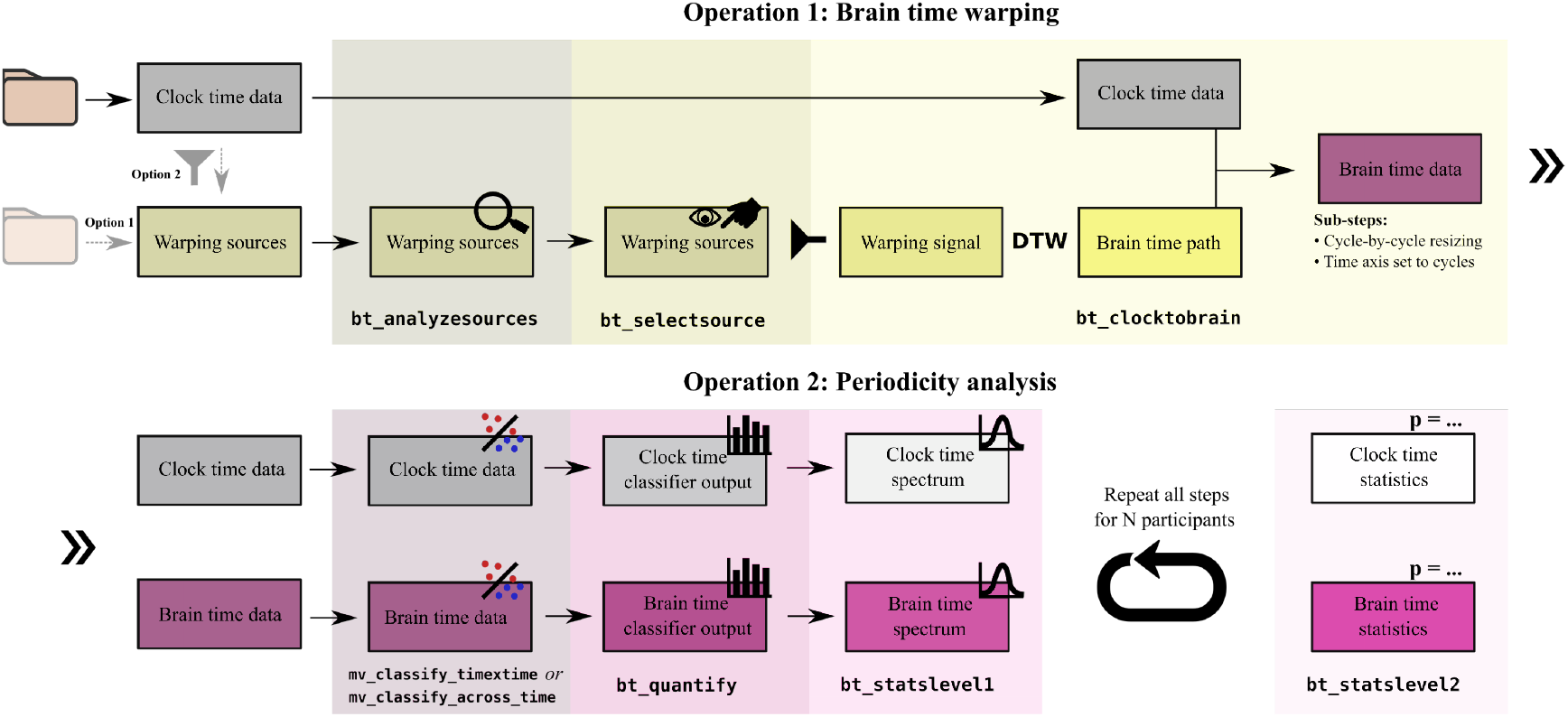
the default Brain Time Toolbox pipeline. *Braintime* has two primary operations. **(Top row)** For brain time warping, the user first loads a FieldTrip formatted data structure that will be transformed (clock time data). Separately, a data structure with warping sources is required. This structure contains potential warping signals – the oscillation used to transform the data. The warping sources may be loaded separately (option 1) or extracted from the clock time data (option 2). `bt_analyzesources` analyzes each warping source for warping signals based on user parameters. Next, using `bt_selectsource`, the user can inspect each warping source’s characteristics (such as spectral information) and designate a warping signal as brain time. Finally, `bt_clocktobrain` dynamically time warps (DTW) the warping signal to either a stationary sine wave or the warping source’s average waveshape. The resulting warping path is applied to the clock time data, cycle-by-cycle, and resized to the original length. The function also sets the time axis from seconds to cycles of brain time. **(Bottom row)** For the periodicity analysis, *braintime* first calls MVPA Light to test a classifier on two classes of data, prespecified by the user. The resulting output is one dimensional for `mv_classify_across_time`, or two dimensional, when testing for time generalization of the classifier using `mv_classify_timextime`. Then, across this output, `bt_quantify` tests for periodic patterns in classification performance, resulting in a periodicity spectrum. Next, `bt_statslevell` performs first-level statistics by comparing the empirical spectrum with a user-specified number of permuted spectra (obtained by shuffling class labels). Finally, `bt_statslevel2` obtains group level results of the classifier’s periodicity, and calls MVPA Light to perform cluster correction over the classifier’s performance.

In this manuscript, we describe *braintime* in extensive technical detail with limited emphasis on user flow. For a practical guide, we recommend the more concise and step-by-step documentation on *braintime*’s Github page (https://github.com/sandervanbree/braintime). For a quick start, we recommend *braintime’s* tutorials.

###### 2.1.1 Requirements

*Braintime* interfaces with the following software packages, requiring each to be installed:

- MATLAB Signal Processing Toolbox (The MathWorks)
- FieldTrip (Oostenveld et al., 2011)
- MVPA Light (Treder, 2020)

Besides these software dependencies, *braintime* requires the electrophysiology data to be structured in a FieldTrip format. For data structures that are not in this format, FieldTrip has built-in tools to convert a wide array of common file types. See the FieldTrip documentation for information on how to convert data.

In case *braintime*’s second operation will be used (testing for periodicity), the input clock time data needs to consist of two classes of data hypothesized to yield the neural signature of the studied cognitive process. Moreover, trials in the data structure need to be labelled based on their class membership. For more details on how to do this, see section 2.3.

###### 2.1.2 Download and installation

Before setting up *braintime*, ensure FieldTrip and MVPA Light are downloaded and fully initialized (download links above). *Braintime* can be downloaded on the toolbox Github page. First, download the ZIP folder and unzip it to a preferred path. Then, run the function `setup_braintime` (in the setup folder). To get started with the toolbox, see *braintime’s* tutorials.

##### 2.2 Operation 1: Brain time warping

Brain time warping transforms data according to the phase dynamics of an oscillation hypothesized to clock the studied cognitive process. This changes the time axis of the data from seconds to cycles of the warping frequency.

###### 2.2.1 Obtaining warping sources

Warping sources may either be ad hoc extracted from the clock time data or loaded from a separate data structure. The warping sources contain warping signals – potential oscillations used for brain time warping. There are only two restrictions to what constitutes warping sources: the data structure needs to be in a FieldTrip format, and the data needs to align in time with the to-be-warped clock time data. Users are encouraged to decide on a case-by-case basis what data serve as the best warping sources. Below, we describe two methods of obtaining warping sources, and provide several examples for each.

###### Ad hoc extraction

Warping sources may be extracted from clock time data on the fly, for example using independent component analysis (ICA). ICA is a popular method to decompose electrophysiology data into additive subcomponents (Makeig, Bell, Jung & Sejnowski, 1995). The resulting components aggregate data with statistically uncorrelated spectral characteristics, making for suitable warping sources. Each component contains potential warping signals. The toolbox automatically detects when the warping sources are ICA components, and suggests removal of the selected component from the brain time warped data when continuing analyses outside the toolbox. For single- and multiunit recordings, an example of ad hoc extraction is to separate one (or more) local field potential channels into a warping source(s) structure.

###### Preloaded data structure

Warping sources may also be obtained from a separate preloaded data structure, in the form of previously extracted data. An example could be a study where magnetometer and gradiometer channels underwent separate preprocessing steps, with their data saved in separate files. Or perhaps a user has created a set of virtual channels using source localization, intending to warp sensor-level data using source-level phase estimation. Warping sources may also be obtained from a separate recording altogether. For example, a user may opt to warp EEG data using warping signal obtained from MEG data, or vice versa.

###### 2.2.2 Analysing warping source

In `bt_analyzesources`, *braintime* uses FieldTrip to perform time frequency analyses on each warping source. Users may choose between a variety of time frequency analysis parameters based on FieldTrip’s ft freqanalysis, and may opt to include 1/F correction. In addition, users select a range of frequencies predicted to contain the coordinating oscillation. For example, a user interested in spatial attention may specify a range of 8 to 12 Hertz (Hz). The toolbox estimates the phase of high power oscillations in this range in anticipation of brain time warping. Upon completion of the time frequency analyses, the toolbox sorts the warping sources based on power in the range of interest. Optionally, for EEG and MEG, the sorting of warping sources may be weighted by the hypothesized topographical activity of warping sources. For example, a researcher may want to focus on warping sources with parietal activity, or a lateralized source. To enable topography weighting, users first call `bt_templatetopo` and draw a topographical profile on a template layout. *Braintime* then loads the template topography and adapts the ranking of warping sources to their topography’s correlation with the template topography, instead of the default setting which sorts solely by maximum power at frequencies of interest.

###### Asymmetry and waveshape

For each warping source, *braintime* detects the average waveshape and the asymmetry of each cycle in it. This allows users who have predictions about the symmetry of their coordinating oscillation to decide between warping sources. In addition, the amount of asymmetry may also inform parameter selection during subsequent steps in *braintime*.

The waveshape and asymmetry of data are obtained via the following steps. First, *braintime* applies a band-pass filter in a 2 Hertz (Hz) window around the warping frequency. Then, trial-by-trial, the toolbox finds local minima and maxima in the filtered data, reflecting troughs and peaks, respectively. Then, based on the location of these minima and maxima, *braintime* extracts associated data in the (non-filtered) warping sources, calculating a peak-triggered wave. Thus, the peaks are located in the filtered data, while the shape of waves themselves are obtained by extracting data samples in the raw data around the peak locations. This is repeated for all cycles. Then, the asymmetry is calculated for each cycle, while the average waveshape is obtained by averaging each cycles’ data. We defined asymmetry as the difference in duration between the ascending and descending flank of a cycle, normalized by its duration (Belluscio et al., 2012):

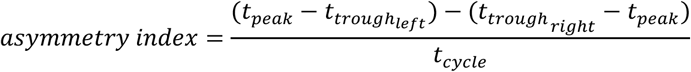

###### 2.2.3 Selecting a warping signal

In `bt_selectsource`, the user selects a warping source and its associated warping signal, designating brain time. In this step, *braintime* creates a graphical user interface that displays each warping source’s information as calculated by `bt_analyzesources` (Supplementary Figure 5). This information comprises (1) a textbox with recommended criteria for warping source selection and browsing instructions, (2) the topography of the warping source (if requested), (3) the asymmetry and shape of the average waveshape, (4) the time frequency representation of the warping source’s data, and (5) its power spectrum. Users may visually inspect each warping source and combine all information to designate the warping signal predicted to clock the studied cognitive process of interest, in keeping with a hypothesis-driven approach.

###### 2.2.4 Warping clock to brain time

In `bt_clocktobrain`, the phase of the selected warping signal is used to warp the clock time data structure (Supplementary Figure 6). Before describing how dynamic time warping (DTW) is employed, we first explain two parameters that determine what DTW is applied to.

###### Warp method

We define clock time as a stationary oscillation, oscillating away faithfully to seconds. This is what the warping signal will be dynamically time warped onto. We implement two ways of generating this stationary oscillation. First, users may choose the default method, where a stationary sine wave is generated with MATLAB. However, for data with asymmetric waveshapes, using a sine wave with perfect symmetry may impede DTW’s attempt to minimize Euclidean distance between the phase of clock and brain time. Thus, we include the option to warp using a concatenation of the smoothed average waveshape of the warping source, yielding a sinusoid with matching asymmetry (Supplementary Figure 6).

###### Phase method

Besides control over what constitutes clock time phase, users may opt to estimate brain time’s phase in two ways. The standard way of estimating brain time’s phase is using Fast Fourier Transform (FFT) of band-pass filtered data around the warping frequency in the selected warping source. However, the toolbox also offers Generalized Eigenvalue Decomposition (GED) as an alternative method to estimate the warping frequency’s phase holistically across all warping sources. This method is appropriate when the warping sources are all considered to contain information about brain time. For example, the user’s warping sources may be local field potential channels in the hippocampus. In this case, the user may not want to choose one local field potential channel given their interest in the phase of theta oscillations across the entire hippocampus. Similarly, a user may have source localized parietal cortex channels which each provide valuable phase information. In these cases, GED is the optimal choice.

**Supplementary Figure 5:**
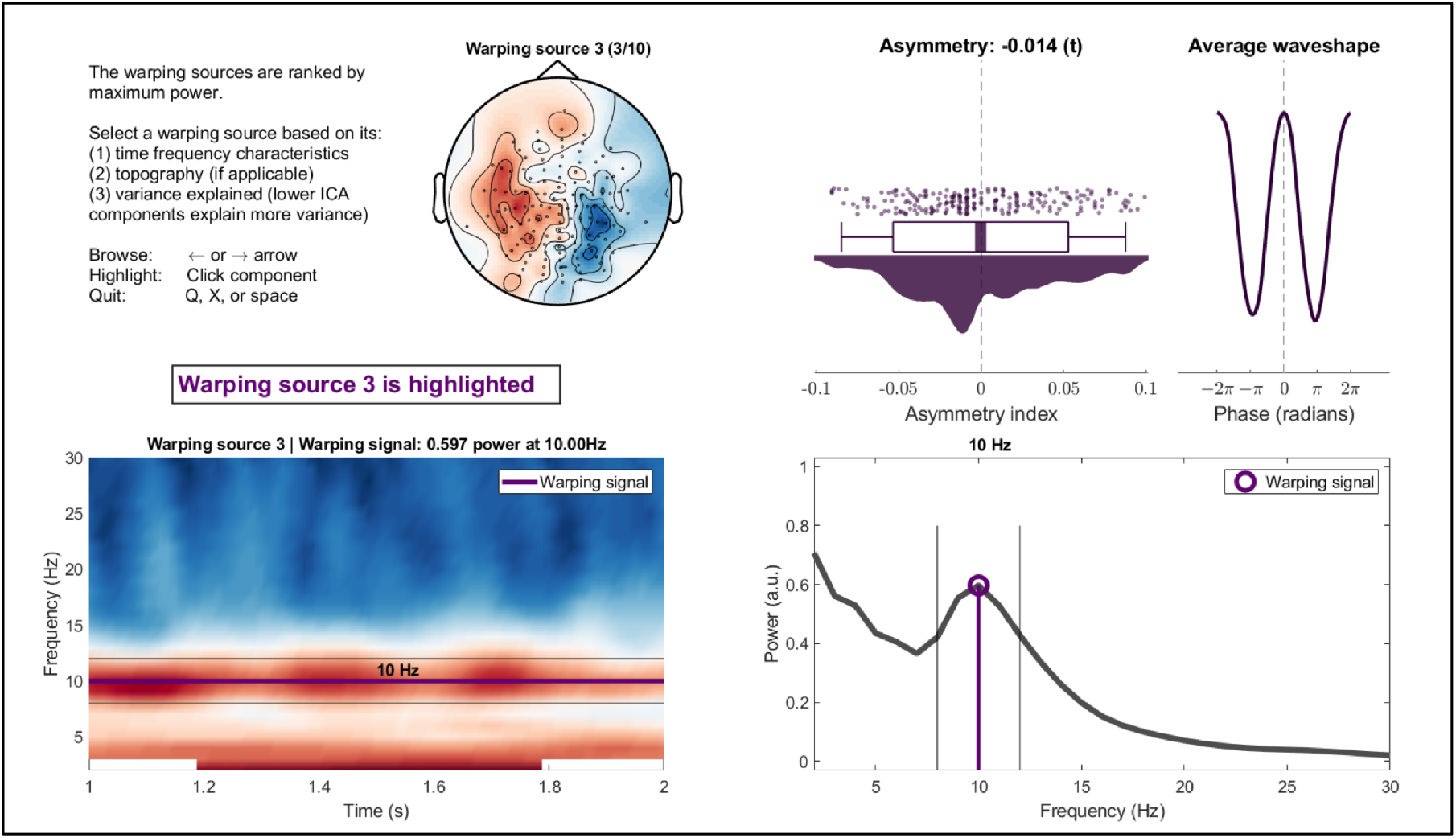
Selecting a warping source. The graphical user interface generated by `bt_selectsource`. **(Top row)** A textbox is displayed with suggested criteria for warping source selection, as well as instructions for browsing. Next, optionally for EEG and MEG, the topography is displayed. On the right, the average asymmetry of cycles in each warping source is displayed (one dot represents one cycle in the warping source). This serves two purposes: users may want to discriminate between sources based on their asymmetry, or check levels of asymmetry to inform parameter selection during subsequent steps. **(Bottom row)** The warping source’s time frequency representation is displayed, along with the power spectrum. For both plots, the borders of the frequency range of interest and the location of the warping signal are highlighted.

GED finds a spatiotemporal filter that differentiates all the warping source data into components with high and low power of the warping frequency (Parra, Spence, Gerson & Sajda, 2005; Cohen, 2017). When users specify this option, *braintime* extracts the component with the highest weight, yielding a time series of the GED component. The toolbox takes the angle of its Hilbert transform, defining brain time. To perform GED, the toolbox uses a modified version of the functions provided by Mike X Cohen (reported in Cohen, 2017). In short, FFT offers a classical and source-specific phase estimation of the warping frequency, while GED estimates the warping frequency’s phase holistically across warping sources.

###### Brain time warping

With both the clock time and brain time phase defined and computed, *braintime* initiates brain time warping. Here, the toolbox dynamically time warps clock and brain time onto each other. Clock time, here, is the unwrapped phase of a stationary sinusoid or average waveshape, while brain time is the unwrapped phase of the warping signal (estimated within a source using FFT or across sources using GED). These two phase vectors are dynamically time warped using MATLAB’s dtw command. DTW (Sakoe & Chiba, 1978) computes the optimal alignment between two signals, minimizing their Euclidean alignment. For a formal specification of DTW, see Berndt & Clifford, 1994 and Sakoe & Chiba, 1978.

**Supplementary Figure 6:**
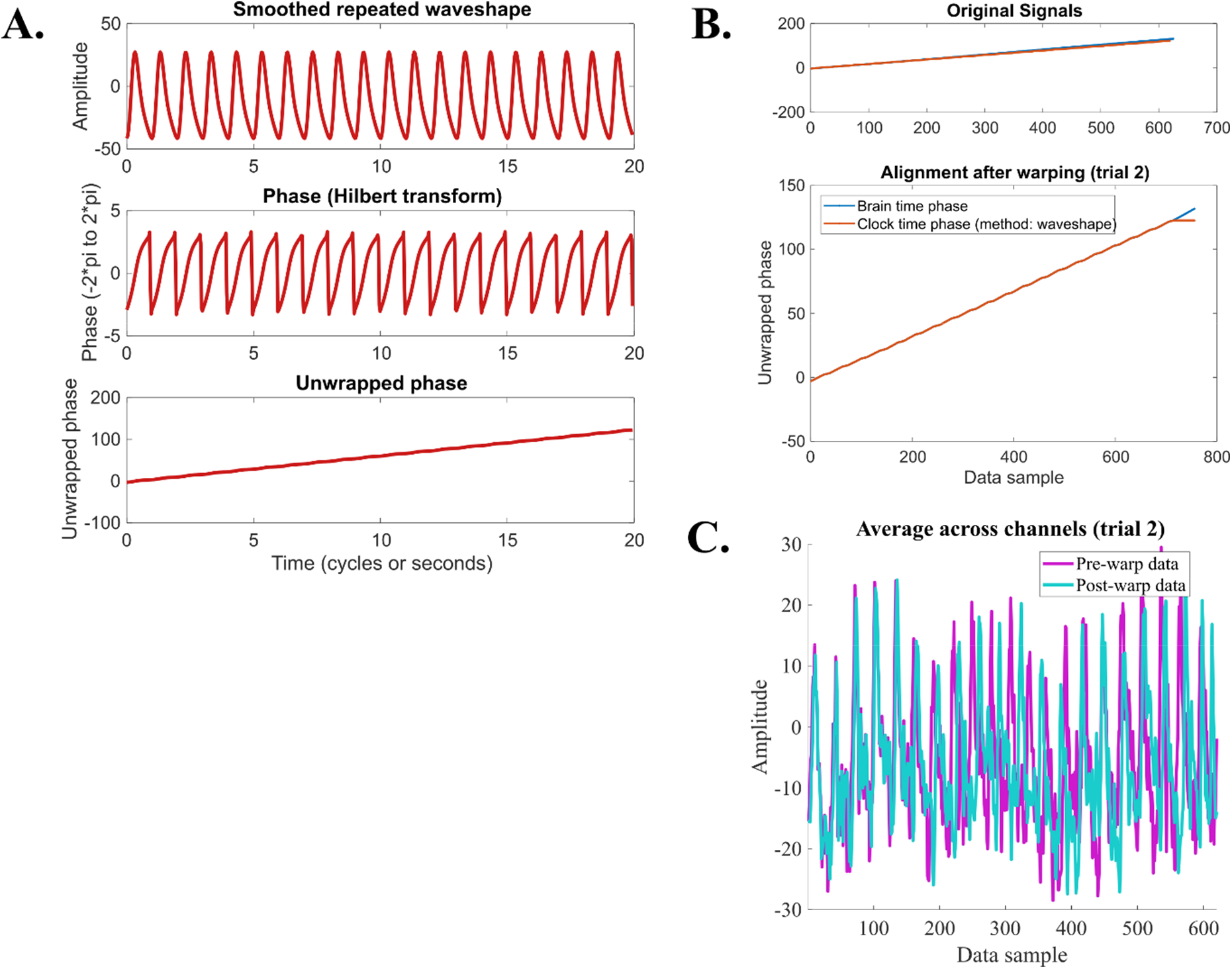
Warping clock to brain time. *Braintime* prints a series of plots upon completion of bt_clocktobrain to show the application of brain time warping to the data at hand. **(A)** In this example, the user has set *braintime* to warp to the smoothed average waveshape of the selected warping source (top). The toolbox extracts the angle of its Hilbert transform (middle) and unwraps the phase in preparation for warping (bottom). **(B)** The unwrapped phase vector of the warping signal and the stationary sine wave (representing clock time) are displayed (top), and so is their alignment after applying dynamic time warping (bottom). **(C)** Here, the warping path has been applied to the original clock time data, yielding a dynamic resampled version of the original data

The toolbox extracts the warping path between clock time (`ct_phs`) and brain time (`bt_phs`) using the following MATLAB code:

> [∼,ix,iy] = dtw(bt_phs,ct_phs)

`ix` is the warping path which, when applied to `bt_phs`, will minimize its distance to `ct_phs`, and vice versa for `iy`. Since `ct_phs` is a stationary signal, `ix` indicates (1) segments of non-stationarity in `bt_phs` (2) and how `bt_phs` needs to be resampled to become a stationary signal. Crucially, rather than applying `ix` to `bt_phs` to get it closer to `ct_phs`, brain time warping instead applies `ix` to the clock time data structure, transforming all of its dynamics based on the non-stationarities in the warping signal. The result is a data structure with dynamics adjusted in accordance with the temporal evolution of oscillations predicted to coordinate cognitive function, enabling an improved readout of its dynamics.

Whilst applying each trial’s `ix` to transform clock time data, the toolbox implements a cycle-by-cycle resizing. This is to ensure that the data of each cycle per trial has a constant number of samples. Without this step, if the warping path contains an extended segment of data repetitions for some trials, this would stretch out the data extensively, introducing a misalignment of brain time across trials. In other words, cycle-by-cycle resizing ensures brain time unfolds with a similar timecourse across trials by aligning its cycles. To resize data, the toolbox uses MATLAB’s `imresize` (which uses nearest-neighbour interpolation). *Braintime* determines the start and end of cycles using the warping path `iy` (the warping path from `ct_phs` to `bt_phs`), which has an equal length to `ix` but more closely reflects the stationary dynamics of clock time. An additional side-effect of cycle-by-cycle resizing is that the number of samples of the original data remains unchanged after brain time warping.

Upon completion of these steps, the data is in brain time. In light of this, the toolbox changes the time axis from milli(seconds) to cycles of the warping signal. Next, users may test for dynamic information patterns using operation 2 of the toolbox or continue analysis outside of the toolbox. In case of the latter, there are some circularity concerns that will be addressed in section 2.4.1. Before that, we describe the toolbox’ second operation.

##### 2.3 Operation 2: Periodicity analysis

*Braintime* allows users to test for dynamic patterns of information using machine learning techniques. This requires the data to contain two classes of conditions, which, when taking their difference in neural activity, reflects the studied cognitive process. We define this difference as the *neural signature.* The assumption of operation 2 is this: if the warping signal clocks the cognitive process, then the warped data should show an increase in the periodic dynamics of the neural signature. Thus, we compare the periodicity of a classifier’s performance between clock and brain time data to test for a difference in such patterns. The general pipeline is that each participant’s clock and brain time data undergoes classification (2.3.1), quantification of classifier performance (2.3.2), and first-level statistics (2.3.3). Then, second-level statistics can be initiated on the first-level results (2.3.4).

##### Requirements

To get started with the second operation, both the clock time and brain time warped data need to have each trial’s class membership labelled. This is necessary for *MVPA Light* and *braintime* to understand the class structure present in the data. *Braintime*’s tutorials describe how to do this using example code, but briefly, the user sets `bt_data`. trialinfo and `ct_data`. trialinfo to a vector of 1’s and 2’s in line with the trial structure (class labels). For example, for a dataset on motor processing, a triplet of trials that are in left, right, and left movement conditions would need to be labelled as `[1 2 1]` or `[2 1 2]`.

When using the second operation, it is important for users to establish the absence of high-pass filtering artefacts in the data. In a recent publication, it is reported that high-pass filtering may result in temporal displacement of information, causing leakage of classification performance (van Driel, Olivers & Fahrenfort, 2021). These artefacts may influence the periodicity analysis. In light of this, we suggest avoiding high-pass filtering when the data is clean. Alternatively, trial-masked robust detrending may be applied as an artefact-free method of removing low frequencies (van Driel et al., 2021). Finally, if users do require high-pass filtering, we recommend a careful comparison between classification performance in the filtered data and non-filtered data to establish whether artefacts are present. Users are made aware of potential high-pass filtering artefacts in all sources of *braintime’s* documentation.

###### 2.3.1 Classifying the data

To perform pattern classification, *braintime* calls *MVPA Light* (Treder, 2020). *MVPA Light* can be used to perform multivariate pattern analysis using different models (e.g., support vector machines, logistic regression, and linear discriminant analysis; LDA), classification metrics (e.g., accuracy, area under the curve, fidelity values), and cross-validation methods (e.g., k-fold, hold-out, leave one out). Classification may be performed in the standard way using `mv_classify_across_time` or with temporal generalization using `mv_classify_timextime` (Supplementary Figure 7). The former results in a one-dimensional (1D) time series that reflects how a classifier trained on each timepoint performs when tested on that timepoint in held out data. The latter results in a two-dimensional (2D) temporal generalization matrix (TGM) that reflects how performance generalizes from each trained timepoint to all other timepoints in held out data (King & Dehaene, 2014). The diagonal of the TGM is identical to 1D classification, while the off-diagonal provides information about generalization of performance. This provides information about the degree of constancy of the neural code (“temporal generalization”). As such, `mv_classify_across_time` is appropriate when the neural signature is predicted to evolve beyond classifier recognition from one moment to the next, while `mv_classify_timextime` is appropriate when there is some constancy of the neural code across time. For analysis with *braintime,* if some constancy is expected, we strongly recommend `mv_classify_timextime` as it yields exponentially more data to perform periodicity analysis over – substantially increasing the power of the analysis.

**Supplementary Figure 7:**
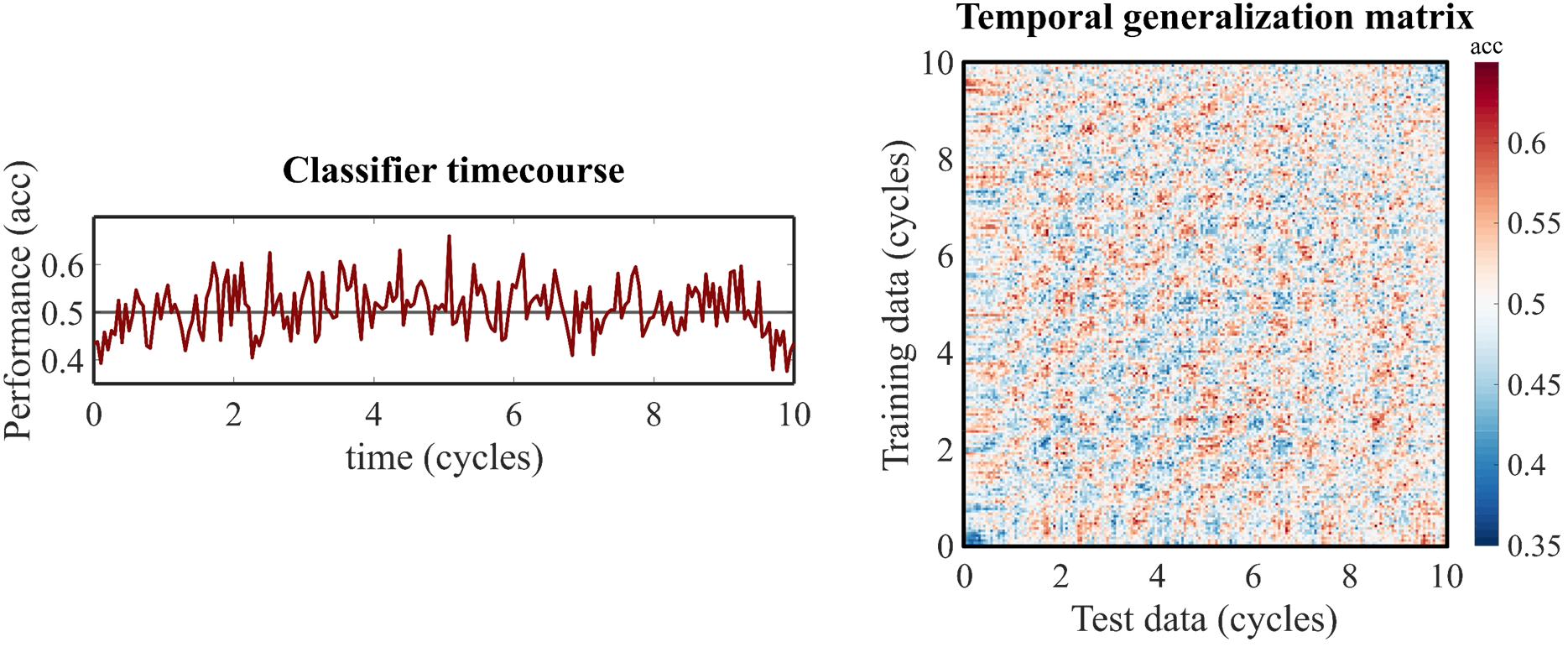
Two types of classification. *Braintime* calls *MVPA Light* to perform standard classification across time (1D; left) or temporal generalization (2D; right). The classification timecourse on the left equals the diagonal of the TGM on the right. The off-diagonal represents how well classification generalizes when trained on one timepoint and generalized to each other timepoint in held-out data. *Braintime* can perform periodicity analyses over both types of classification. *(acc* = accuracy).

###### 2.3.2 Quantifying classifier periodicity

In `bt_quantify`, classifier periodicity is quantified by an FFT over the classifier timecourse (1D results), or over each row and column of the time generalized classifier performance (TGM; 2D results). We use FieldTrip to apply the FFT using a multitaper frequency transformation (using discrete prolate spheroidal sequences and Hanning window tapers). The FFT is performed across frequency ranges specified by the user.

For the time generalized classifier performance, users can specify whether the periodicity is done over the rows and columns of the TGM itself, or over its autocorrelation map (Supplementary Figure 8). The autocorrelation map correlates the TGM with all shifted versions of itself. That is, it iteratively shifts the TGM in each X and Y direction and plots the correlation of the shifted map with itself in coordinate space. The most powerful option depends on the uniformity of periodic patterns. Generally, the autocorrelation map is a more powerful approach, as it accentuates periodic patterns that are present in the TGM. However, when multiple periodicity rates are present in the TGM, the autocorrelation may drown out the weaker periodicity. Similarly, when periodicity is present in only a small part of the TGM, changing its rate or disappearing across time, the autocorrelation map may fail to detect it. In short, when uniform periodicity patterns in the data are expected, analysing the autocorrelation map of TGMs is a powerful approach. When the pattern is expected to only be present partially, or when multiple rates are predicted, it is more powerful to analyse the TGM itself.

###### Periodicity referenced to brain time

For brain time data, just as the time axis has been changed from seconds to cycles, periodicity is redefined from clock time to brain time. Specifically, each participant’s periodicity is normalized to their warping frequency 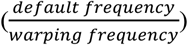. For example, if a user decides to quantify periodicity between 5 and 13 Hz for a participant with a 9 Hz warping frequency, the resulting brain time periodicity will range from between 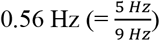 and 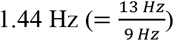, see Supplementary Figure 8 (bottom). This referencing to brain time allows participants to be compared with reference to each brain’s own dynamics. For example, if periodicity across participants were to cluster at 0.5 Hz, this would mean participants tend to show periodic patterns in their neural signature at half the rate of their clocking oscillations.

###### 2.3.3 First-level statistics of periodicity

In `bt_statslevel1`, quantified periodicity is subjected to first-level statistics to establish its statistical robustness. The user specifies a range of frequencies that will be tested, and the desired number of first-level permutations (`numperms1`). *Braintime* obtains a null distribution by repeating data classification (2.3.1) and quantification of its periodicity (2.3.2) `numperms1` times with the class labels shuffled randomly (destroying correspondence between trials and their associated condition). This yields a pool of *permuted periodicity spectra* that offers a confidence interval for each frequency. P-values are not provided until the second-level statistics are completed.

**Supplementary Figure 8:**
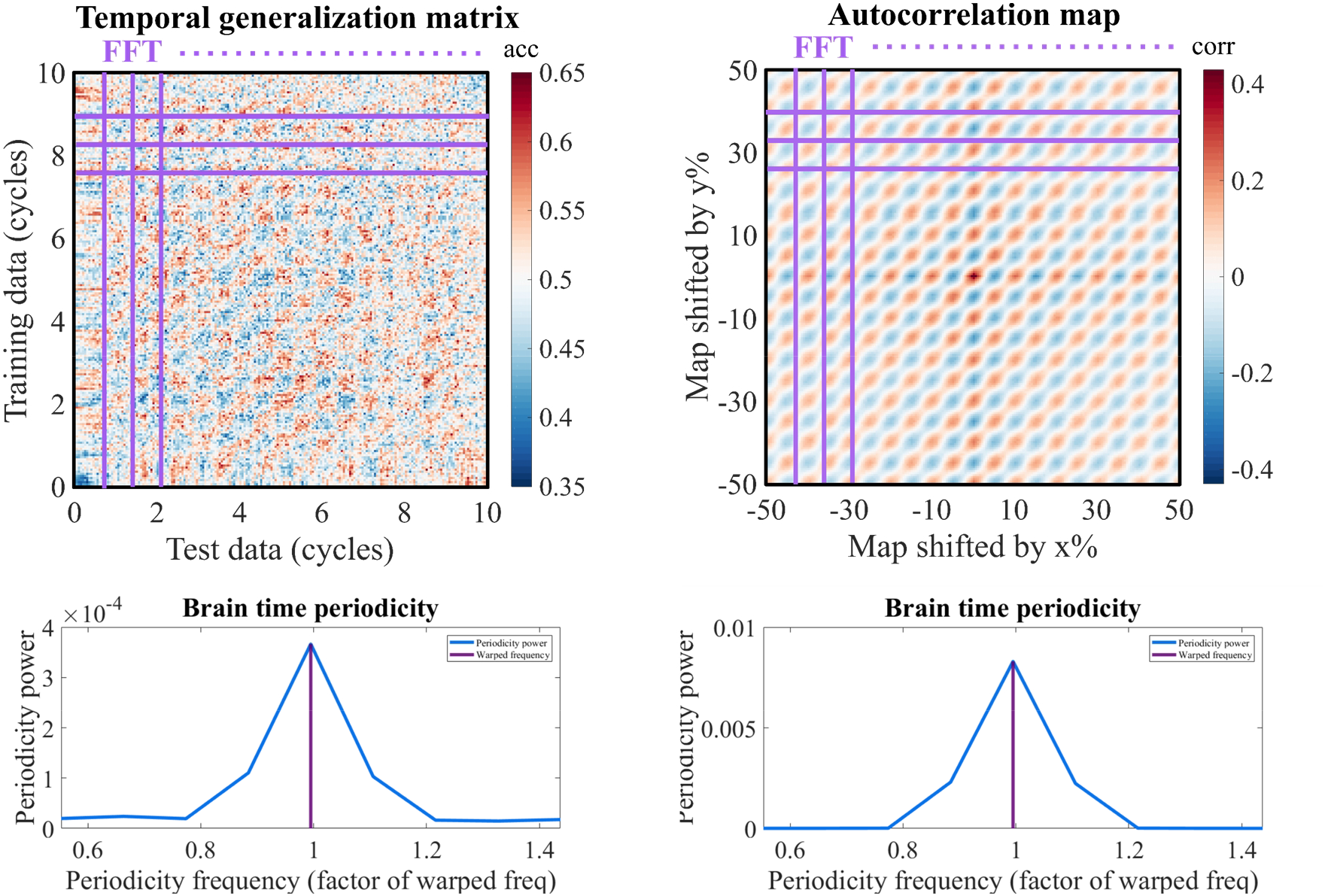
Two ways of performing 2D periodicity analysis. **(Top)** Row-by-row and column-by-column, *braintime* performs an FFT over the TGM by itself, or its autocorrelation map. The former is more powerful when periodicity is present in small segments of the TGM, or if multiple periodicity frequencies are present. The latter is more powerful when the periodic pattern is present in most of the TGM (as is the case in this dataset). **(Bottom)** The resulting power spectra are averaged across all rows and columns, resulting in one participant’s “periodicity spectrum”. *(corr* = correlation)

###### 2.3.4 Second-level statistics of periodicity

Upon completion of the previous steps for all participants, users are requested to combine the first-level periodicity data structure of each participant into one superstructure. Next, in anticipation of `bt_statslevel2`, users specify the number of second-level permutations (`numperms2`) and the desired method of multiple testing correction across periodicity frequencies (false discovery rate; FDR, Bonferroni correction, or no correction). `bt_statslevel2` undertakes the following steps:

###### Z-scoring

*Braintime* performs z-scoring on each participant’s periodicity spectra for two reasons. First, the output of FFT has amplitudes with arbitrary values, creating arbitrary differences in amplitude between participants. Second, we want to avoid participants with high periodicity (even after correcting for arbitrary differences) to drive a group effect. The toolbox implements z-scoring by normalizing both the permuted spectra and empirical spectra to the mean and standard deviation of the permuted spectra (i.e., by subtracting the mean and dividing by the standard deviation). With this method, the permuted spectra are considered a noise floor for which the empirical spectra are corrected.

###### Filtering to common frequencies

As discussed before (2.3.2), participants’ range of tested periodicity frequencies are normalized to their warping frequency. This creates some variability in the range of tested frequencies across participants. For second-level statistics, *braintime* filters each participant’s spectra to the frequencies that are common to all participants. To overcome potential variations in the frequency resolution between participants, *braintime* uses `imresize` to interpolate all participants’ spectra according to the mode spectral resolution across participants.

###### Obtaining p-values

On the second-level, each clock and brain time periodicity frequency is statistically tested across participants. This is done in the following way. For each participant, the toolbox grabs one random spectrum of each participant’s `numperms1` permuted spectra. This is repeated `numperms2` times, yielding a pool of `numperms2` new spectra. For each frequency, *braintime* compares the average empirical periodicity power to the distribution of power in the `numperms2` pool spectra. The p-value of each frequency is defined as the proportion of `numperms2` power values that are equal or higher to the average empirical power value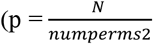, where N is the number of times the permuted pool equals or exceeds empirical power).

###### Multiple testing correction

Users may correct for multiple comparisons (where the comparisons are the tested frequencies) by using false discovery rate (FDR) or Bonferroni correction. For brain time spectra, three frequencies are exempted by default for multiple testing correction because, in line with the hypothesis-driven approach of *braintime,* periodicity is a priori expected at these rates. Specifically, these are at 0.5×, 1×, and 2× the warping frequency (i.e., 0.5 Hz, 1 Hz, and 2 Hz) in the brain time spectra. Depending on both the underlying dynamics and the classifier method used (`mv_classify_across_time` or `mv_classify_timextime`), periodicity is predicted at one or more of these rates (if the warping signal clocks the cognitive process of interest). In this sense, *braintime* performs two statistical tests: one primary test across the three hypothesized frequencies, and an additional one across the specified frequency range of interest. In the second-level periodicity spectra, the uncorrected p-values at the three frequencies of interest are combined with the corrected p-values of the remaining frequencies of interest.

###### Cluster statistics

Besides establishing the statistical robustness of periodicity in the classifier’s performance, *braintime* also displays robustness of classifier performance per se. That is, it displays at which timepoints the classifier is able to differentiate the two classes of data above chance. The toolbox calls *MVPA Light* to perform cluster correction over the results obtained from `mv_classify_across_time` or `mv_classify_timextime`. Users may specify a variety of parameters for cluster correction that are read and implemented by *MVPA Light,* including test methods (e.g., binomial or permutation based-cluster correction) as well as the number of permutations used to determine significant clusters. *MVPA Light* implements cluster correction as described by Maris & Oostenveld (2007). More information on cluster correction in *MVPA Light* can be found on its Github page.

###### 2.4 Methodological considerations

Here, we describe several methodological concerns and considerations when using *braintime*.

###### 2.4.1 Circularity concerns

As mentioned in the main text, a concern with brain time warping could be that it trivially imposes oscillatory structure at the warping frequency, causing either the activity or neural signature of a cognitive process to fluctuate even when warping to oscillations that do not clock it. A first point is that the warping path generated by DTW is not oscillatory, but instead represents how the warping signal needs to be transformed to reduce its non-stationarity. That is to say, application of the warping path to a random signal is not expected to alter its oscillatory structure in any way. Nevertheless, the concern persists. If the warping signal is present in the data, the oscillatory structure of the data is likely enhanced because the warping signal has its non-stationarities removed as a consequence of the path being applied to all the data. Below, we describe for which analyses circularity could be a concern, and how it can be avoided.

###### Analysis

Circularity is not a concern when testing for periodic patterns using the toolbox (operation 2). This is because of *braintime’s* method of obtaining a null distribution against which empirical periodicity is tested. Specifically, the only difference between this permuted pool and the empirical data is the shuffling of class labels. Besides this difference in data-to-condition allocation, the trials themselves remain unchanged. As a consequence, any trivially imposed oscillatory structure into the data is present equally between the permuted and empirical set. To the extent a classifier is able to exploit circular patterns (if they are present), it will be able to do so for both dimensions. Thus, where circularity is present, it is expected to cancel out through *braintime’s* statistical method.

However, when using brain time warped data outside the toolbox, it may be that a researcher’s analyses tap into the circular patterns (if they are present). Following the same logic, circularity is an issue to the extent it differentially affects the empirical and null distribution.

###### Dependence

Since circularity may arise from the warping signal, whether circularity is present at all critically depends on whether the warping signal is present in the transformed data. We call this concept data dependence. If the warping signal is present in the transformed data, the transformed data are said to be dependent. Below, we provide several examples with varying levels of dependence. Most caution and vigilance is needed for highly dependent data. To emphasize, the level of dependence is only relevant when analysing brain time warped data outside *braintime*, and only with analyses that can exploit circular patterns.

*High dependence*

H1) The warping signal was obtained from a channel in the clock time data, and it is still present in the brain time data.
H2) The warping signal was obtained from an ICA component in the clock time data, and it is still present in the brain time data.

*Medium dependence*

M1) The warping signal was obtained from source localized virtual channels obtained from within the clock time data, and these channels are still present in the brain time data.
M2) The warping signal comes from a set of anterior channels of an EEG or MEG dataset, which was used to warp posterior sensors.
M3) The warping signal is from high pass filtered MEG data, which is used to warp low pass filtered data.

*Independence*

These include all cases where the signal used for warping is absent from the brain time data. For example:

L1) The warping signal is obtained from local field potentials (LFP) in multi-unit recordings, which is used to warp single-cell recordings.
L2) The warping signal is from an ICA component that is absent in the brain time data.
L3) The warping signal is from a channel that is absent and spectrally orthogonal to channels in brain time data.

###### Solution

Circularity can be minimized by removing the warping source from the brain time warped data. An obvious example is the case where the warping source structure comprises ICA components (H2). *Braintime* automatically detects when warping sources are ICA components, and both recommends their removal and assists in this process (H2 → L2). Similarly, most of the aforementioned examples can be made independent. For example, in example M1, the spatial filters used for virtual channel generation could be used to filter those channels from brain time warped data. Or, if a brain time warped data structure comprises both LFP and single-unit channels, and the LFP channels were used to warp, then these channels can be removed from the brain time warped data.

###### 2.4.2 Warping artefacts

Sometimes, brain time warping introduces a long series of sample repetitions at the start and/or end of the data that are too extended for the cycle-by-cycle resizing operation to compress them (Supplementary Figure 9). These extended data repetitions tend to occur at the start of brain time warped data, because this is where DTW attempts to repair initial differences in the phase between clock time and the warping signal. In other words, typically clock and brain time start out of tune, and so brain time warping repeats brain time data from the get-go until the two dimensions fall back in tune. These data repetitions are not intrinsically artefactual – they represent a true mismatch between the brain’s dynamics and clock time that brain time warping accounts for. Nevertheless, they could in principle affect subsequent analyses, such as *braintime’s* periodicity analysis, causing them to exert an artefactual low-frequency footprint on it. For simplicity, we therefore refer to these data repetitions as warping artefacts.

Generally, warping artefacts are too small to exert an effect on *braintime’s* periodicity analysis. Their effects on analyses outside the toolbox depends on the degree to which the data repetitions affect it. The toolbox has built-in tools to avoid the artefact, if so desired. Specifically, the toolbox has a “cutartefact” parameter that brain time warps an additional 0.5 seconds before and after the specified window of interest, which is subsequently removed at the end of `bt_clocktobrain`. Here, the data repetitions at the start (end) of the brain time warped data are shifted to the 0.5 seconds before (after) the window of interest. The advantage of this method is that the warping artefact is cut. The downside of this method is that there is a slight mismatch across trials in the true and specified time window of interest in the order of several milliseconds (for this reason, the default method that implements no artefact cutting is called “consistenttime”). Researchers may decide on a case-by-case basis which method is appropriate – cutartefact or consisttenttime – depending on the relative importance of the absence of warping artefacts and the constancy of the data’s timing.

The two methods warp over different time periods and may therefore vary in their determination of which data belong to which cycle (the cycle-by-cycle resizing step, see 2.2.4). *Braintime* has a special function (`bt_checkallocation`) that visualizes how each of the two methods allocates data to cycles, allowing users to make informed decisions about which method to use and how much they diverge.

**Supplementary Figure 9:**
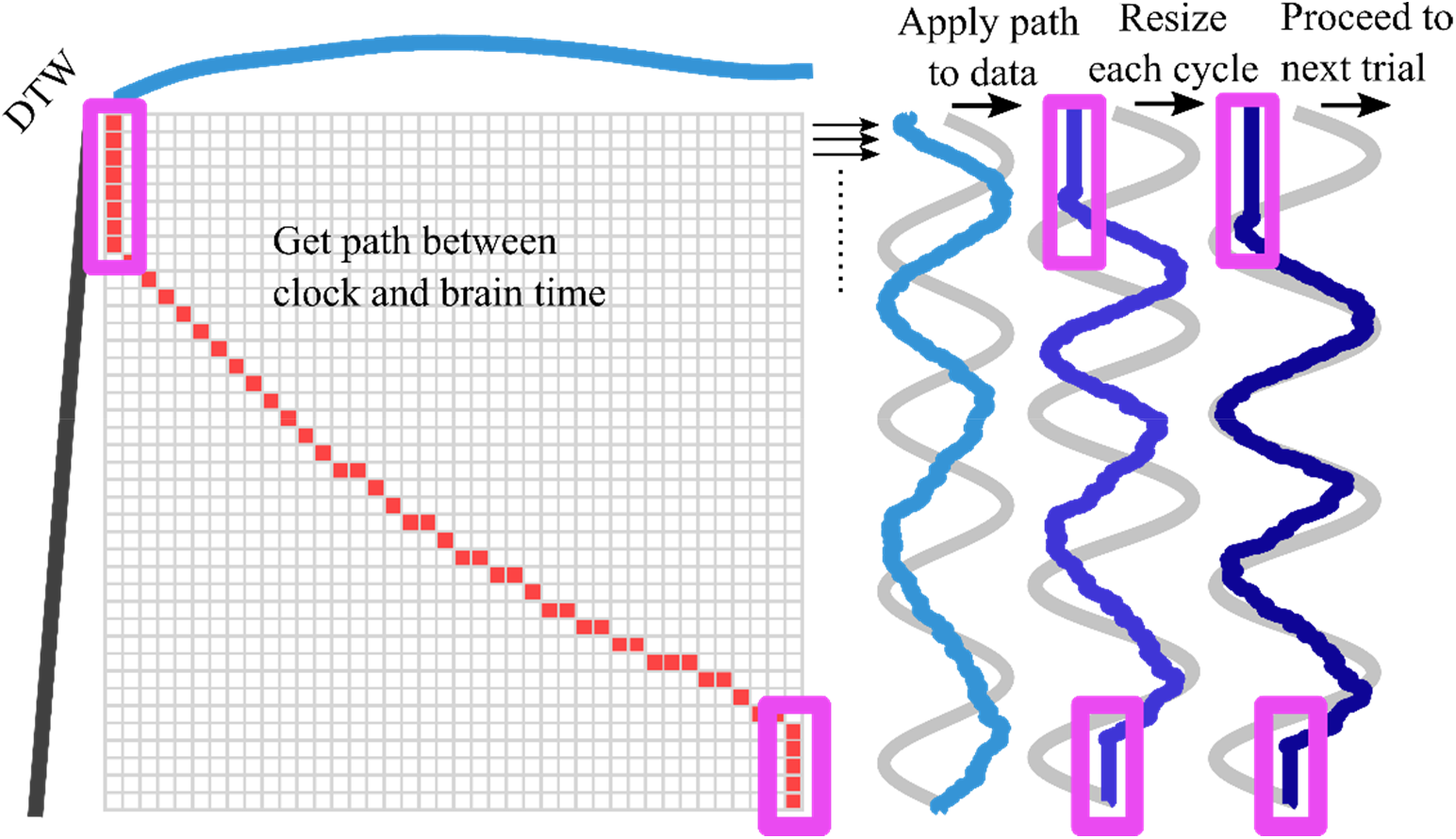
Warping artefacts. Brain time warping may repeat samples for an extended period at the start and end of the data, depending on the disharmony between clock and brain time at the onset of trials (purple boxes). *Braintime* includes a method to remove the artefact, which warps additional data before and after the window of interest which is subsequently removed.

#### Supplementary Results

For the plots of each dataset analysis, the y-axis of periodicity spectra and the colour scale of TGMs and autocorrelation maps are fixed to a constant range across (1) participants and (2) clock and brain time results to enable a visual comparison of magnitude. For a description of each type of classification plot, see section 2.3.

##### 3.1 Simulation

###### 3.1.1 ERP difference

To test whether the simulation yielded a fluctuating signature, we analysed the difference in average activity (event-related potentials; ERP) between left and right hemifield conditions. When using brain time warping to overcome non-stationarities, the simulated periodic neural signature becomes unveiled.

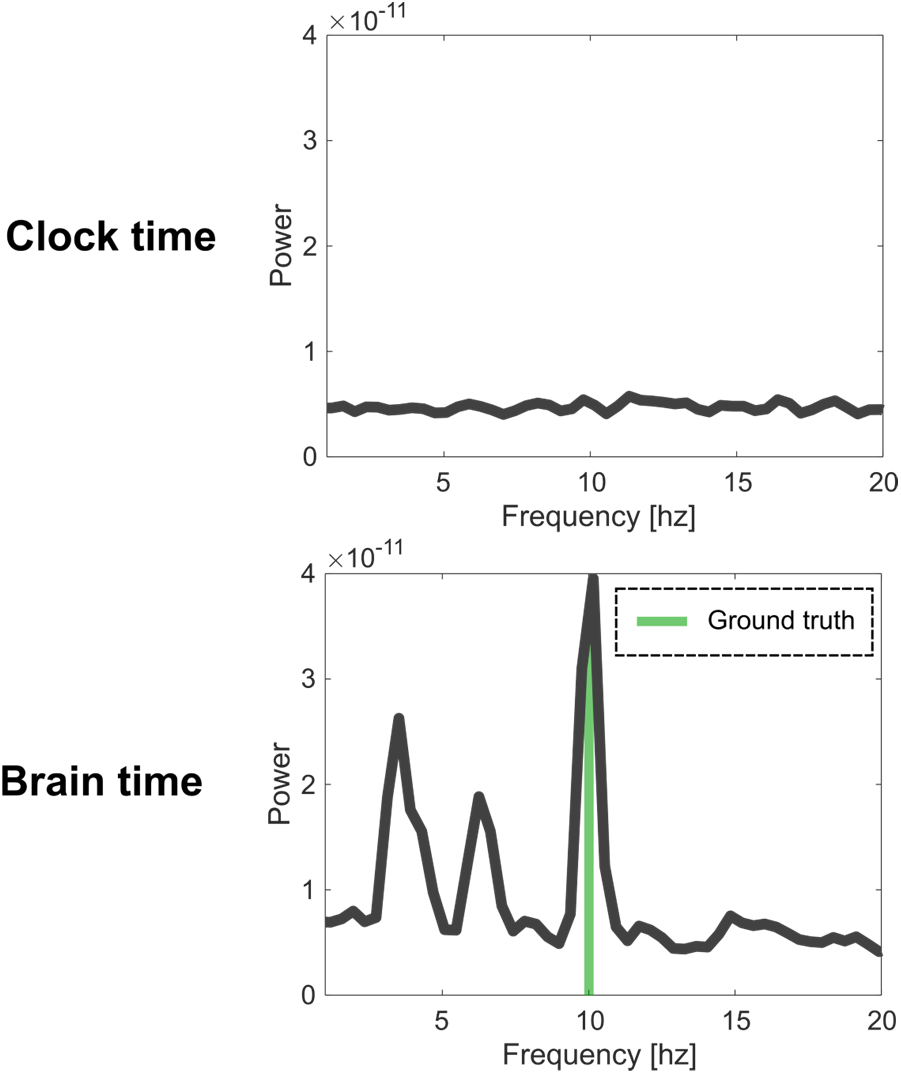

###### 3.1.2 Advanced (main analysis)

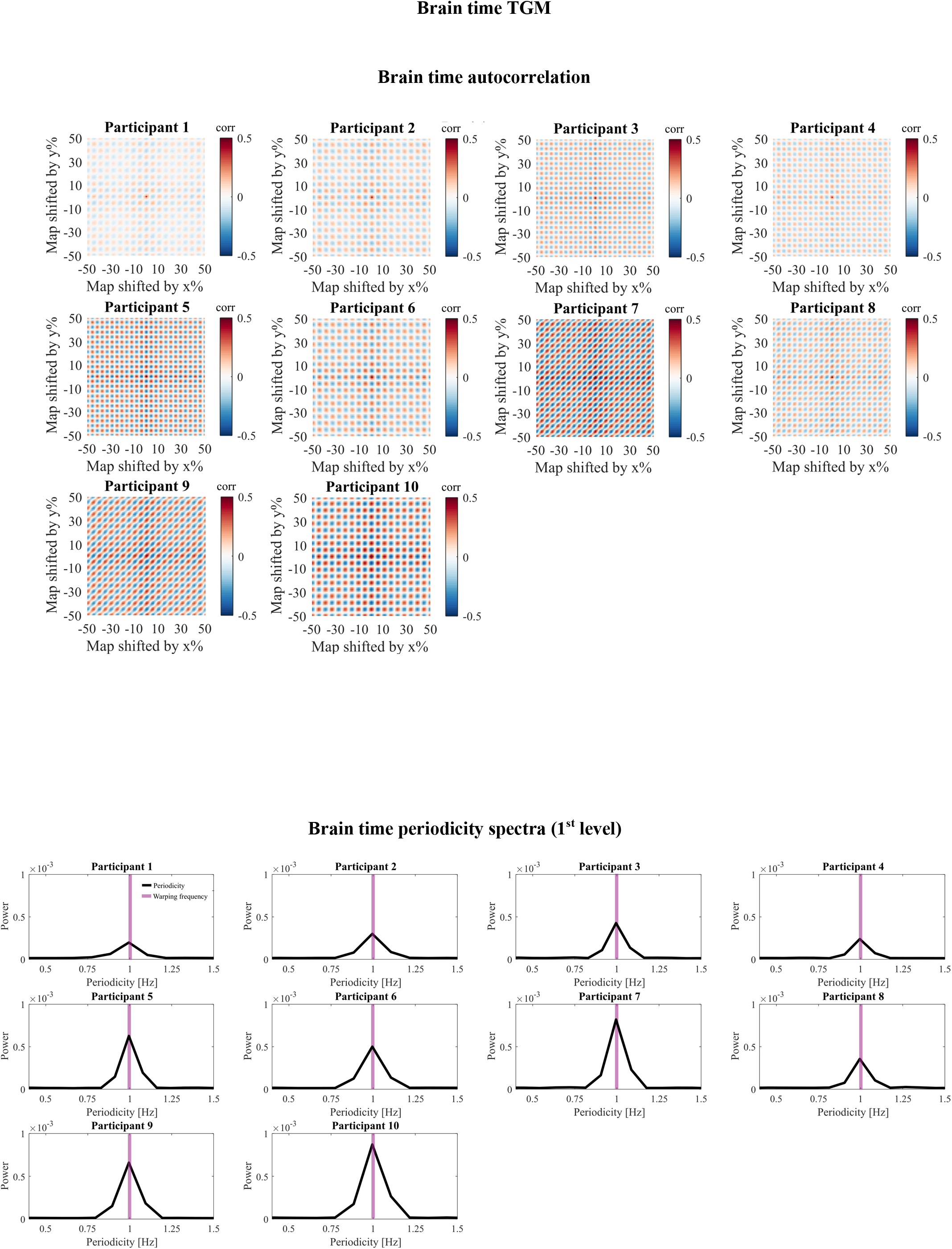

###### Brain time clusters

For the simulated dataset, we found few significant clusters with an alpha set at 0.05 and a critical t-value set at 1.96. We interpret these findings as indicating that the difference in simulated activity between left and right hemifield conditions is too small to be robustly detected by the classifier. However, since we know there is a difference in the ground truth between classes, we know this lack of significant results reflects a type II error. The fact that the periodicity analysis does show significant results indicates that, between empirical and permuted data, differences in the fluctuations of the neural signature are easier to detect than a mean baseline shift in signature fidelity.

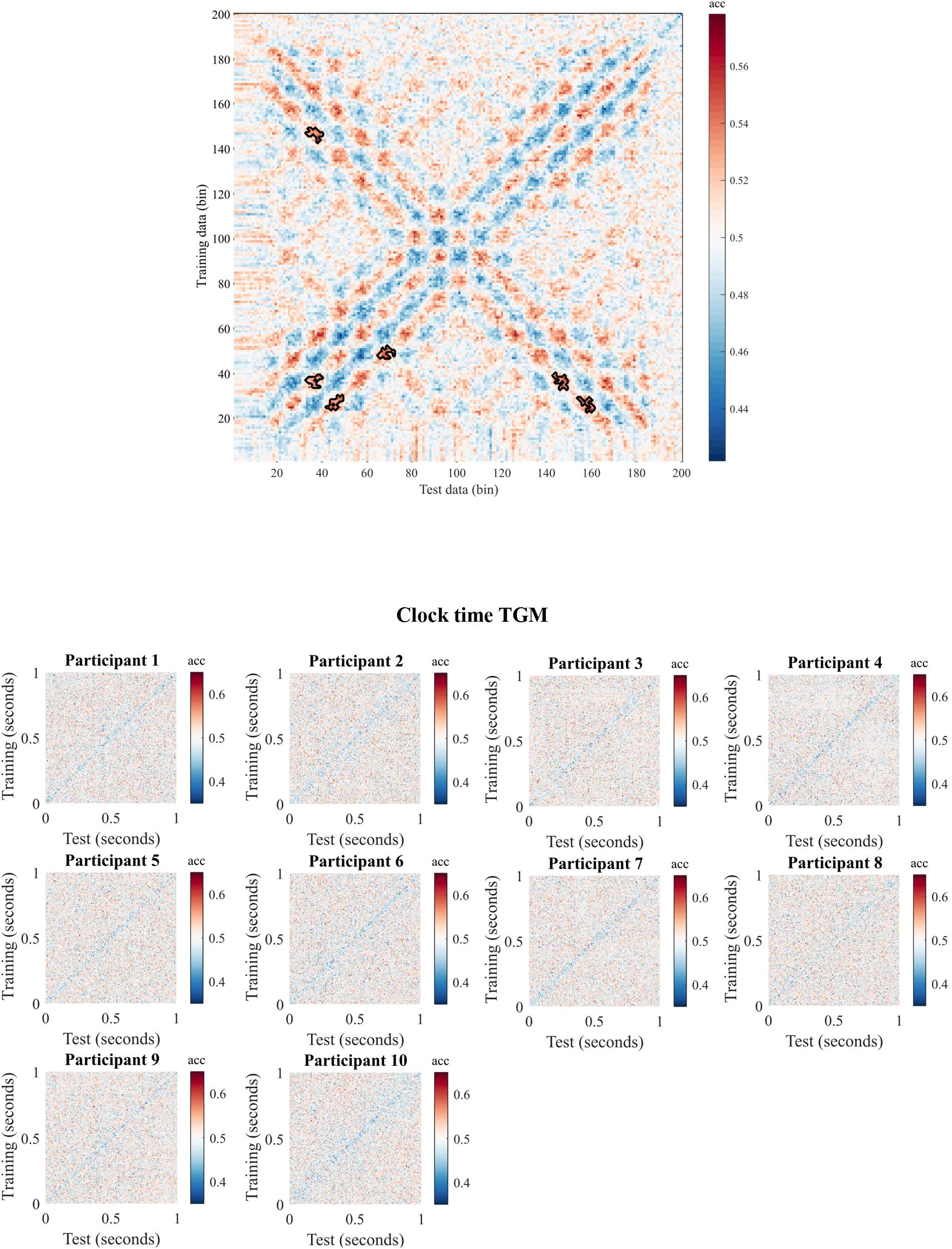

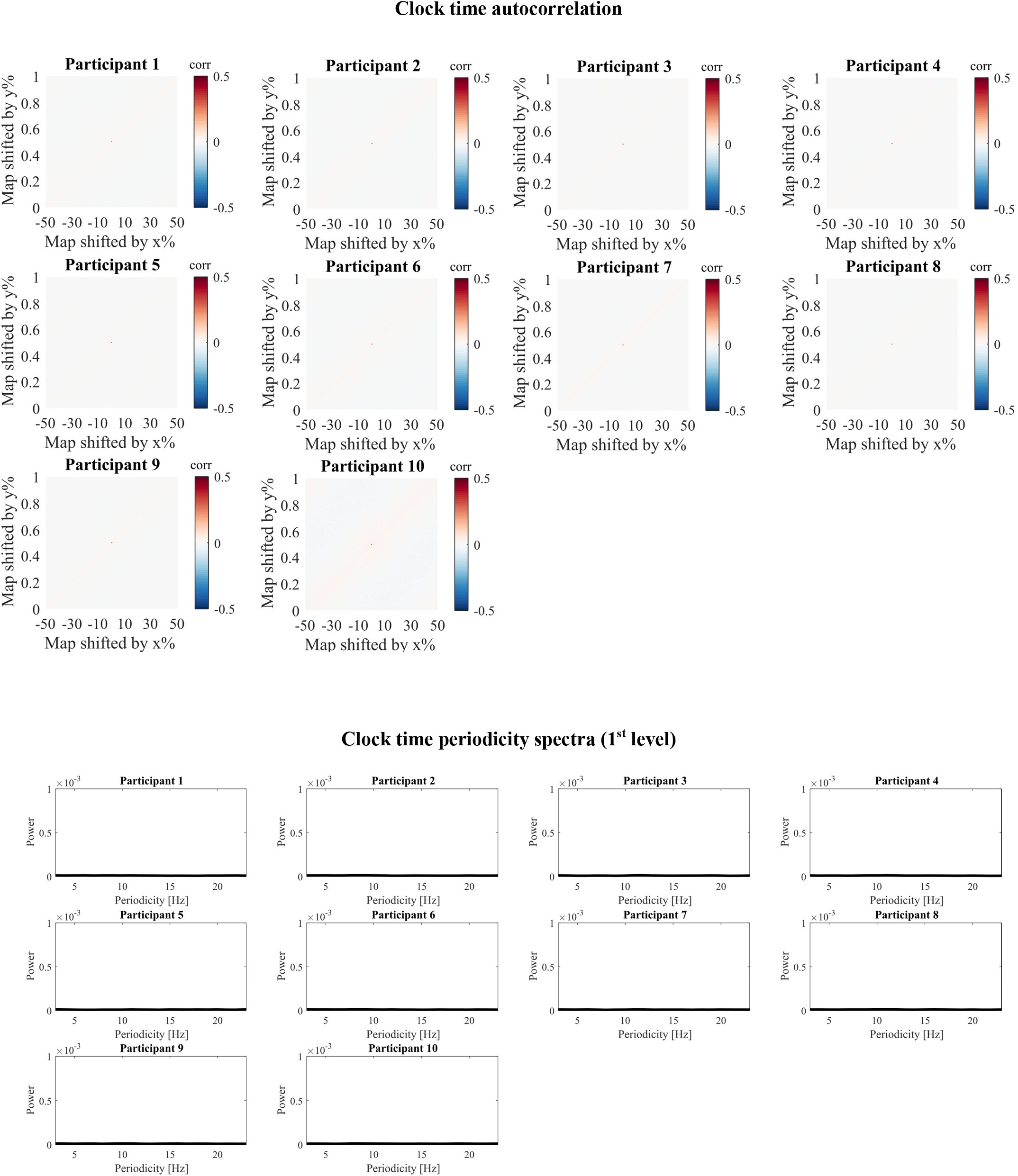

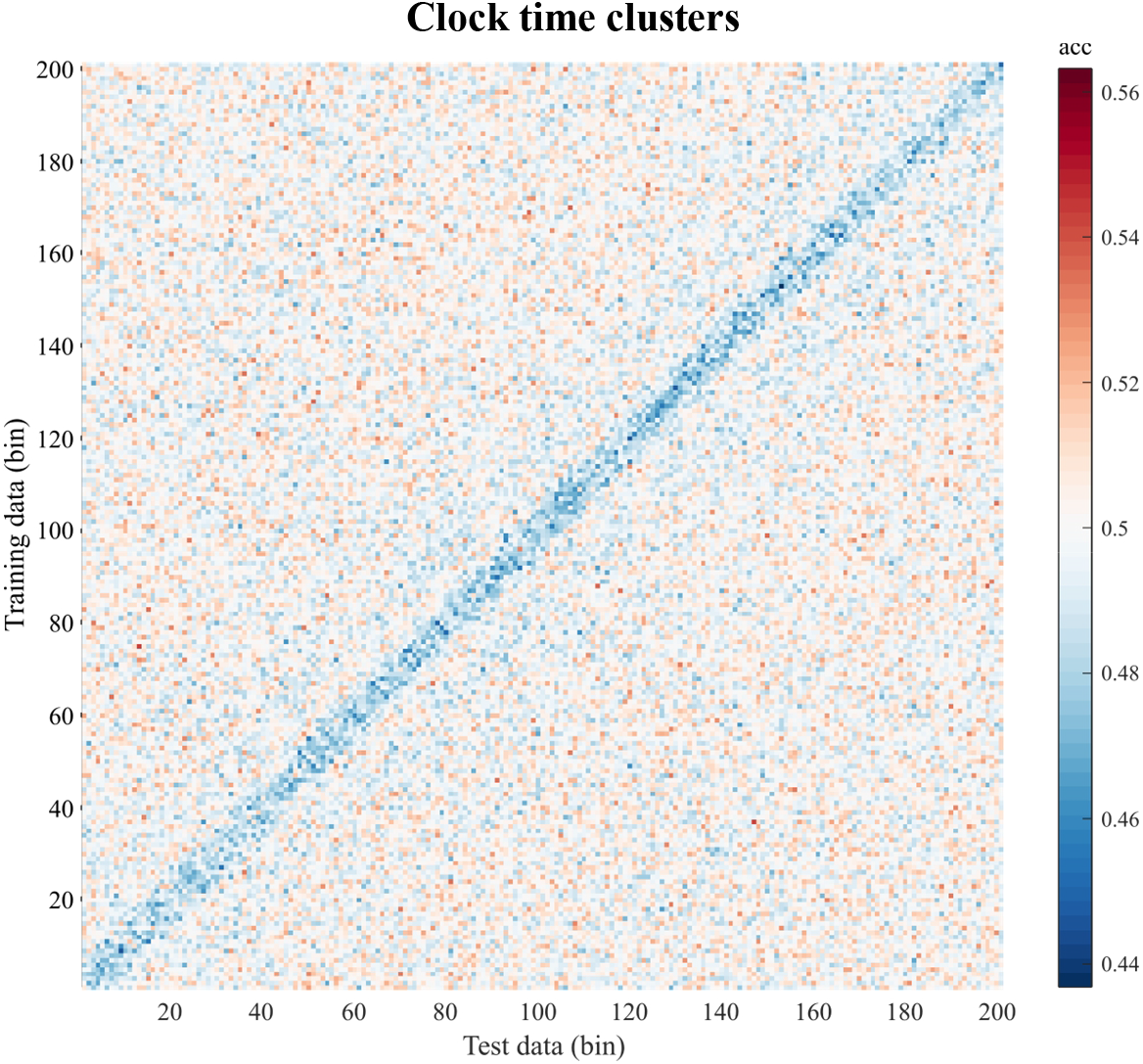

###### 3.1.3 Advanced (control analysis)

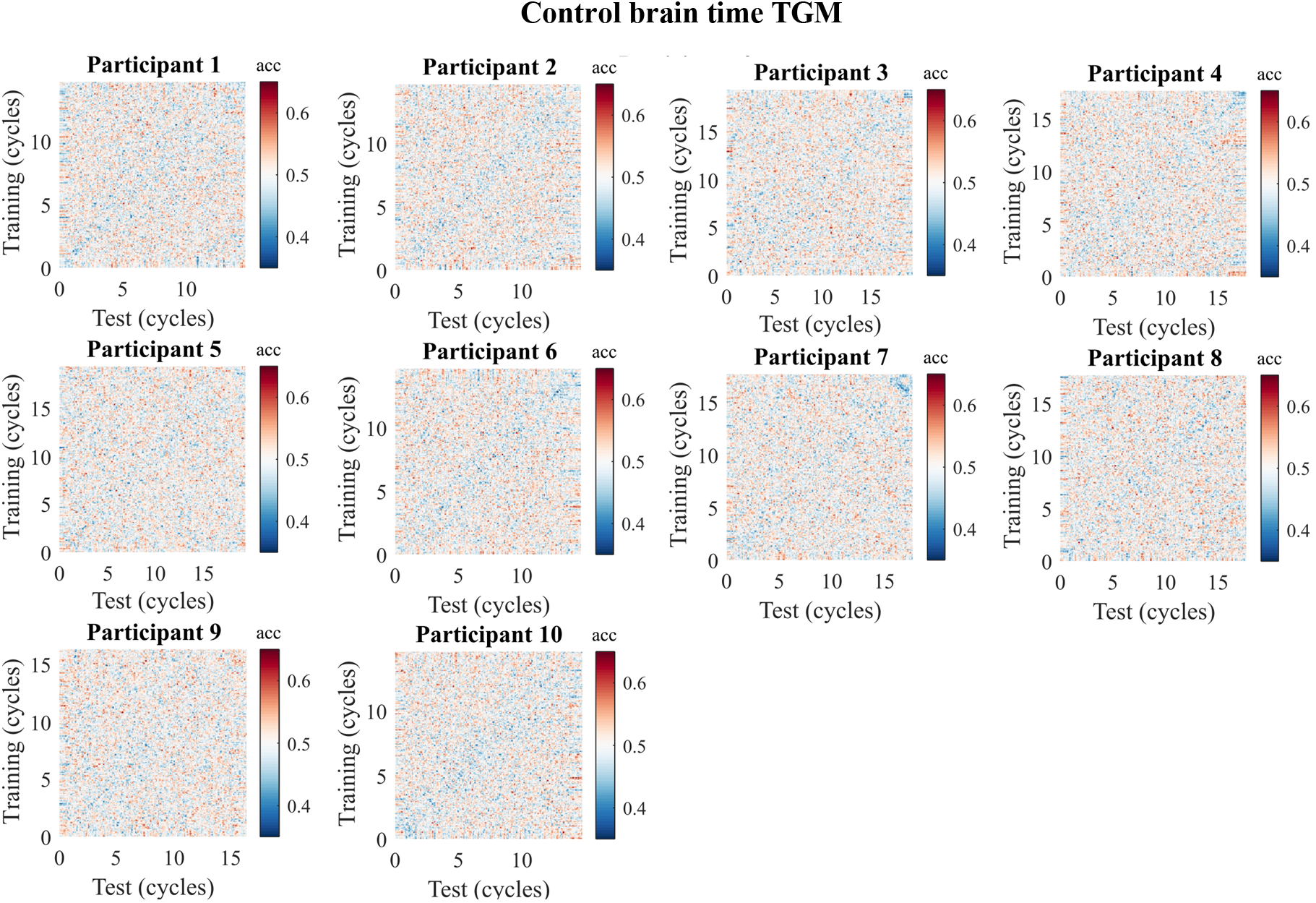

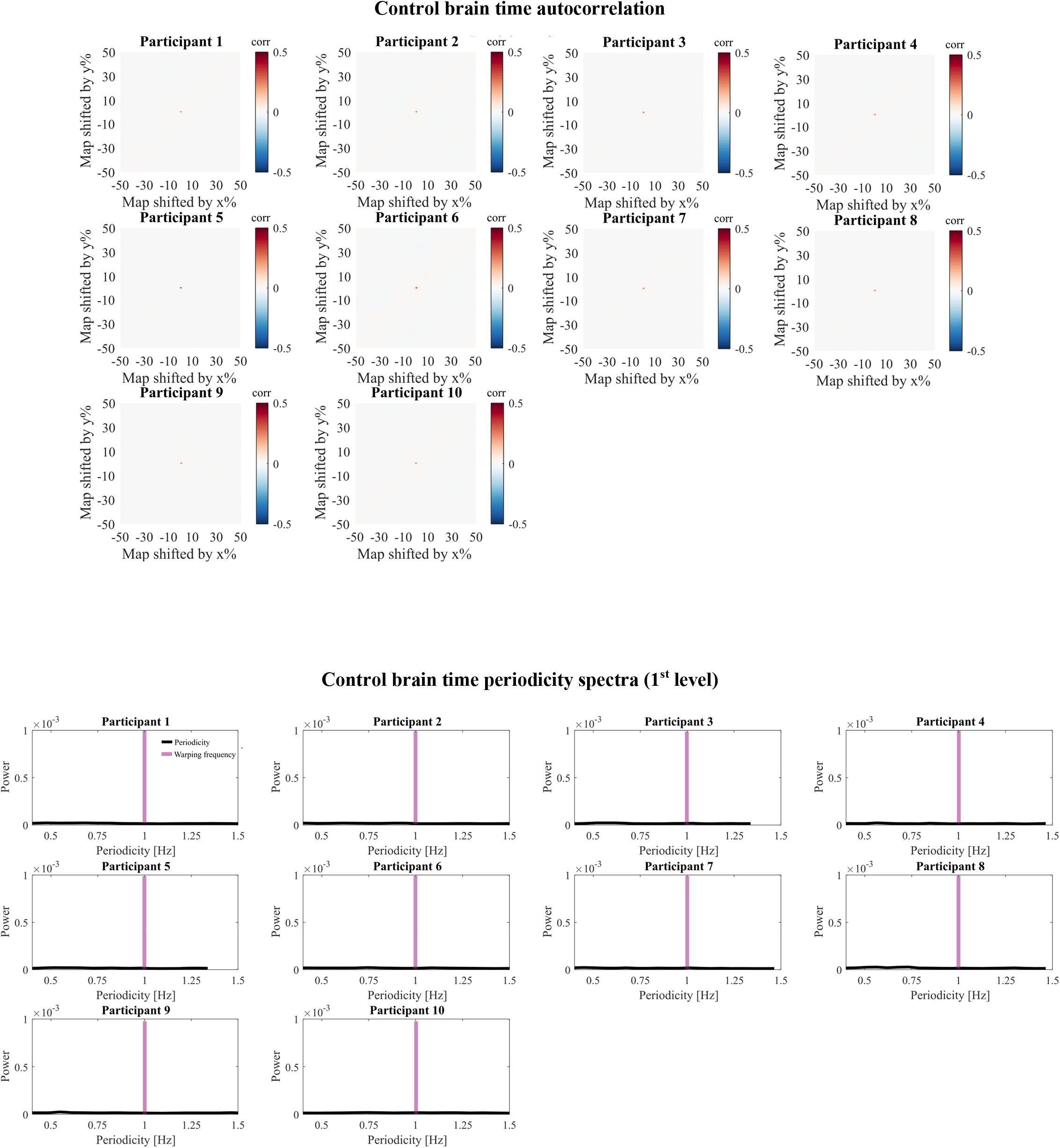

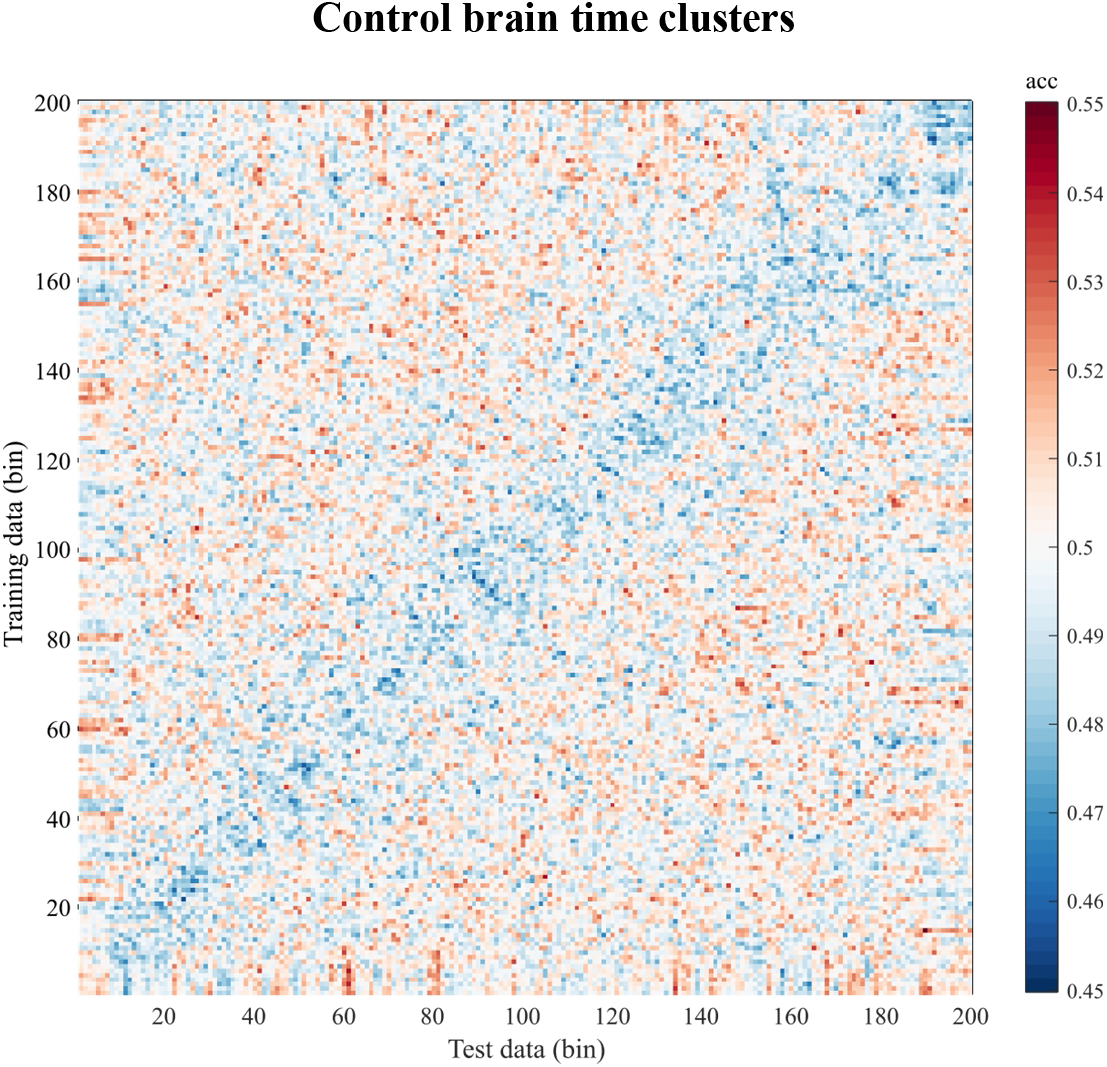

##### 3.2 Rodent

###### 3.2.1 Basic (main analysis)

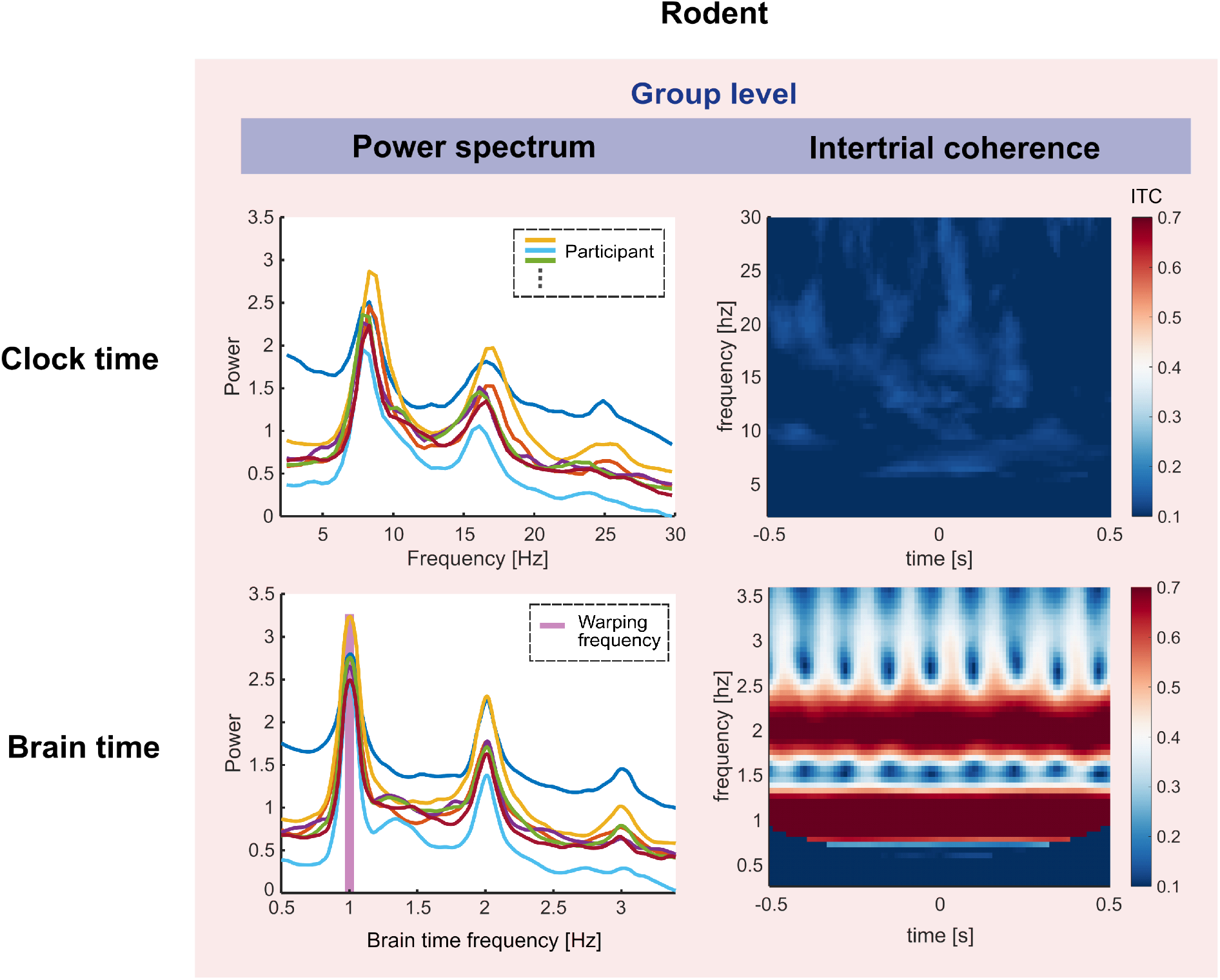

###### 3.2.2 Grid cells

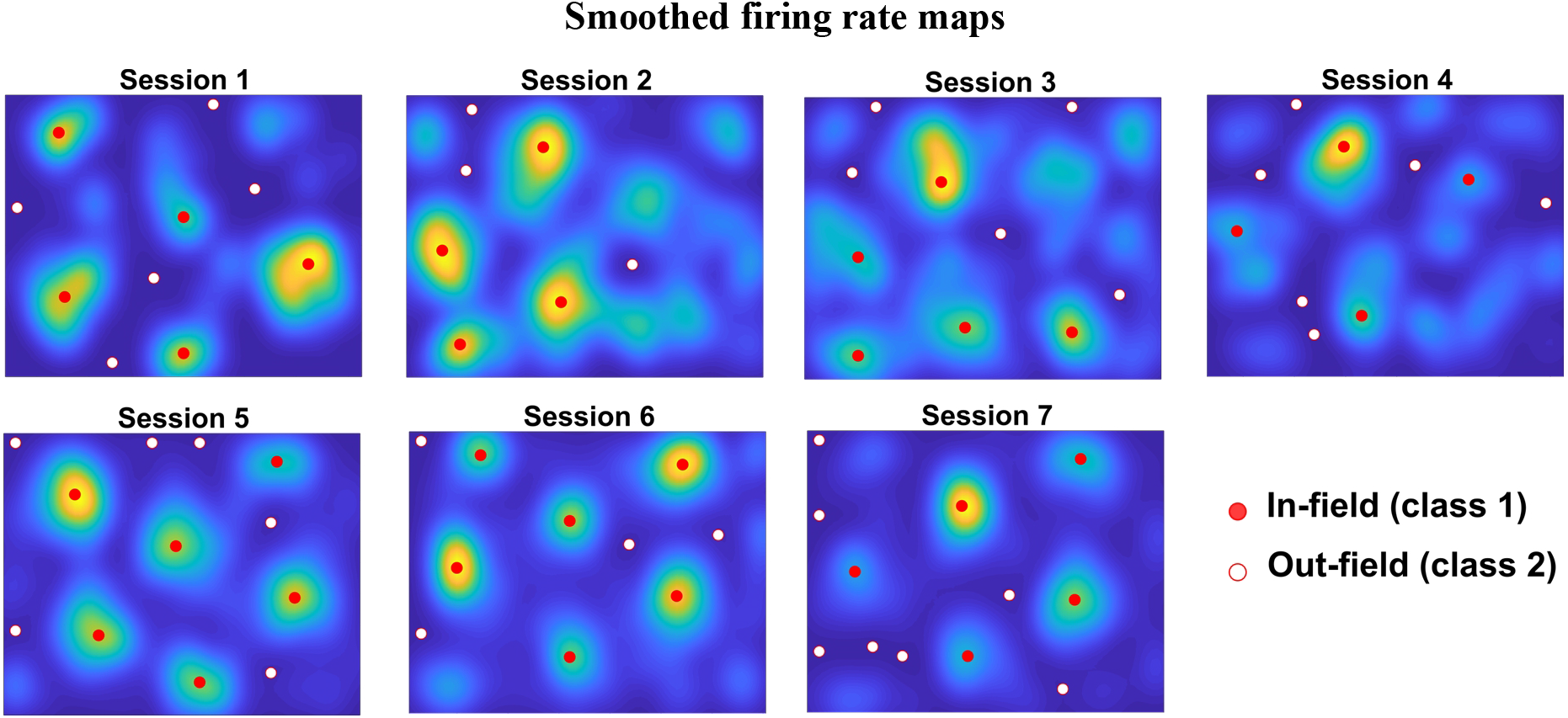

###### 3.2.3 Spike-field coupling

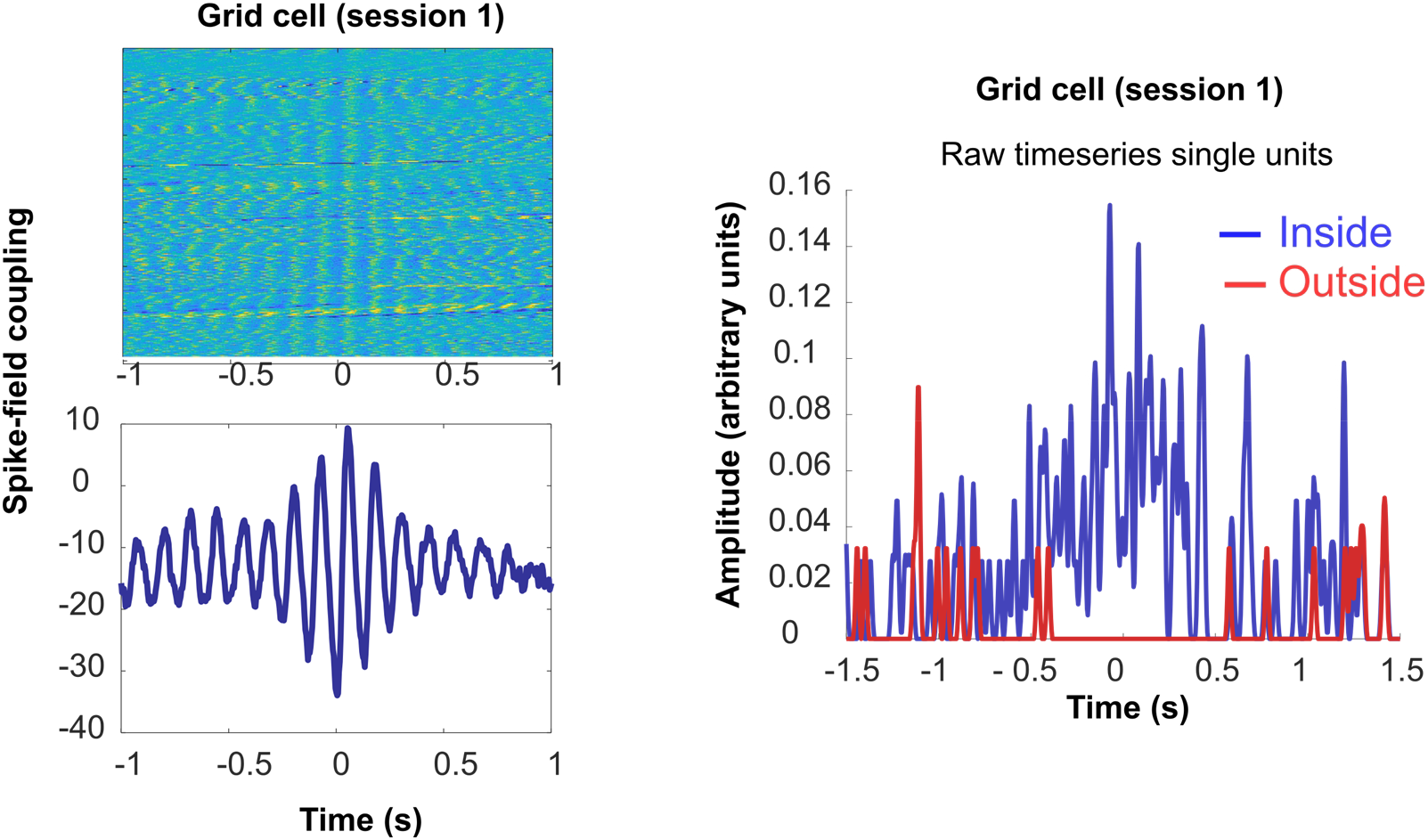

###### 3.2.4 Advanced

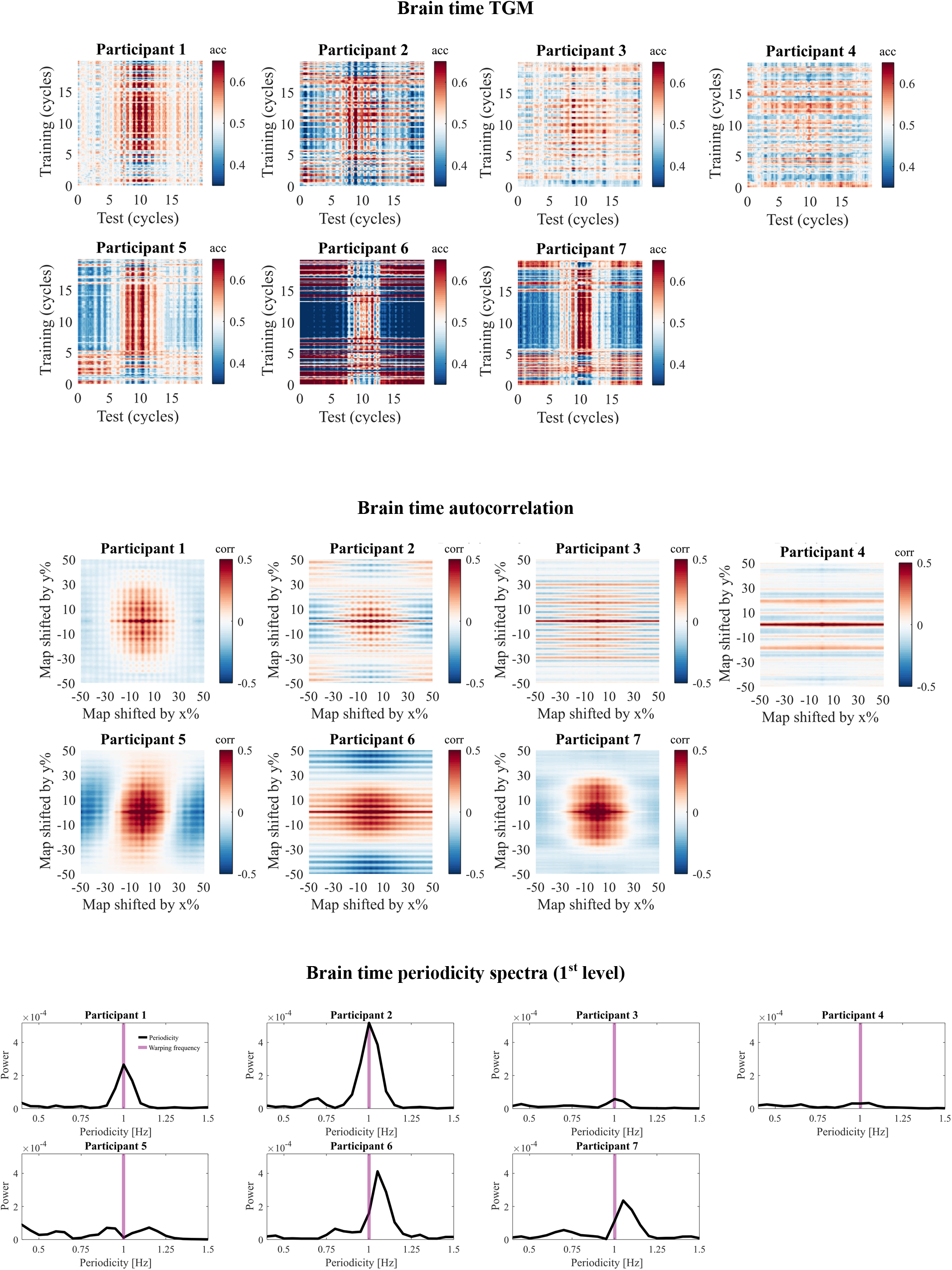

###### Brain time clusters

In line with our prediction, significant clusters occur in the period around t = 0, where the data was defined to contain a class difference.

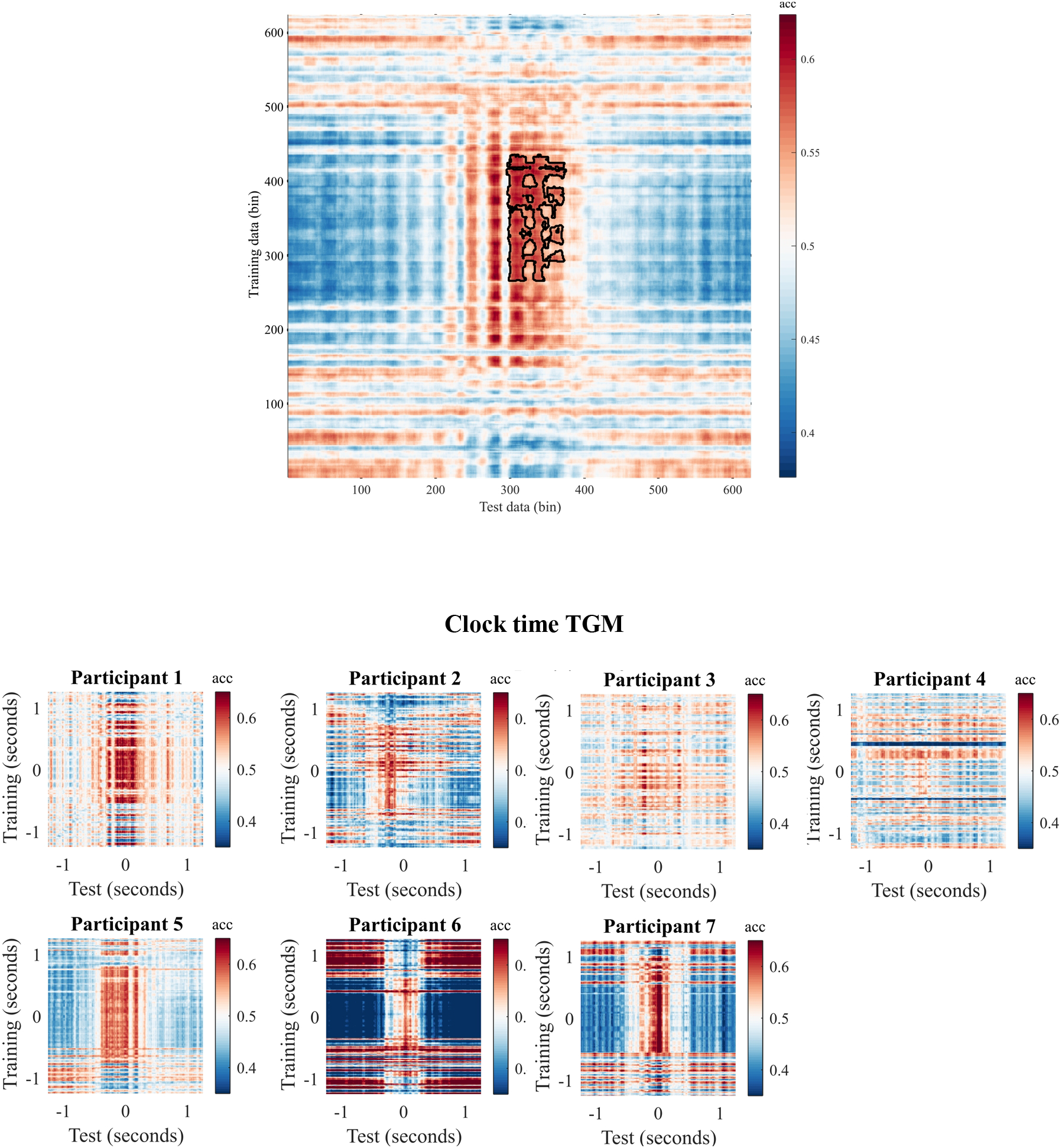

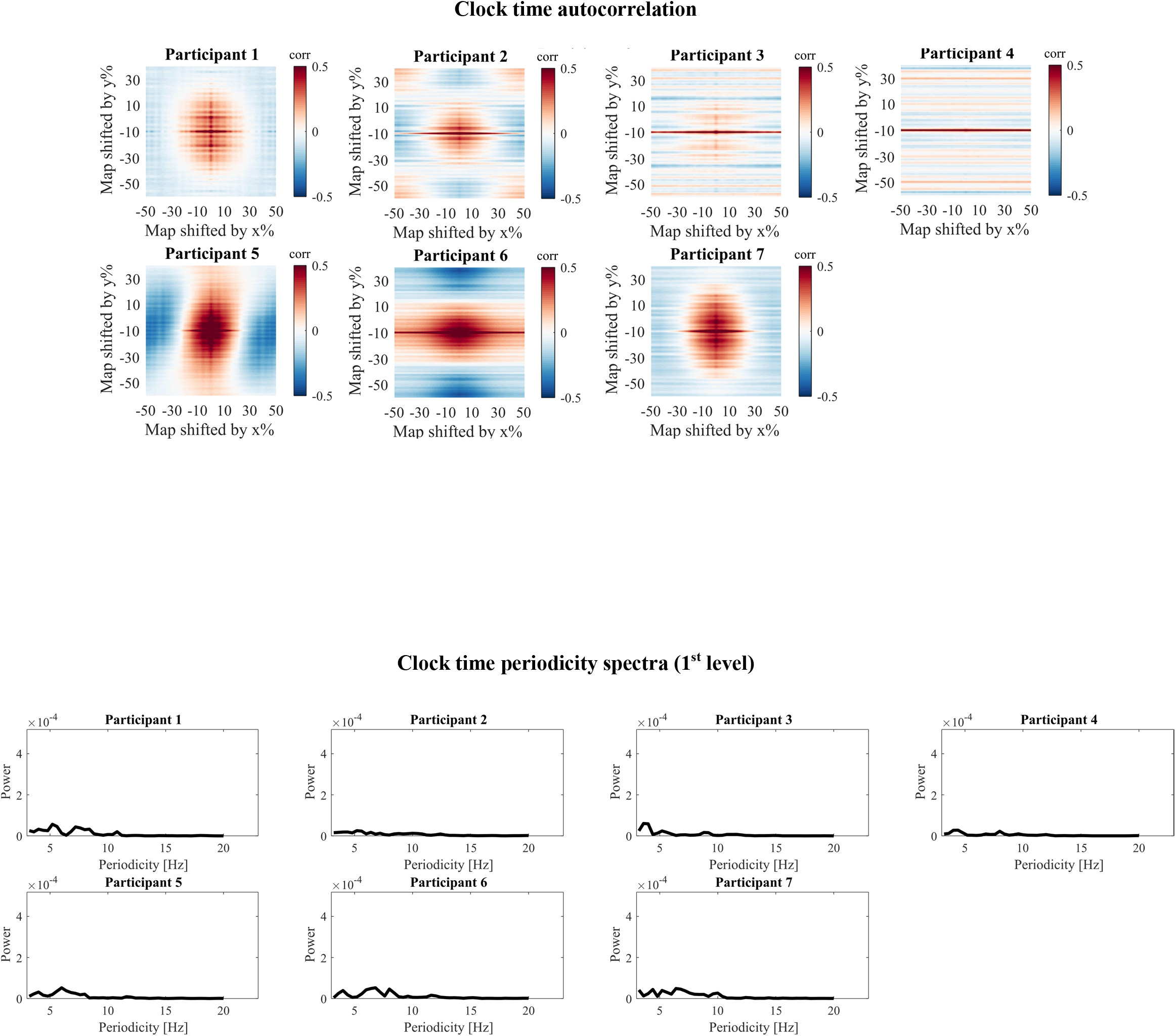

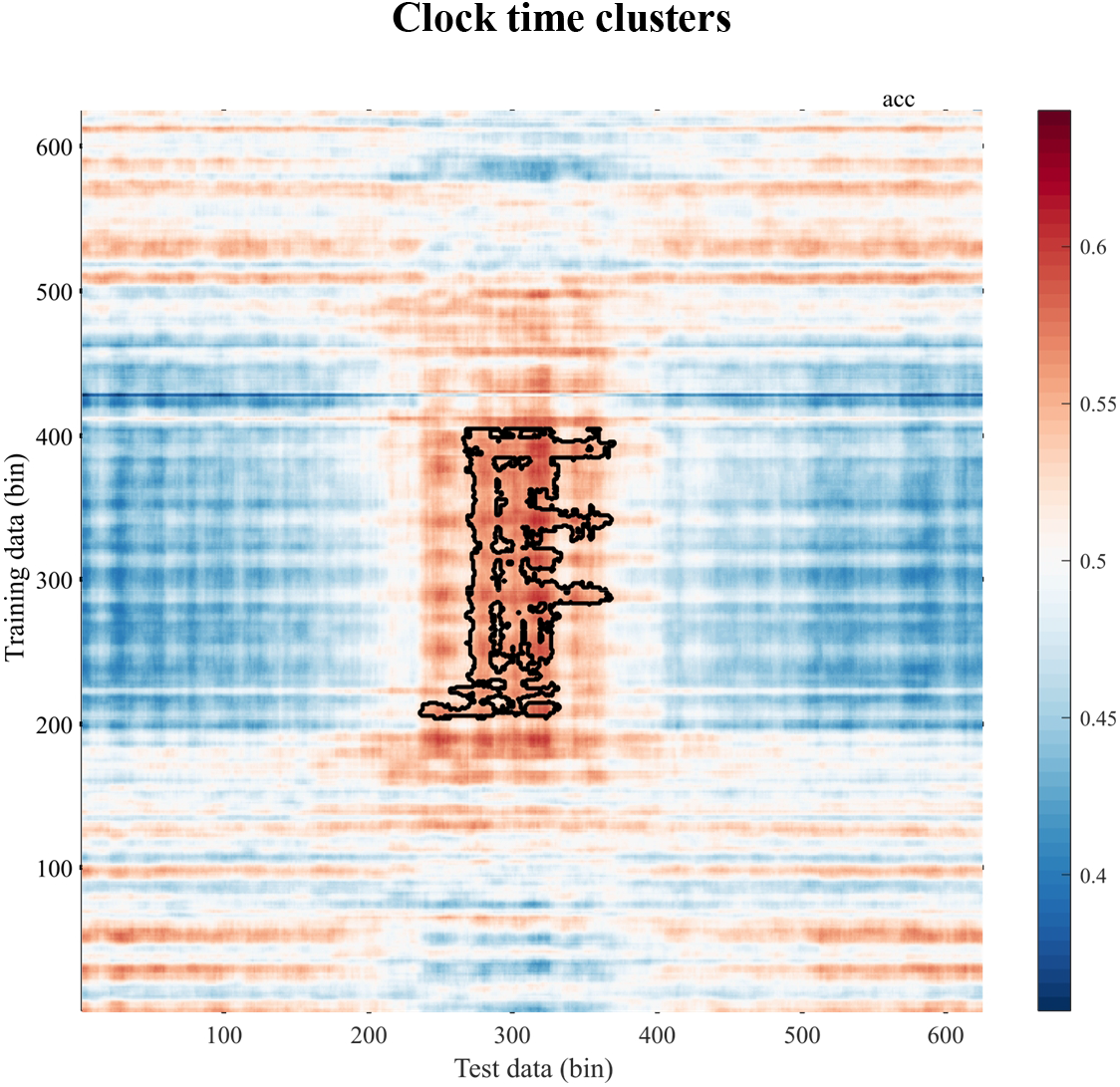

##### 3.3 Human

###### 3.3.1 Basic (main analysis)

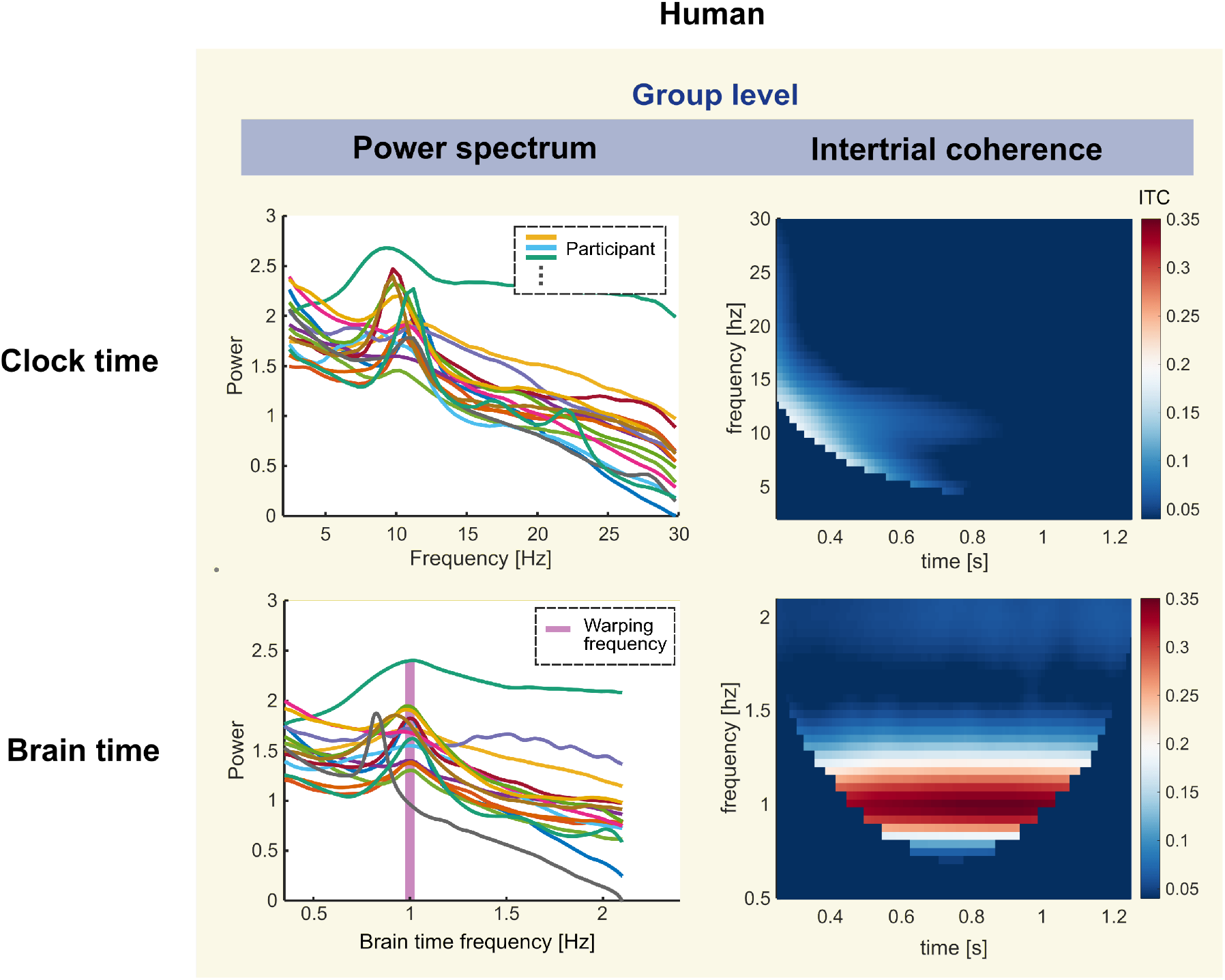

###### 3.3.2 Advanced (main analysis)

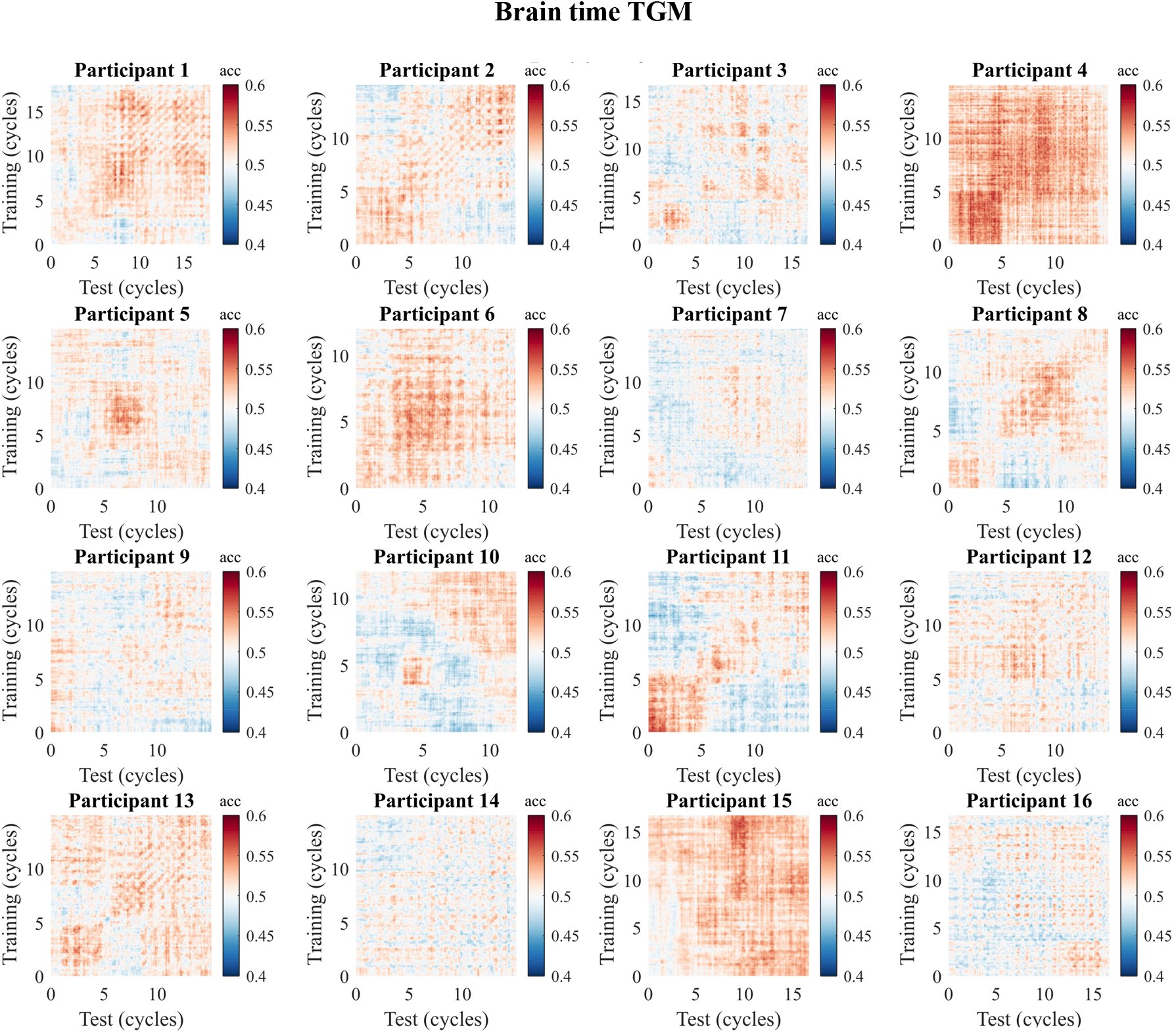

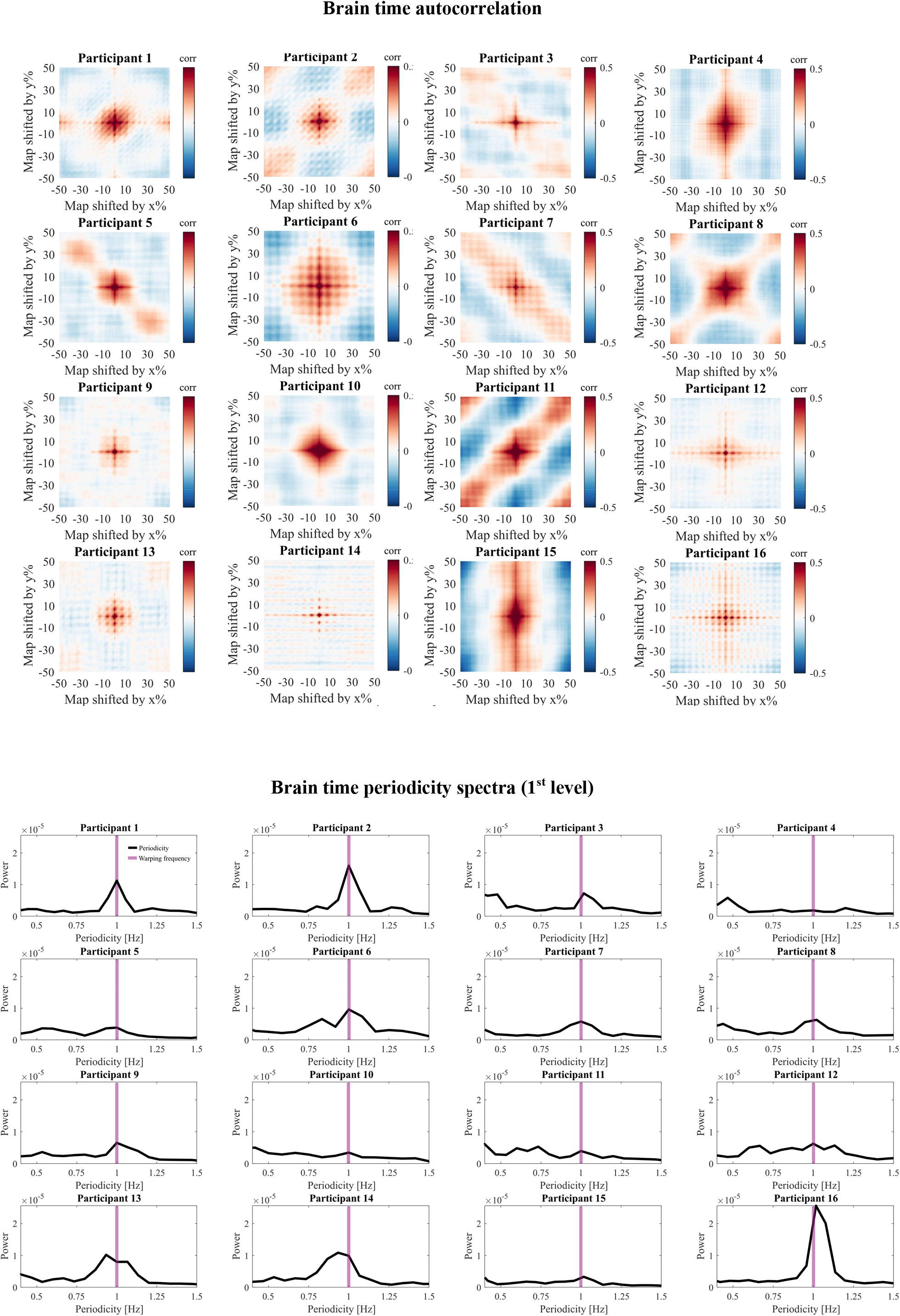

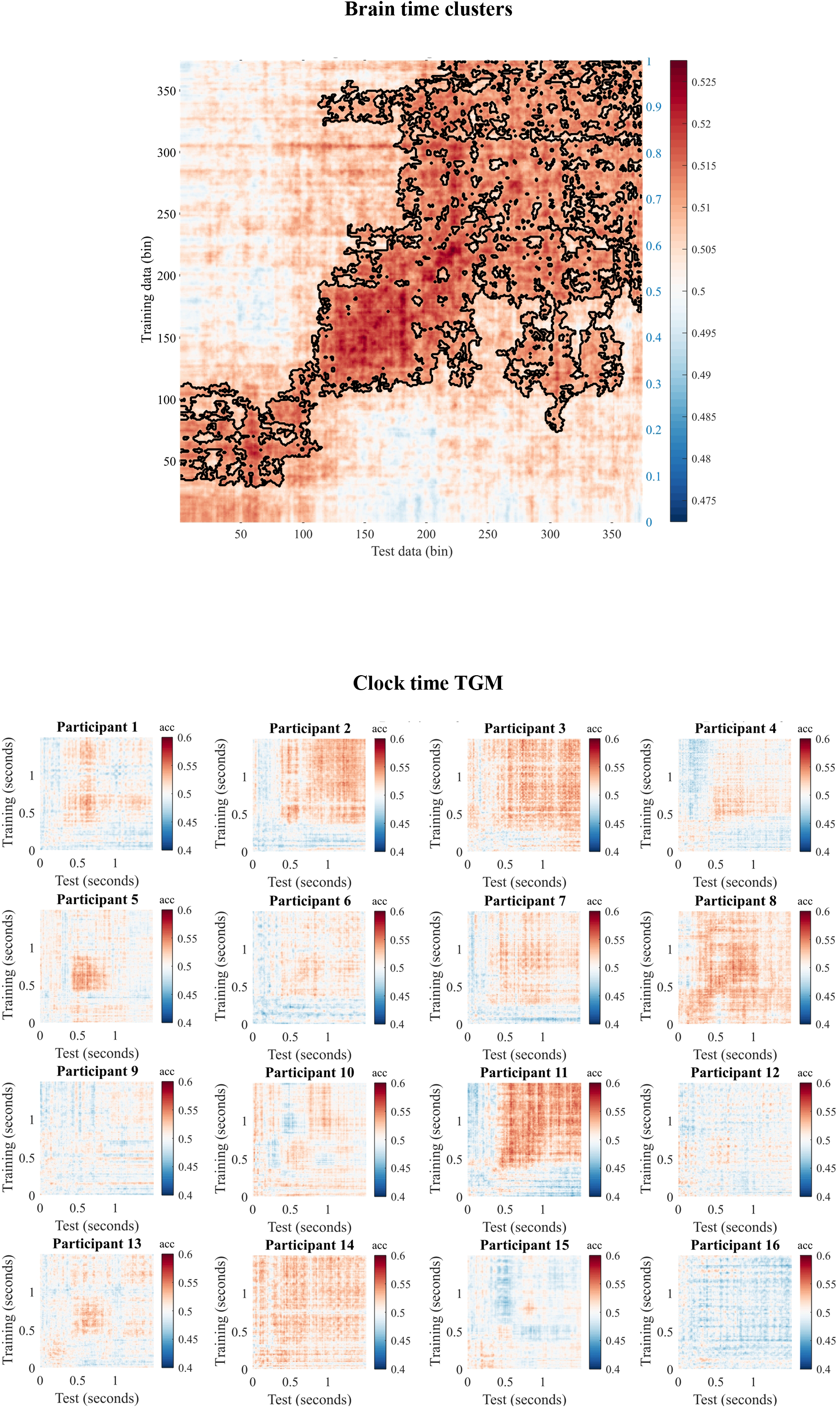

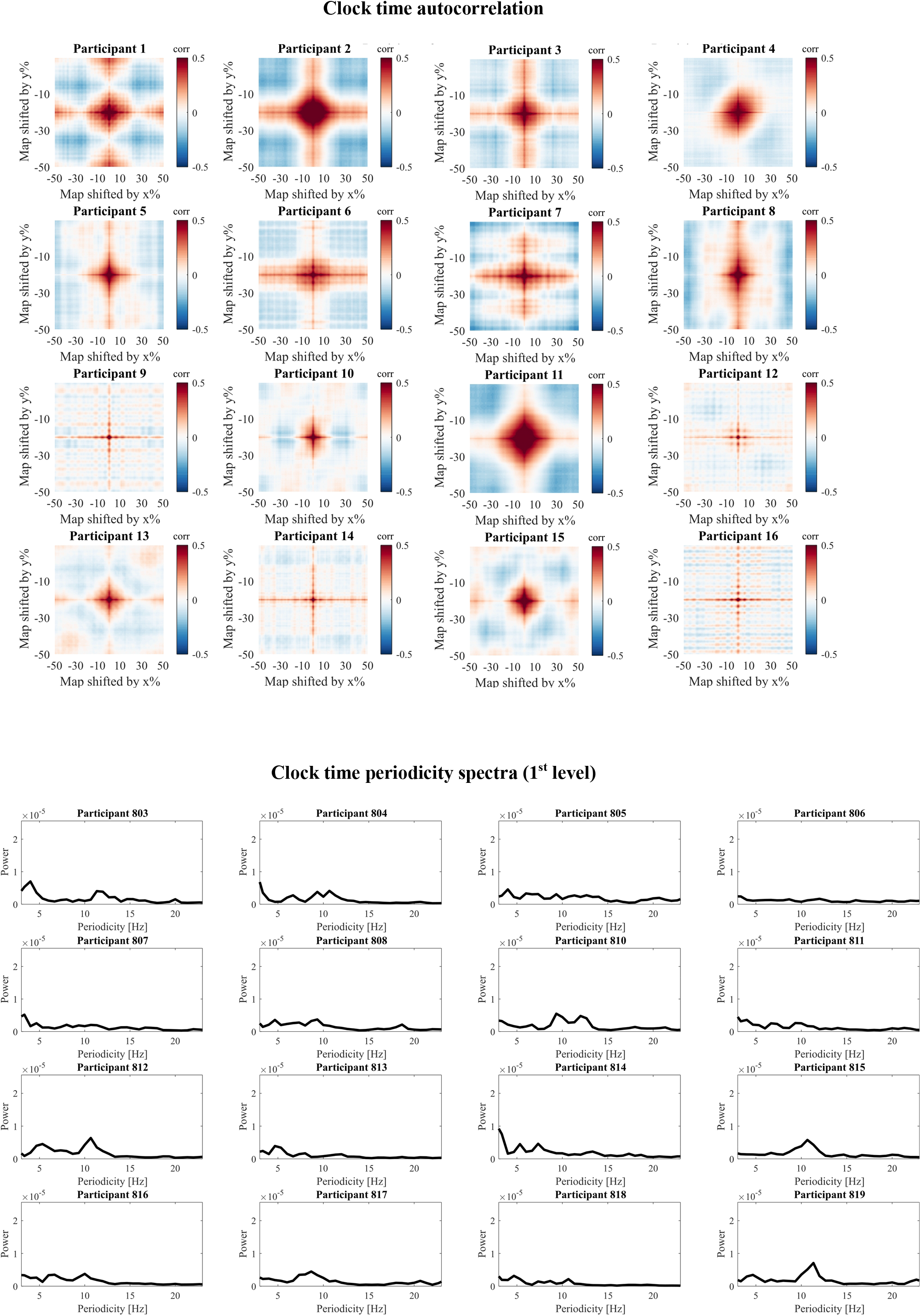

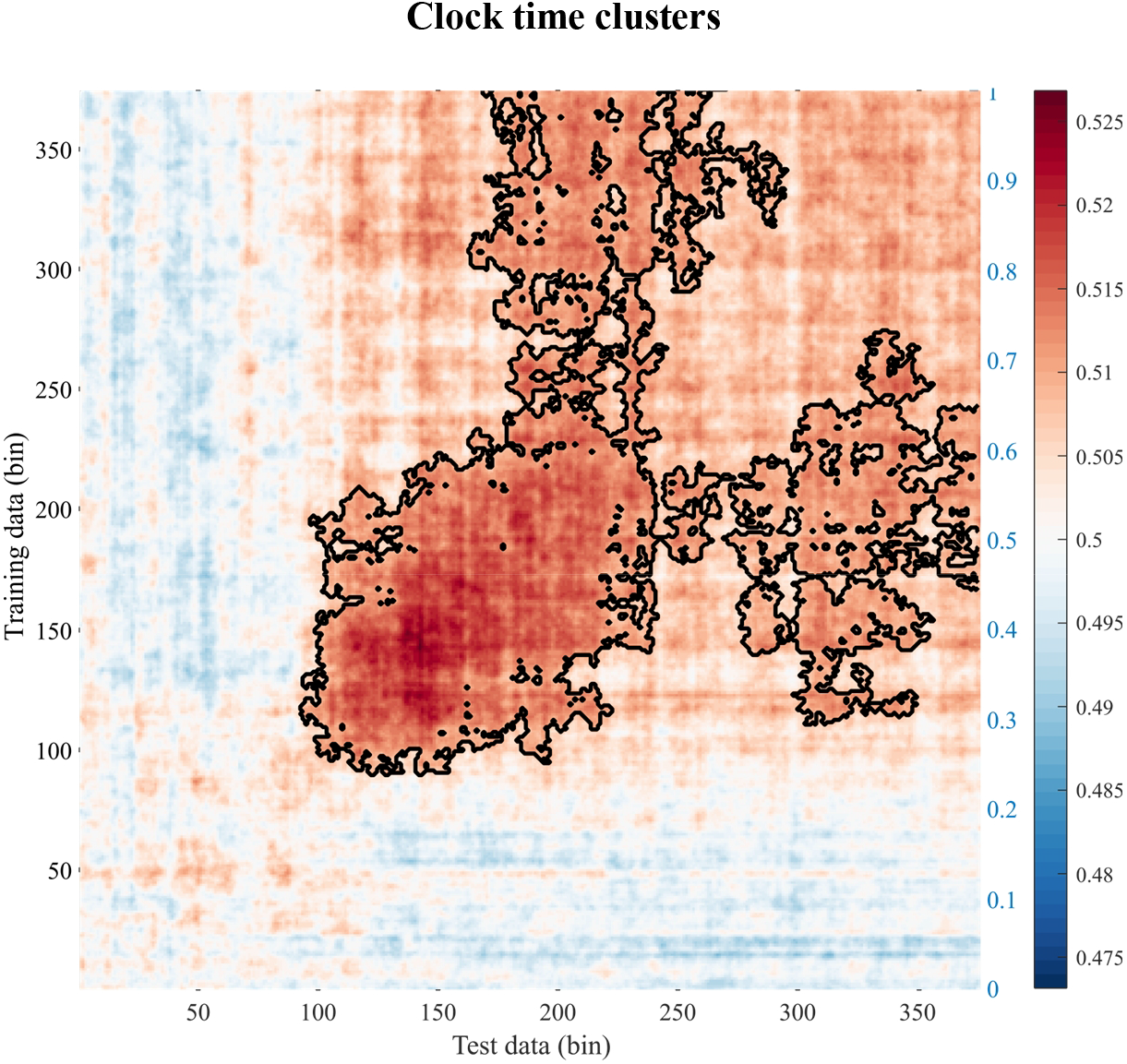

